# Bioenergetic-Related Gene Expression in the Hippocampus Predicts Internalizing vs. Externalizing Behavior in a F_2_ Cross of Selectively-Bred Rats

**DOI:** 10.1101/2022.07.14.500129

**Authors:** Elaine K. Hebda-Bauer, Megan H. Hagenauer, Daniel B. Munro, Peter Blandino, Fan Meng, Keiko Arakawa, Apurva S. Chitre, A. Bilge Ozel, Pejman Mohammadi, Stanley J. Watson, Shelly B. Flagel, Jun Li, Abraham A. Palmer, Huda Akil

## Abstract

Selectively-bred High Responder (bHR) and Low Responder (bLR) rats model the extreme externalizing and internalizing behavior accompanying many psychiatric disorders. To elucidate gene expression underlying these heritable behavioral differences, bHRs and bLRs (generation 37) were used to produce a F_0_-F_1_-F_2_ cross. We measured exploratory locomotion, anxiety-like behavior, and reward cue sensitivity (Pavlovian Conditioned Approach), and performed hippocampal RNA-Seq in male and female F_0_s (n=24) and F_2_s (n=250). Behaviors that diverged during selective breeding remained correlated in F_2_s, implying a shared genetic basis. F_0_ bHR/bLR differential expression was robust, surpassing differences associated with sex, and predicted expression patterns associated with F2 behavior. With bHR-like behavior, gene sets related to growth/proliferation were upregulated, whereas with bLR-like behavior, gene sets related to mitochondrial function, oxidative stress, and microglial activation were upregulated. This differential expression could be successfully predicted based on F0 genotype using cis-expression quantitative trait loci (cis-eQTLs) identified in the F2s. Colocalization of these cis-eQTLs with behavioral Quantitative Trait Loci pinpointed 16 differentially expressed genes that were strong candidates for mediating the influence of genetic variation on behavioral temperament. Our findings implicate hippocampal bioenergetic regulation of oxidative stress, microglial activation, and growth-related processes in shaping behavioral temperament, modulating vulnerability to psychiatric disorders.

## Introduction

Psychiatric disorders can be classified using an internalizing versus externalizing model ^1–3^. Internalizing disorders are characterized by negative emotion, including depression, anxiety, and phobias, whereas externalizing disorders are characterized by behavioral disinhibition, including conduct disorder, antisocial behavior, and impulsivity^3^. These internalizing and externalizing tendencies arise from a complex interplay of genetic and environmental factors, including stress and childhood adversity ^1–3^, and are associated with stable personality or temperament traits that emerge early in development ^4–11^. Neuroticism is strongly associated with internalizing disorders ^12–15^, whereas sensation-seeking is associated with externalizing disorders ^16–20^. Temperament traits are highly heritable, and human genome wide association studies (GWAS) of neuroticism, impulsivity, and externalizing tendencies have demonstrated strong correlations between genetic variation underlying temperament and genetic risk for developing mood and substance use disorders ^10,21–23^. Thus, elucidating the genetic contribution to temperament could provide insight into the etiology of a variety of psychiatric disorders.

To tackle this issue, we produced two rat lines representing temperament extremes by selective breeding for greater than 37 generations for a high and low propensity to explore a novel environment ^24–26^. The bred high responders (bHRs) have high exploratory locomotion and disinhibited, sensation-seeking, externalizing-like behavior. They show greater impulsivity, low anxiety, and an active coping style. They are highly sensitive to reward cues, which can become attractive and reinforcing in their own right within a Pavlovian conditioned approach (PavCA) task (“sign-tracking”^27^). In contrast, bred low responders (bLRs) have low exploratory locomotion and inhibited, internalizing-like behavior. They show elevated anxiety- and depressive-like behavior, stress reactivity, a passive coping style ^24,26,28–30^, and primarily use reward cues for their predictive value (PavCA “goal-tracking”^27^). These behavioral phenotypes appear early in development ^31,32^ similar to temperament in humans ^10,33^. Thus, the highly divergent bHR/bLR phenotypes model heritable, temperament extremes predictive of internalizing and externalizing psychiatric disorders in humans. They can also model two paths to substance use disorders: sensitivity to reward cues and sensation-seeking makes bHRs more likely to initiate and re-initiate substance use, whereas bLRs increase substance use following stress ^24^.

The divergent bHR/bLR behavioral phenotypes make them a powerful model for investigating functional genomics underlying temperament. We have carefully minimized in-breeding via rotational breeding, but following many generations of selective breeding some phenotype differences may still represent artifacts of inbreeding. Therefore, to further elucidate the genetic contribution to temperament, we bred bHRs with bLRs from generation 37 (F_0_) to produce F_1_ cross offspring (Intermediate Responders, “IR”). These F_1_ offspring were then bred with each other to produce a re-emergence of diverse behavioral phenotypes in the F_2_ generation (**Fig 1**). We initially performed exome and whole genome sequencing on the F_0_ and F_2_ rats ^34,35^, which revealed coding differences segregating the bHR/bLR lines (F_0_s) and chromosomal regions associated with variation in exploratory locomotion and other bHR/bLR distinctive behaviors in F_2_ adults and juveniles (quantitative trait loci (QTLs)). Several behaviors were also deemed highly heritable, especially exploratory locomotion. Our goal in the current study was to obtain gene expression data for integrating convergent evidence among this genetic variation, behavioral differences, and brain function underlying behavioral temperament.

**Fig 1.**
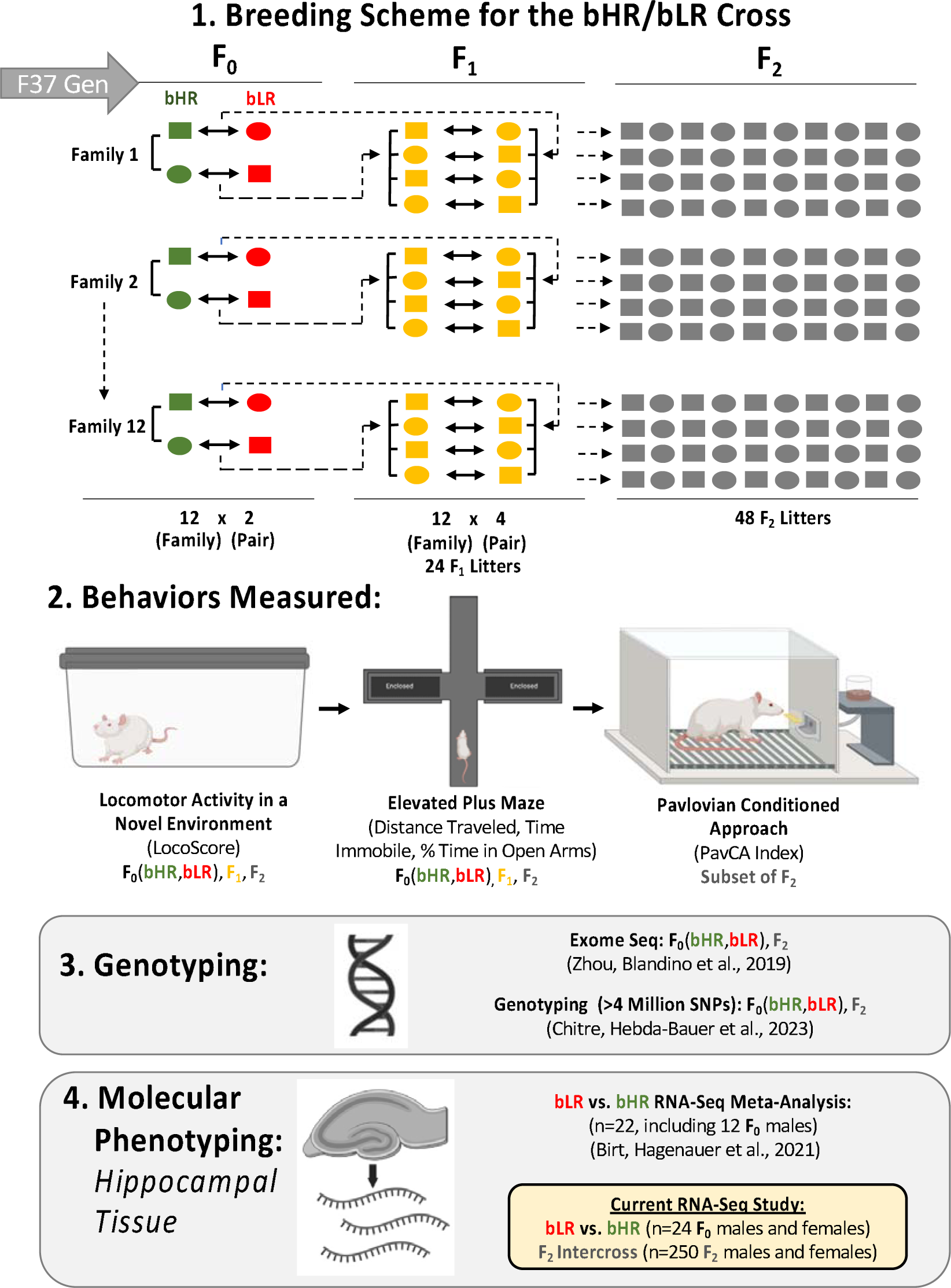
Experimental design: Crossbreeding bHR and bLR rats to identify genes implicated in exploratory, anxiety-like, and reward-related behaviors. After 37 generations of selectively breeding rats for a high or low propensity to explore a novel environment, we have generated two lines of rats (high responders to novelty (bHRs) or low responders to novelty (bLRs)) with highly divergent exploratory locomotion, anxiety-like behavior, and reward-related behavior (Pavlovian conditioned approach (PavCA)). 1. Breeding Scheme: An initial set (F_0_) of bHRs were bred with bLRs to create 12 intercross families. The offspring of this intercross (F_1_) were then bred with each other to produce a re-emergence of diverse phenotypes in the F_2_ generation. 2. Behavior: All rats were assessed for locomotor activity in a novel environment (LocoScore) as well as exploratory and anxiety-like behavior in the elevated plus maze (EPM). For a subset of F_2_ rats, sensitivity to reward-related cues (Pavlovian Conditioned Approach (PavCA) behavior) was also measured. 3. Genotyping: To identify genomic loci associated with bHR/bLR phenotype (F_0_ population segregation) and behavior in F_2_ adults and F_2_ juveniles (quantitative trait loci, QTLs), exome sequencing was initially performed on both F_0_ and F_2_ rats ^35^, followed by a broader genome wide association study (GWAS) ^34^. Within the GWAS, the whole genome was deeply sequenced for the F_0_ rats. This whole genome sequencing (WGS) data was then used in conjunction with low pass (∼0.25x) WGS data from the larger cohort of F_2_ rats to impute the genotype of 4,425,349 single nucleotide variants (SNPs) for each rat ^34^ 4. Molecular Phenotyping: RNA-Seq was used to characterize gene expression in the hippocampus of a subset of male F_0_ and F_1_ rats (n=6/subgroup), which was included in a cross-generational bHR/bLR meta-analysis ^43^. In our current study, RNA-Seq was used to characterize hippocampal gene expression in an independent set of males and females in the F_0_ (n=24, n=6 per phenotype per sex) and F_2_ (n=250) rats to identify gene expression related to both bHR/bLR lineage and exploratory locomotion, anxiety-like behavior, and PavCA behavior.

Gene expression differences via RNA-Sequencing in the hippocampus were identified in both F_0_ and F_2_ animals. The hippocampus was chosen for RNA-Sequencing due to its importance in behavioral inhibition, emotional regulation, environmental reactivity, and stress-related responses ^36–40^ and its previously-identified role in the development and expression of bHR/bLR phenotypes ^31,41,42^. The RNA-Sequencing results from hippocampal tissue of both male and female bHRs and bLRs (F_0_, *n*=24) confirmed our earlier results from a meta-analysis of eight small transcriptional profiling datasets comparing hippocampal gene expression in bHR versus bLR males, collected over 43 generations of selective breeding ^43^. We then performed RNA-Sequencing of hippocampal tissue from a large F_2_ intercross sample (*n*=250) to identify differential expression that continued to correlate with exploratory locomotion, anxiety-like behavior, and reward-related behavior independent of the linkage disequilibrium and genetic drift specific to our bred lines and compared these results to differential expression from other rat models.

To determine which differential expression was most likely driven directly by genetic variation, we integrated our current F_2_ RNA-Seq data with our previous whole genome sequencing results ^34^ to identify genes with expression tightly correlated with proximal genetic variation (expression QTLs: *cis*-eQTLs). We then determined which of these *cis*-eQTLs were segregated in the bHR/bLR lines and located within regions associated with bHR/bLR-like behavior (QTLs) within the larger F_2_ sample (adults and juveniles: ^34^) (**Fig 2**). The convergence of genetic and functional genomic evidence from our bred lines and intercross cohorts was then used to identify a set of intriguing candidates for mediating the influence of selective breeding on behavioral temperament.

**Fig 2.**
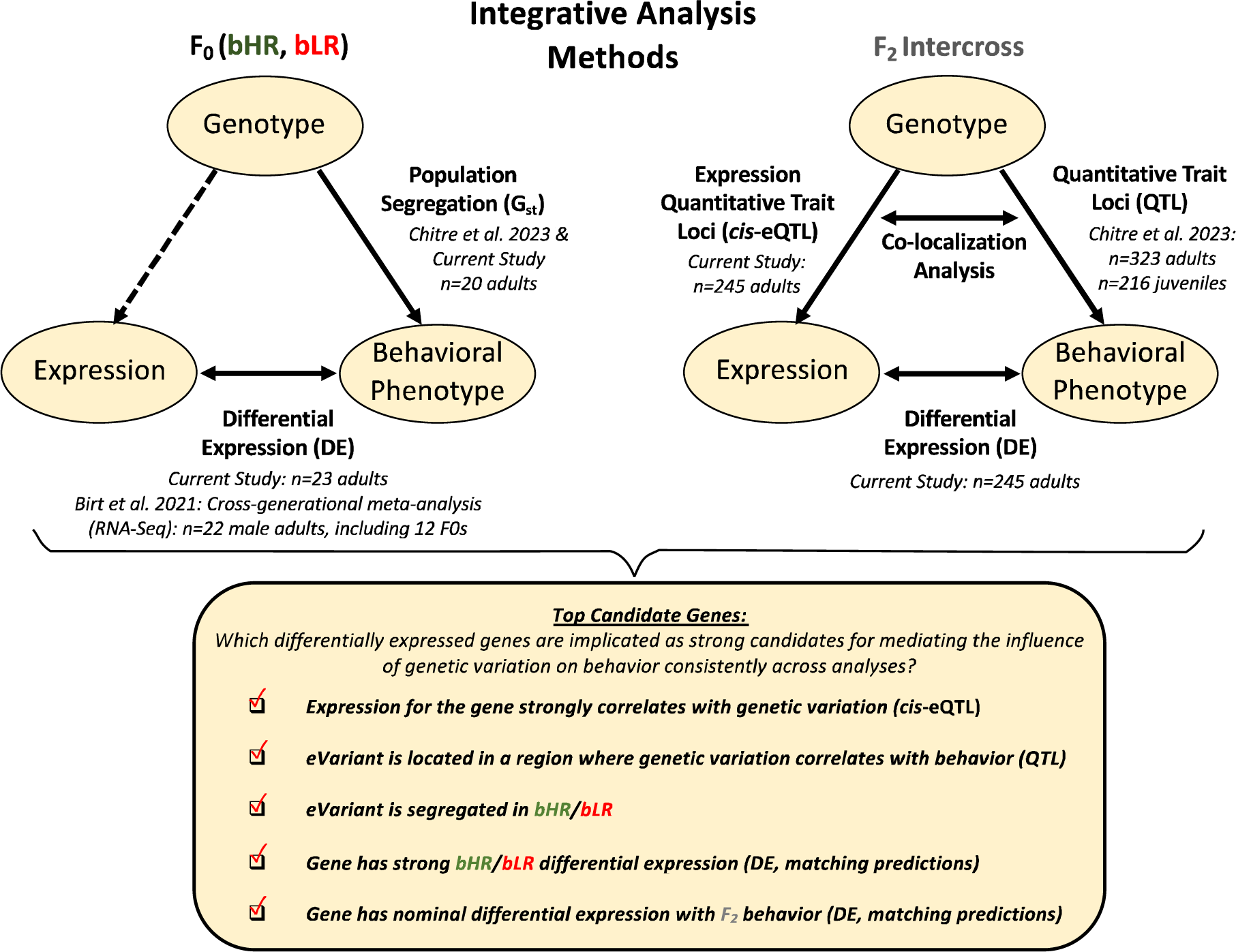
Analysis methods: Using convergent evidence to identify differentially expressed genes that are the strongest candidates for mediating the influence of genetic variation on behavioral temperament. We first identified expression in the hippocampus that differentiated the bHR and bLR lines, using information from both our current F_0_ sample and previous cross-generational meta-analysis. We then identified gene expression that continued to correlate with bHR/bLR divergent behaviors in our large F_2_ intercross sample, indicating that the differential expression (DE) was not an artifact of genetic drift. To determine which DE was most likely driven by proximal genetic variation, we performed a cis-expression Quantitative Trait Loci (cis-eQTL) analysis to determine which genes (eGenes) had expression that strongly correlated with nearby genetic variants (eVariants) using the F_2_ genotype and F_2_ gene expression data, and then estimated the magnitude of that effect in Log2 fold change units (allelic fold change or aFC). We determined which eVariants were segregated in the bHR/bLR lines and confirmed that the predicted effect of this genetic segregation on gene expression matched the bHR/bLR DE observed in the F_0_s. We also determined which eVariants co-localized in regions of the genome previously identified as having strong relationships with behavior in the full sample of F_2_ adults and F_2_ juveniles QTLs) using Summary-Data Based Mendelian Randomization (SMR) and compared the predicted direction of effect of this genetic variation on gene expression to the DE observed in the F_2_ adults in association with behavior. Finally, to be considered a top candidate gene for mediating the influence of genetic variation on behavioral temperament, we required that a gene have hippocampal DE related to both bHR/bLR phenotype and F_2_ behavior which appeared plausibly driven by genetic variation (cis-eQTL) that segregated in the bHR/bLR lines and correlated with behavior (SMR colocalization with QTL). Note that the sample sizes listed in the diagram reflect the sample sizes used in the analysis following quality control.

## Results

### Locomotor activity in a novel environment reflects a broader behavioral temperament in both selectively-bred bHR/bLR lines and F_2_ intercross rats

The bHR/bLR (F_0_) crossbreeding scheme produced F_2_ animals with behaviors ranging between the more extreme bHR and bLR phenotypes (all behaviors: *p*<1.5e-06 for effect of group (F_0_ bHR vs. F_0_ bLR vs. F_2_); examples: **Fig 3A-B**; **Fig S4**, full statistics: **Table S1**). F_2_ behavior sometimes appeared more similar to bLRs than bHRs (*e.g.,* **Fig 3B**) suggesting a floor effect or that genetic contributions to internalizing-like behavior may be more dominant.

**Fig 3.**
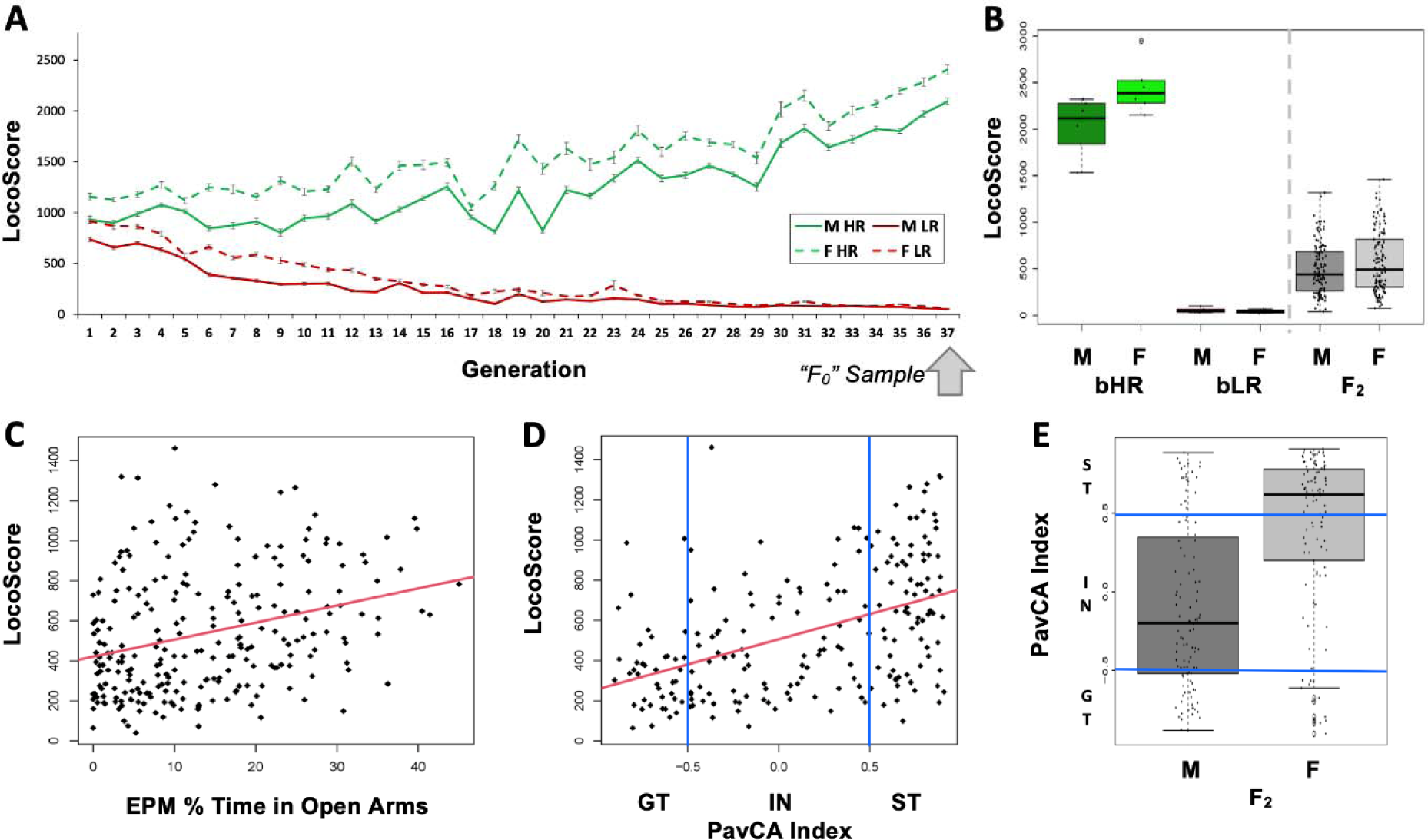
Behavioral phenotype: Locomotor activity in a novel environment reflects a broader behavioral temperament in both selectively-bred bHR/bLR lines and F_2_ intercross rats. **A)** Locomotion in a novel environment (LocoScore: lateral + rearing counts over 60 minutes) is strongly influenced by genetics, as indicated by the divergence in LocoScore observed following the selective breeding of the bHR/bLR lines in both males (M) and females (F). The “F_0_” animals used in our crossbreeding experiment were taken from the 37^th^ generation of selective breeding. **B)** The bHR/bLR (F_0_) crossbreeding scheme produced F_2_ rats with intermediate behavior which fell between the extreme bHR and bLR phenotypes. Depicted are the LocoScores for the F_0_ and F_2_ rats included in the current RNA-Seq experiment (effect of group: F(2, 268)=212.764, p<2.2E-16). The graphs for other measured behaviors (EPM Distance traveled, EPM time immobile, EPM % Time in Open Arms) are in **Fig S4**. For all measured behaviors, group differences (ANOVA: F_0_ bHR vs. F_0_ bLR vs. F_2_) were highly significant (p<1.5e-06, **Table S1**). **C-D)** Behaviors that diverged during bHR/bLR selective breeding for LocoScore remained correlated in F_2_s. Example scatterplots illustrate correlations between LocoScore and other bHR/bLR distinctive behaviors in the F_2_s. Red lines illustrate the relationship between the variables across both sexes. The correlations between all variables can be found in **Fig S4** & **Table S2**. **C)** Greater LocoScore predicted greater EPM % Time in Open Arms in the F_2_s (R=0.30, p=1.72E-06). Greater time spent in the open arms of the EPM is typically interpreted as indicating low anxiety. **D)** Most F_2_s (n=209) were tested for PavCA behavior. Greater LocoScore predicted an elevated PavCA Index in the F_2_s (R=0.46, p=3.22E-12). Rats with greater PavCA Index (>0.5) are considered sign-trackers (ST) and rats with lower PavCA Index (<-0.5) are considered goal-trackers (GT). **E)** All behavioral variables showed a significant sex difference (p<0.007) except for LocoScore. As an example, sex differences in PavCA index are illustrated with a boxplot. A higher percent of females than males were classified as Sign Trackers (ST) vs. Goal Trackers (GT) (Intermediate=IN) (Fisher’s exact test: p=1.472E-06, OR: 0.13; CI: 0.05-0.34). These observed behavioral sex differences are difficult to interpret, as the two sexes were tested on all tasks in separate batches but supported the inclusion of sex as a covariate in all statistical models.

Exploratory locomotion, anxiety-like behavior, and PavCA behavior remained strongly correlated within the F_2_s, as previously observed in the bHR/bLR lines (examples: **Fig 3C-D**; **Fig S4**, full statistics: **Table S2**). Although these behaviors often differed by sex (example: **Fig 3E, Fig S4**, full statistics: **Table S1**), sex differences were not responsible for driving the correlation between different behaviors (with sex in the model: all behavior-behavior relationships still *p*<0.0284). These findings are consistent with results using the full F_2_ cohort ^34^ and imply that locomotor activity in a novel environment echoes a broader behavioral temperament, reflecting genetic and environmental influences shared across anxiety, mood, and reward-related behaviors.

### F_0_ RNA-Seq: Selective breeding produced a robust molecular phenotype

Within the F_0_ RNA-Seq dataset, there were 131 differentially expressed genes with elevated hippocampal expression in bLRs versus bHRs, and 86 differentially expressed genes with higher expression in bHRs (False Detection Rate (FDR)<0.10, **Fig 4A&C, Table S3**). In contrast, there were only 21 genes upregulated in females (versus males) and 22 genes upregulated in males (versus females) (**Fig 4B-C, Table S3)**. The effect sizes (Log(2) Fold Changes, or Log2FC) for bHR/bLR differentially expressed genes were also larger than those for sex, with the exception of a few X and Y chromosome genes (**Fig 4A-B, Table S3)**. There were no significant interactions between the effects of Lineage and Sex on gene expression (FDR>0.10), but our sample size was underpowered to detect these effects (n=5-6/subgroup). The presence of robust bHR/bLR hippocampal differential expression replicated previous studies ^43^.

**Fig 4.**
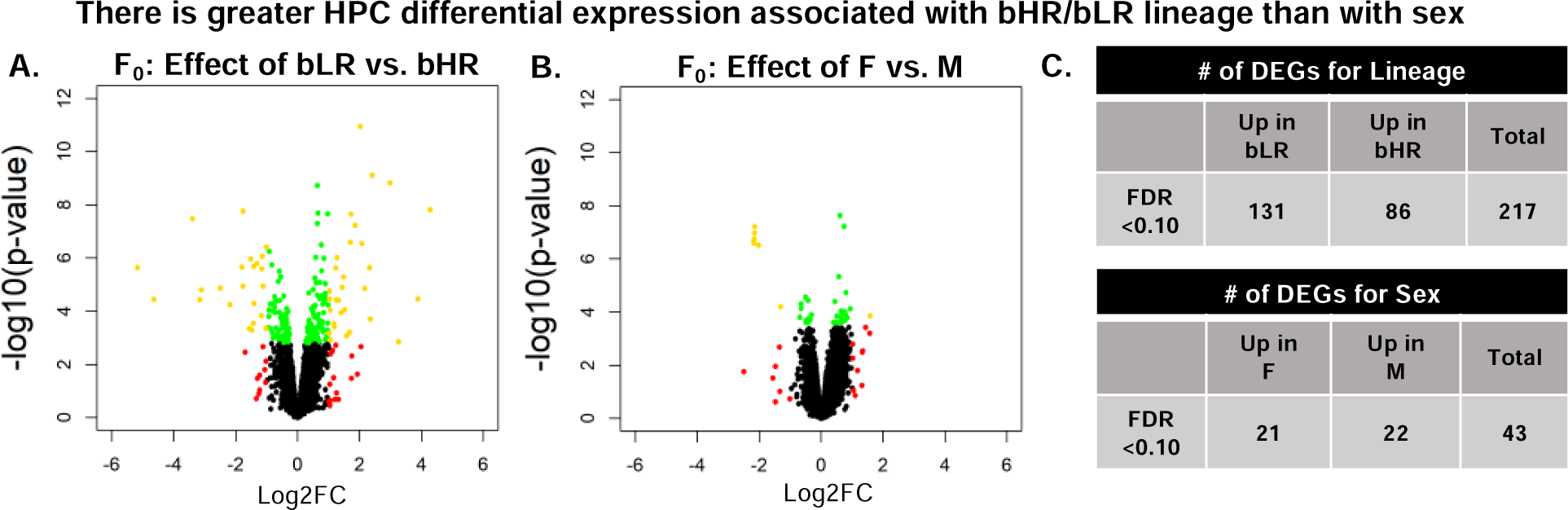
A robust hippocampal (HPC) molecular phenotype: There was greater hippocampal differential expression associated with F_0_ bHR/bLR lineage than with sex. Differential expression associated with bHR/bLR Lineage and Sex were examined in the same dataset (F_0_) with comparable subgroup sample sizes (total n=23). Shown are two volcano plots illustrating the differential expression associated with bHR/bLR phenotype (**A**) and sex (**B**). For both volcano plots, red depicts genes with a log2 fold change (Log2FC)>1.0, green depicts genes with a False Discovery Rate (FDR)<0.1, and gold indicates genes satisfying both criteria. In (**A**), the reference group was defined as bHR, therefore positive Log2FC coefficients indicate upregulation in bLRs, and negative Log2FC coefficients indicate upregulation in bHRs. In (**B**), males served as the reference group, therefore positive Log2FC coefficients indicate upregulation in females (F), and negative Log2FC coefficients indicate upregulation in males (M). For ease of visualization, six X and Y chromosome genes weren’t plotted due to extreme p-values (ranging from p=5.71E-13 to 8.97E-26: Kdm5d, Eif2s3y, Uty, Ddx3, ENSRNOG00000055225, AABR07039356.2). The summary table (**C**) shows the number of differentially expressed genes (DEGs) for bHR/bLR Lineage and Sex.

### Current F_0_ study replicated bHR/bLR gene expression differences detected in previous studies

The bHR/bLR hippocampal differential expression in our current study replicated many effects observed in our previous meta-analysis of bHR/bLR hippocampal transcriptional profiling studies ^43^, with the F_0_ Log2FC correlating positively with the bLR versus bHR estimated effect size (d) observed in our late generation RNA-Seq meta-analysis ^43^ (**Fig 5A-B**). Sixty-two of the 984 bHR/bLR differentially expressed genes in either dataset (FDR<0.10) were significant (FDR<0.10) in both datasets (**Fig 5C, Table S3**). More genes showed replication of nominal bHR/bLR effects (p<0.05) with consistent direction of effect in both datasets, so that, in total, 1,063 genes had evidence of bHR/bLR differential expression (**Fig 5D-E, Table S3)**. As many generations of selective breeding for a behavioral phenotype are likely to produce an enrichment of eQTL alleles influencing the phenotype, these 1,063 bHR/bLR differentially expressed genes were prioritized in downstream analyses.

**Fig 5.**
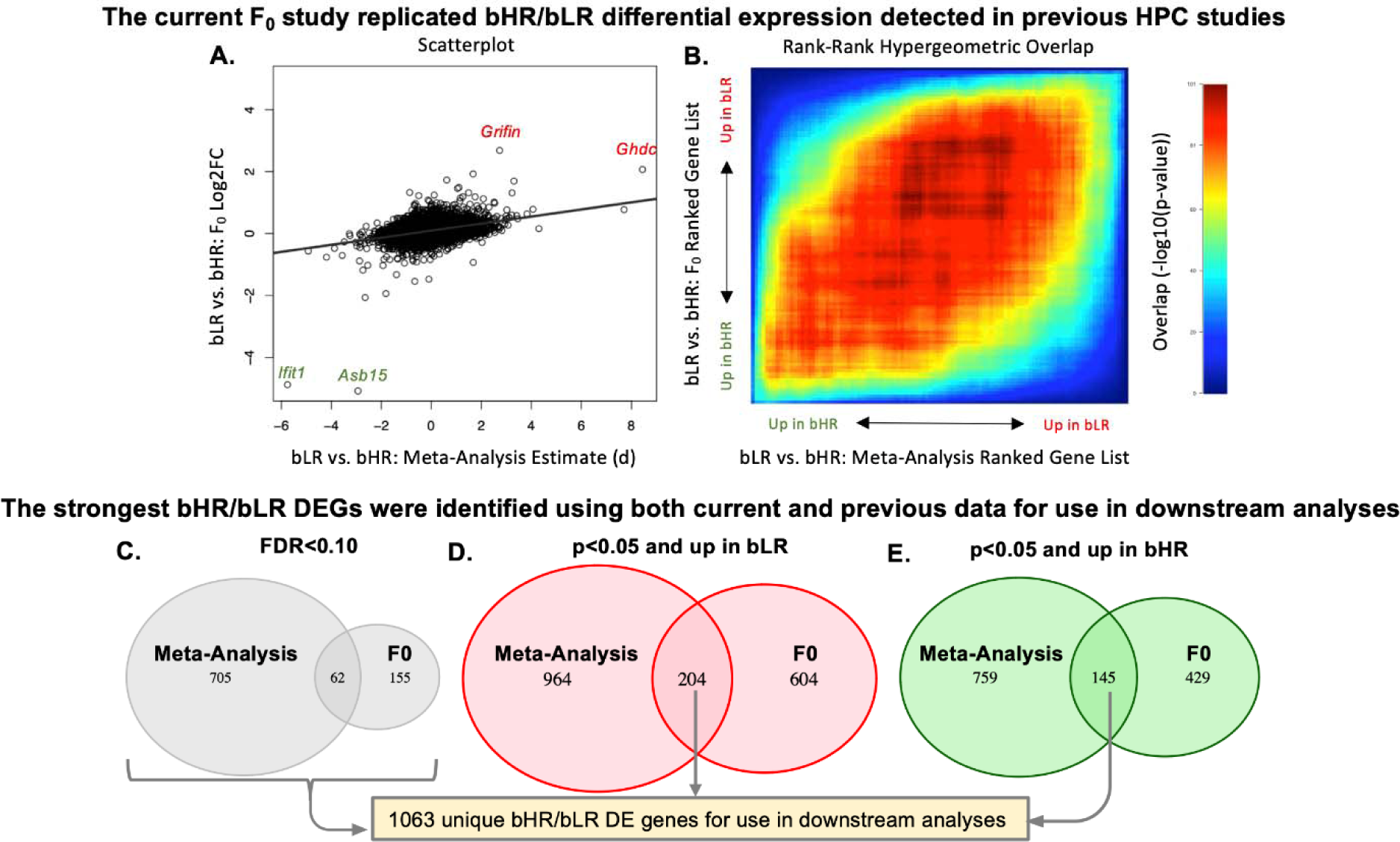
Validation: The current F_0_ study replicated bHR/bLR differential expression (DE) detected in previous hippocampal (HPC) studies. **A**) A scatterplot illustrates the positive correlation (n=11,175 genes, R=0.35, p<2.2E-16) between the bHR/bLR effect sizes from our current F_0_ dataset (bLR vs. bHR Log2FC for all genes) and the bLR vs. bHR effect sizes identified by our previous late generation RNA-Seq meta-analysis (^43^: bLR vs. bHR estimated cohen’s d for all genes). Genes with particularly large effects in both datasets are labeled in green (upregulated in bHRs) or red (upregulated in bLRs). **B**) This positive correlation is also visible when using a non-parametric analysis of the results ranked by t-statistic, as illustrated with a two-sided Rank-Rank Hypergeometric Overlap (RRHO) plot. Warmer colors illustrate the strength of the overlap (-log10(p-value)), with the visible red diagonal indicating a positive correlation between the ranked results. **C-E)** To identify the strongest bHR/bLR differentially expressed genes (DEGs) for use in downstream analyses, we referenced results from both the current F_0_ dataset and previous late generation RNA-Seq meta-analysis ^43^. Shown are three Venn diagrams illustrating the overlap of the DEG lists from the two studies, with DE defined either using a traditional threshold of FDR<0.10 in either study (**C**) or using a nominal p-value threshold (p<0.05) and a specified direction of effect (**D**: upregulated in bLR vs. bHR, **E**: upregulated in bHR vs. bLR). In each case, the overlap exceeded what would be expected due to random chance (OR>4.7, p<2.2e-16). The 1,063 unique genes with either FDR<0.10 in either study or nominal replication with a consistent direction of effect in both studies were considered to have the strongest evidence of bLR vs. bHR DE and highlighted in downstream analyses.

### F_0_ DE predicts differential expression related to F_2_ behavior

Since some differential expression may be due to either linkage disequilibrium with causal variants or genetic drift specific to our bred lines, we performed RNA-Seq on hippocampal tissue from a large F_2_ intercross sample (n=250) to identify differential expression that continued to independently correlate with exploratory locomotion, anxiety-like behavior, and reward-related behavior. Hippocampal gene expression associated with bLR lineage resembled the expression associated with lower F_2_ LocoScore, as indicated by the negative correlation between the F_0_ bLR vs. bHR Log2FCs for all genes and the F_2_ LocoScore Log2FCs for all genes (R=-0.20, p<2.2e-16, **Fig 6A, Table S4**). Similarly, gene expression associated with bLR lineage partially resembled expression in F_2_s exhibiting lower exploration (distance traveled) and greater anxiety (greater time immobile, less time in the open arms) on the elevated plus maze (EPM) task (**Table S4**), and greater goal-tracking behavior on the PavCA task (lower PavCA Index; **Table S4**). This pattern of correlations confirmed that a portion of the hippocampal differential expression that emerged following selective breeding was related to behavioral temperament.

**Fig 6.**
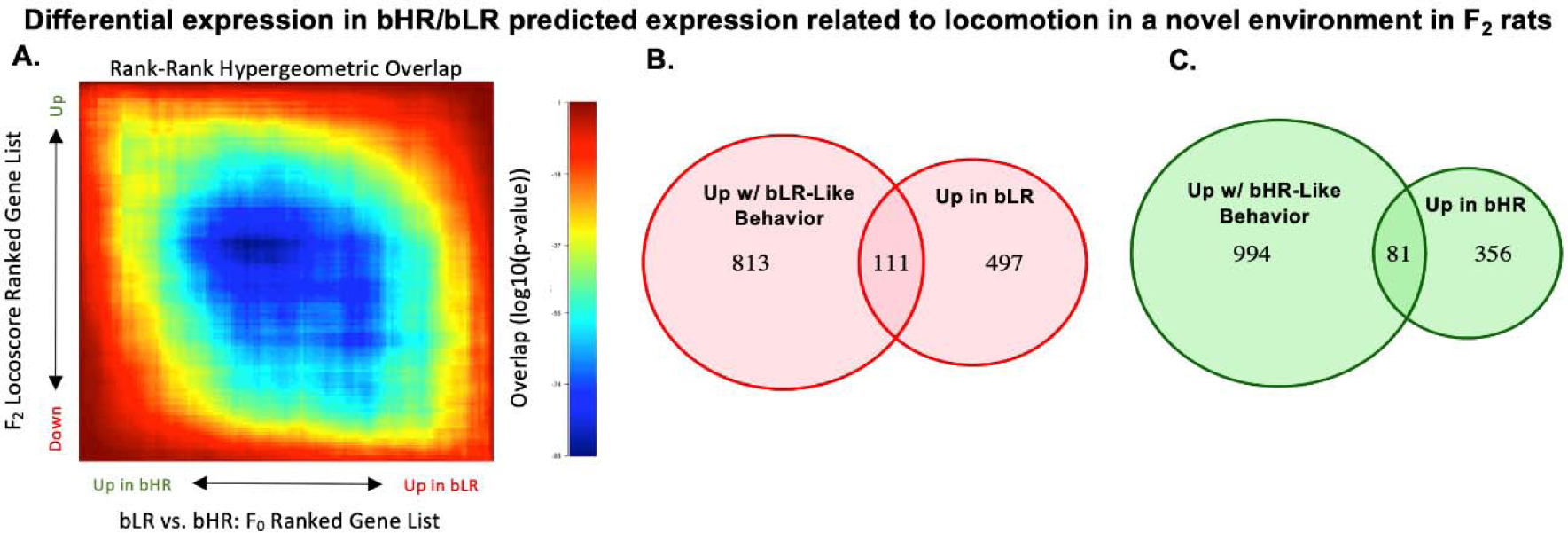
Hippocampal gene expression in bLR vs. bHR F_0_ rats predicts the pattern of gene expression associated with bLR-like vs. bHR-like behavior in F_2_ intercross rats. **A)** For example, there is a negative correlation between the Log2FC associated with locomotion in a novel environment (LocoScore) in the F_2_s and the Log2FC for bLR vs. bHR lineage in the F_0_s (n=13,339 genes, R=-0.196, p<2e-16), which matches the prediction that a bLR-like pattern of gene expression resembles the expression associated with lower exploratory activity. Following the plotting conventions from **Fig 5B**, this negative correlation is illustrated using a two-sided RRHO plot. Within the RRHO, the results are ranked by t-statistic. The visible blue diagonal indicates a negative correlation between the ranked results. The correlation between bLR vs. bHR differential expression and gene expression associated with the other F_2_ behaviors can be found in the supplement (**Table S4**). **K**) A pink Venn Diagram illustrates the enrichment of bLR-upregulated differentially expressed genes for nominal (p<0.05) associations with bLR-like behavior in the F_2_s (i.e., gene expression correlated with decreased locomotor activity, decreased distance traveled, increased immobility, decreased % time in the open arms of the EPM, or decreased PavCA Index) (enrichment: Fisher’s exact test: OR: 3.27, p<2.2e-16). **L)** A green Venn Diagram illustrates the enrichment of bHR-upregulated differentially expressed genes for nominal (p<0.05) associations with bHR-like behavior in the F_2_s (i.e., gene expression correlated with increased locomotor activity, increased distance traveled, decreased immobility, increased % time in the open arms of the EPM, or increased PavCA Index) (enrichment: Fisher’s exact test: OR 2.72, p=7.14e-13). We prioritized the 192 genes that satisfied both criteria (i.e., the intersection of the Venn Diagrams) in downstream analyses as differential expression that might mediate the effect of selective breeding on behavior (111 genes upregulated in both bLRs and with bLR-like behavior, 81 genes downregulated in both bHRs and with bHR-like behavior).

These correlations strengthened when focusing specifically on bHR/bLR differentially expressed genes (1,063 genes in **Fig 5C-E**), of which 1,045 were present in the F_2_ dataset (**Table S4)**. Of these genes, 111 showed both upregulation in bLR rats and at least one nominal (p<0.05) association with bLR-like behavior in the F_2_s (*i.e.,* expression correlated with decreased locomotor activity, decreased EPM distance traveled, increased EPM time immobile, decreased EPM % time in open arms, or decreased PavCA Index), a 3.27X enrichment beyond random chance (Fisher’s exact test: 95%CI: 2.61-4.08, p<2.2e-16, **Fig 6B**), and 81 genes showed both upregulation in bHR rats and at least one nominal (p<0.05) association with bHR-like expression in the F_2_s, a 2.72X enrichment beyond random chance (Fisher’s exact test: 95%CI: 2.10-3.51, p=7.14e-13, **Fig 6C**). However, we were unable to identify differentially expressed genes for F_2_ behavior that had strong enough effects to survive false discovery rate correction (FDR<0.10). Therefore, the differential expression with the strongest converging evidence supporting its potential to mediate behavioral temperament (the 111 genes that were upregulated in bLRs and nominally with bLR-like behavior in the F_2_s and 81 genes upregulated in bHRs and nominally with bHR-like behavior in the F_2_s) were used for downstream analyses.

### Multiple genes have hippocampal differential expression consistently associated with behavioral temperament in other rat models as well as in our F_0_ and F_2_ studies

To determine generalizability, we compared our list of differentially expressed genes implicated in behavioral temperament by converging evidence from the bred lines and the F_2_s (identified in **Fig 6B&C**) to a database of 2,581 genes previously identified as differentially expressed in the hippocampus of other bLR-like and bHR-like rat models targeting hereditary behavioral traits resembling extremes on the internalizing/externalizing spectrum (database from ^43^, summarized in **Fig 7A**; ^44–52^). Sixteen of 111 genes that were upregulated in bLRs and with bLR-like behavior in our study were also upregulated in other bLR-like models (**Fig 7B**, enrichment OR: 2.50 (95%CI: 1.37-4.29), Fisher’s exact test: *p*=0.00242) and 14/81 genes that were upregulated in bHRs and with bHR-like behavior in our study were downregulated in other bLR-like models (**Fig 7C**, enrichment OR: 2.07 (95%CI: 1.07-3.74), p=0.0189). Notably, *Tmem144* had elevated hippocampal expression in three other bLR-like rat models (**Fig 7D**, ^44–46^). Five other genes were differentially expressed in two other rat models (**Fig 7D**, *Bphl, Ist1, RGD1359508, Nqo2, Fcrl2*). We expect less than one gene (0.39) in our dataset to show this degree of convergence due to random chance (***Supp. Methods***).

**Fig 7.**
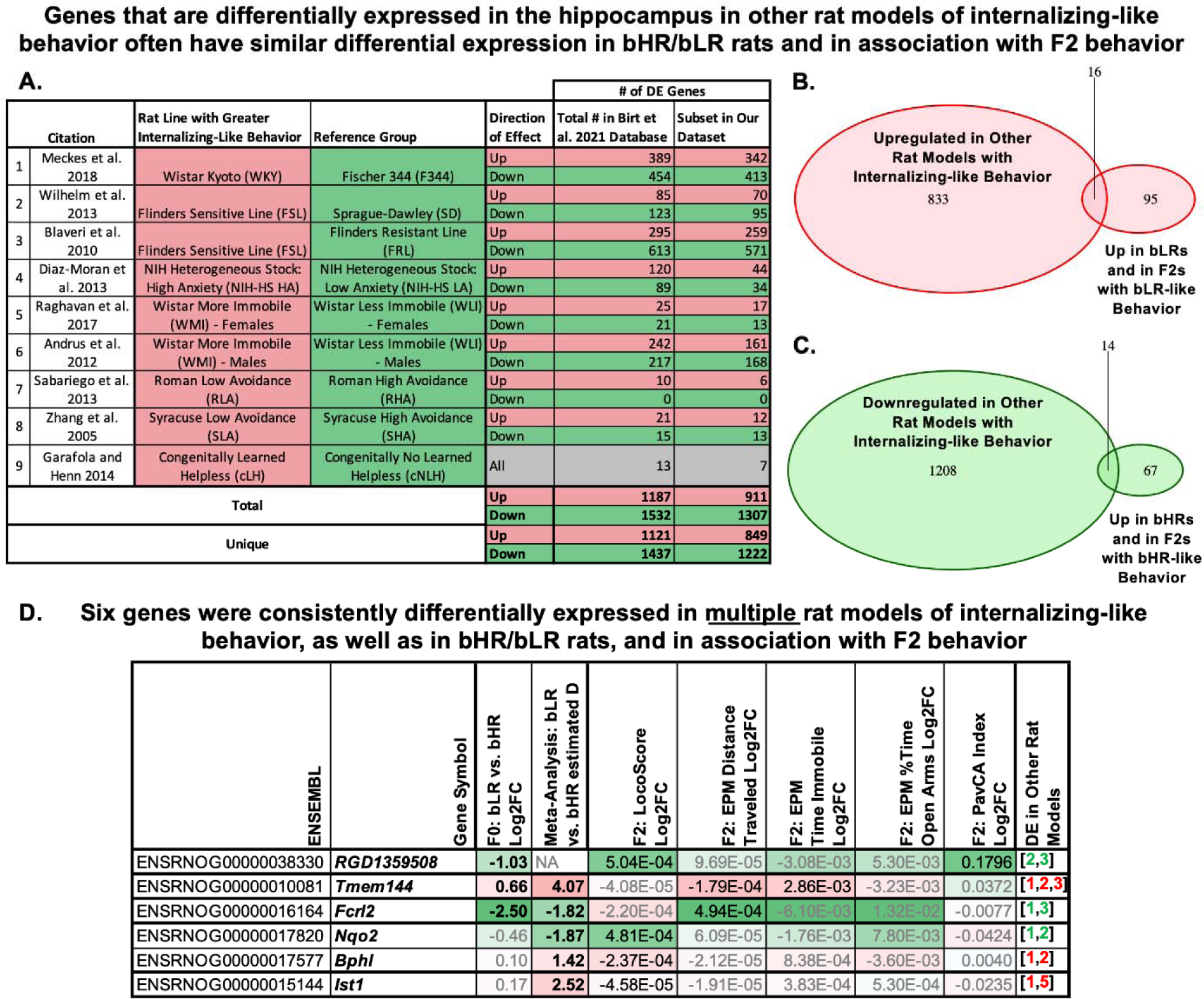
Multiple genes have hippocampal differential expression (DE) consistently associated with hereditary behavioral temperament in other rat models as well as in our F_0_ and F_2_ studies. **A)** To perform this analysis, we compared our current results to a database of 2,581 genes that had been previously identified as differentially expressed in the hippocampus of other bLR-like and bHR-like rat models targeting hereditary behavioral traits resembling extremes on the internalizing/externalizing spectrum (database compiled in ^43^, results from: ^44–52^. The table lists the rat models characterized in the referenced publications, the number of genes up-regulated or down-regulated in association with the rat model exhibiting more internalizing-like behavior (in total, as well as the subset represented in our current datasets). **B**) A pink Venn Diagram illustrates the enrichment of overlap between genes identified as upregulated in other rat models with internalizing-like behavior and the 111 genes that were both upregulated in bLR’s and nominally upregulated in F_2_s with bLR-like behavior in our study (16/111, enrichment OR: 2.50, Fisher’s exact test: p=0.00242). **C)** A green Venn Diagram illustrates the enrichment of overlap between genes identified as down-regulated in other rat models with internalizing-like behavior and the 81 genes that were both upregulated in bHRs and nominally upregulated in F_2_s with bHR-like behavior in our study (14/81, enrichment OR: 2.07, p=0.0189). **D)** A table overviewing the differential expression results for the six genes that were consistently differentially expressed in <u>multiple</u> rat models with internalizing-like behavior, as well as in bHR/bLR rats and in nominal association with F_2_ behavior. Within the table, genes with elevated hippocampal expression in bLRs and in association with bLR-like behavior in the F_2_s are highlighted pink, genes with elevated hippocampal expression in bHRs and in association with bHR-like behavior in the F_2_s are highlighted green. The table includes the Log2FC for bLR vs. bHR Lineage in the F_0_ dataset, the estimated effect size (d) from the late generation bLR vs. bHR RNA-Seq meta-analysis (from ^43^), and the Log2FC for LocoScore, EPM Time Immobile, EPM Distance traveled, EPM % Time in Open Arms, and PavCA index in the F_2_ dataset (Bold=FDR<0.1; black=p<0.05). Note that the F_2_ differential expression analysis includes behavior as a continuous predictor variable, therefore the Log2FC units are defined per unit of LocoScore and not directly comparable to the Log2FC units for bred line (bLR vs. bHR). The final column provides references for similar hippocampal differential expression in other bLR-like (red) or bHR-like (green) rat models following the numbering in the table in panel **A**. Full gene names (when applicable): Tmem144: Transmembrane Protein 144; Fcrl2: Fc Receptor-like 2; Nqo2: N-ribosyldihydronicotinamide:quinone dehydrogenase 2; Bphl: biphenyl hydrolase like; IST1: Factor Associated With ESCRT-III.

### Behavioral temperament is associated with genes involved in growth and proliferation, mitochondrial function, oxidative stress, and microglia

To ascribe functional trends to the differential expression associated with behavioral temperament, we performed Gene Set Enrichment Analysis using a combined score for each gene summarizing bLR-like vs. bHR-like expression across the bHR/bLR and F_2_ analyses. Sixty-three gene sets were upregulated (FDR<0.05 with a bHR-like phenotype (*i.e.,* in bHRs and with bHR-like F_2_ behavior; **Table S5**). Nineteen of these implicated hippocampal subregions or cell types, mostly neuronal (n=10), emphasizing GABA-ergic cells (n=3) and dendrites (n=3). The dentate gyrus was implicated (n=1), epithelial cells (n=4, including gene *C2cd3*) and vasculature (n=3, including *Mfge8*). Fourteen gene sets were derived from previous differential expression experiments ^53,54^, with most related to stress or fear conditioning (n=12, upregulated: n=9, including gene *C2cd3)*. Other implicated functions included nervous system development, proliferation, and cell fate (n=13, including genes *Mfge8, Nqo2, Ucp2,* and *C2cd3*) and transcription regulation (n=9, including *Ucp2*).

Thirty-seven gene sets were upregulated (FDR<0.05) with a bLR-like phenotype (*i.e.,* in bLRs and with bLR-like F_2_ behavior). Eleven of these implicated hippocampal subregions or cell types, especially microglia (n=8, including gene *Tmem144*). Other emphasized pathways included mitochondrial function, oxidative phosphorylation, and cellular respiration (n=6, including genes *Wdr93* and *Idh1*), metabolism (n=5, including *Pex11a, Lsr, Ist1,* and *Idh1*), and immune response (n=4). A non-directional analysis produced weaker results (12 gene sets with FDR<0.10) highlighting similar functions (metabolism: n=3, including *Pex11a, Lsr, Ist1, Mcee,* and *Idh1;* and microglia: n=4, including *Fcrl2* and *Tmem144)*. Gene sets related to a bLR-like model, Flinders Sensitive Line, were also highlighted (n=3).

### Constructing a hippocampal cis-eQTL database to determine which differential expression is most likely driven directly by proximal genetic variation

We integrated our current F_2_ RNA-Seq data (*n*=245) with previous genotyping results (*n*=4,425,349 single nucleotide polymorphisms (SNPs), ^34^) to identify 5,351 genes (eGenes) with hippocampal expression tightly correlated (FDR<0.05) with nearby genetic variation (*cis*-eQTLs: within +/-1 MB of the transcription start site (TSS)). Using stepwise regression, we identified additional conditionally-independent *cis*-eQTLs beyond the strongest *cis*-eQTL for each eGene (**Fig S5A**), distinguishing a final total of 5,937 *cis*-eQTLs representing 5,836 unique eVariants. Like previous *cis*-eQTL analyses, these eVariants were predominantly located within +/-400 kB of the TSS of their respective eGene (**Fig S5B**). A comparison with existing rat *cis*-eQTL databases (RatGTEx: ^55,56^, Mitchell et al, *unpublished*; and Telese et al, *unpublished*), **Fig S5C,** indicated that most hippocampal eGenes were also significant eGenes within at least four other tissues (out of 11 tissues characterized, **Fig S5D**), and confirmed that previously-identified brain *cis*-eQTLs showed a similar direction of effect on gene expression within the hippocampus (R=0.67-0.75, rho=0.65-0.77, **Fig S7-S8**) when there was at least a nominal (*p*<0.05) relationship in our dataset, although many *cis*-eQTLs remained region specific. As our hippocampal *cis*-eQTL database represents a valuable resource for the interpretation of rat genomic results, we have shared it on RatGTEx (https://ratgtex.org/download/study-data/#HPC_F2).

### bHR/bLR differential expression can be predicted using the hippocampal cis-eQTL database

We used our *cis*-eQTL database to predict the effect of genetic variation that segregates the bHR/bLR lines on gene expression. Many eVariants (2,452) showed at least partial bHR/bLR segregation in the F_0_ rats (n=10 bHR/n=10 bLR sequenced in ^34^), such that if all subjects from one phenotype (*e.g.,* bHRs) had 2 reference alleles (0/0), all subjects from the other phenotype had at least 1 alternate allele (0/1; population segregation statistic G_st’_>0.27 ^57^). To predict the effect of these bHR/bLR segregated eVariants on gene expression, we calculated the allelic Log2FC (aFC) for each eVariant and assigned the direction of effect based on the allele frequency within the bLR vs. bHR F_0_ rats (**Fig 8A**). These predictions correlated strongly with the F_0_ differential expression results (**Fig 8B-C**, 2,500 eGene/eVariant combinations: R=0.77, rho=0.63, *p*<2e-16) and our previous bHR/bLR late generation meta-analysis effect sizes (2,114 eGene/eVariant combinations: R=0.52, rho=0.61, *p*<2e-16, **Fig S9**). These results validated our hippocampal *cis*-eQTL database and confirmed that bHR/bLR differential expression of eGenes is likely driven by bHR/bLR genetic segregation.

**Fig 8.**
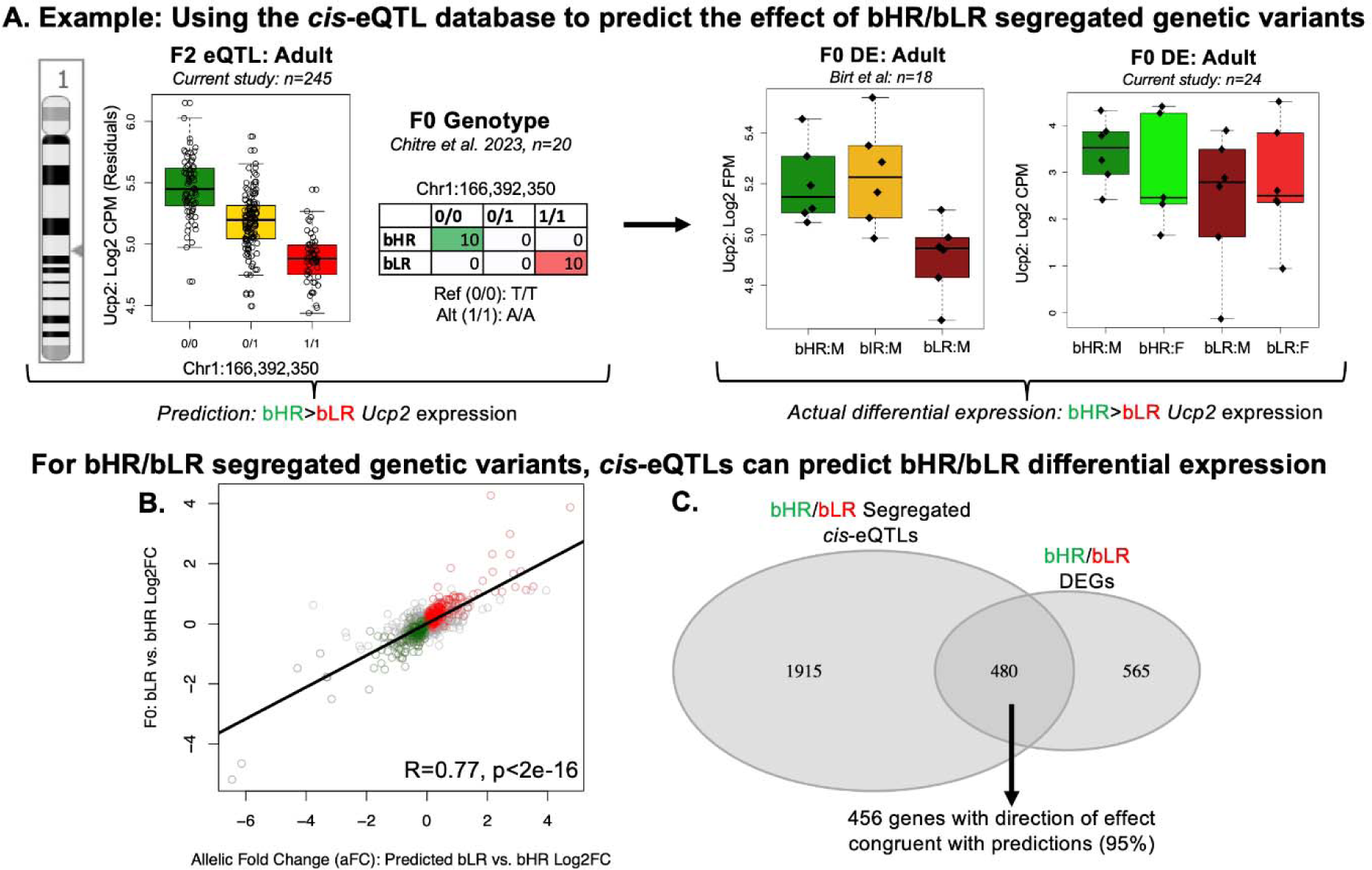
bHR/bLR differential expression (DE) related to segregated genetic variation can be successfully predicted using our hippocampal cis-eQTL database. Green vs. red coloring is used to indicate either bHR vs. bLR phenotype or the allele overrepresented in each respective phenotype. Gold is used to indicate heterozygotes (0/1). **A)** To create the cis-eQTL database, we integrated our current F_2_ transcriptional profiling data (n=245) with our previous whole genome sequencing results (n=4,425,349 single nucleotide polymorphisms (SNPs), ^34^) to identify genes with hippocampal expression tightly correlated with nearby genetic variation (cis-eQTLs, FDR<0.05). As an example, a boxplot illustrates a cis-eQTL for Ucp2 (Uncoupling Protein 2). Within this cis-eQTL, the alternate allele for the top eVariant (Chr1:166,392,350) is associated with decreased expression of Ucp2. In the boxplot, genotype (x-axis) is indicated by alternate allele count, with 0/0 (two reference alleles), 0/1 (heterozygote), and 1/1 (two alternate alleles). Gene expression (y-axis: Log2 CPM) is plotted as residual expression after quality control and controlling for technical co-variates included in our differential expression model (n=245). We used the cis-eQTL database to predict the effect of genetic variation that segregates the bHR/bLR lines (2,452 eVariants with partial segregation (G_st’_>0.27) in the F_0_ rats: n=20 ^34^) on gene expression, with the direction of effect for the allelic Log2 fold change (aFC) for each eVariant assigned to reflect bLR vs. bHR allele frequency (n=20 ^34^). To illustrate this, a table shows the bHR/bLR segregation for th top eVariant (Chr1: 166,392,350) for Ucp2. The alternate allele (1/1) for the top eVariant is mor prevalent in bLRs (red), whereas bHRs are more likely to carry the reference allele (0/0, green) ^34^. Since the alternate allele was associated with decreased hippocampal Ucp2 expression in our cis-eQTL analysis, we predict that bHRs would have greater Ucp2 expression than bLRs. This prediction is correct when we examine our previous differential expression results from the male bHR vs. bLR rats (example boxplot from our previous F_0_ sample ^43^: n=18, y-axis: Log2 FPM)). This prediction is also correct when we examine the differential expression results from our current F_0_ sample of male (M) and female (F) bHR and bLR rats (n=24, boxplot y-axis: Log2 CPM), although the effect appears larger in males. **B)** When considering the full sample of bHR/bLR segregated cis-eQTLs, there is a strong positive correlation between predicted bLR vs. bHR differential expression (scatterplot x-axis: bLR vs. bHR aFC) and our F_0_ differential expression results (y-axis: bLR vs. bHR Log2FC) (n=2,452 cis-eQTLs, R=0.77, p<2e-16). A similar positive correlation with bLR vs. bHR meta-analysis results is shown in **Fig S9**. Within the scatterplot, color indicates the subset of cis-eQTLs that were associated with differentially expressed genes upregulated in the bLRs (red) or bHRs (green) within the F_0_ differential expression study or bHR/bLR meta-analysis (**Fig 4F-H**) that had differential expression reflecting bHR/bLR segregation at their eVariant (n=492 cis-eQTLs representing 456 eGenes). This subset is also indicated with **C**) A Venn diagram illustrating overlap between the bHR/bLR differentially expressed genes (DEGs) (1,045 of which were present in the F_2_ dataset) with the significant eGenes identified in our hippocampal cis-eQTL database that had eVariants segregated in the bHR/bLR rats (2,395 eGenes). Out of the 480 genes satisfying both criteria, 456 (95%) showed a direction of effect in the differential expression results congruent with what would be predicted based on bHR/bLR genotype segregation.

### cis-eQTLs that strongly co-localize with QTLs for behavior are predominantly located on chromosome 1

We determined which hippocampal *cis*-eQTLs co-localized with regions of the genome associated with bHR/bLR-like behavior (QTLs) within the larger F_2_ sample (adults: n=323 adults, juveniles: n=216 ^34^) using Summary Data-based Mendelian Randomization (SMR; ^58^). We focused on QTLs for behaviors measured in F_2_ adults that were included in our differential expression analysis (LocoScore, EPM time immobile, EPM distance traveled, EPM % time in open arms, PavCA Index), and for analogous behaviors measured in an independent sample of F_2_ juveniles (open field (OF) time immobile, OF distance traveled, OF % time in center). This analysis identified 79 *cis*-eQTLs that were co-localized with QTLs for LocoScore (FDR<0.10), including 1 *cis*-eQTL that was also co-localized with a QTL for EPM distance traveled (FDR<0.10), and 13 of the 14 *cis*-eQTLs that were co-localized with QTLs for OF distance traveled (FDR<0.10). Most *cis*-eQTLs that strongly co-localized with behavioral QTLs were on chromosome 1, as expected due to the strength of the QTLs on this chromosome (**Fig 9A&B**).

**Fig 9.**
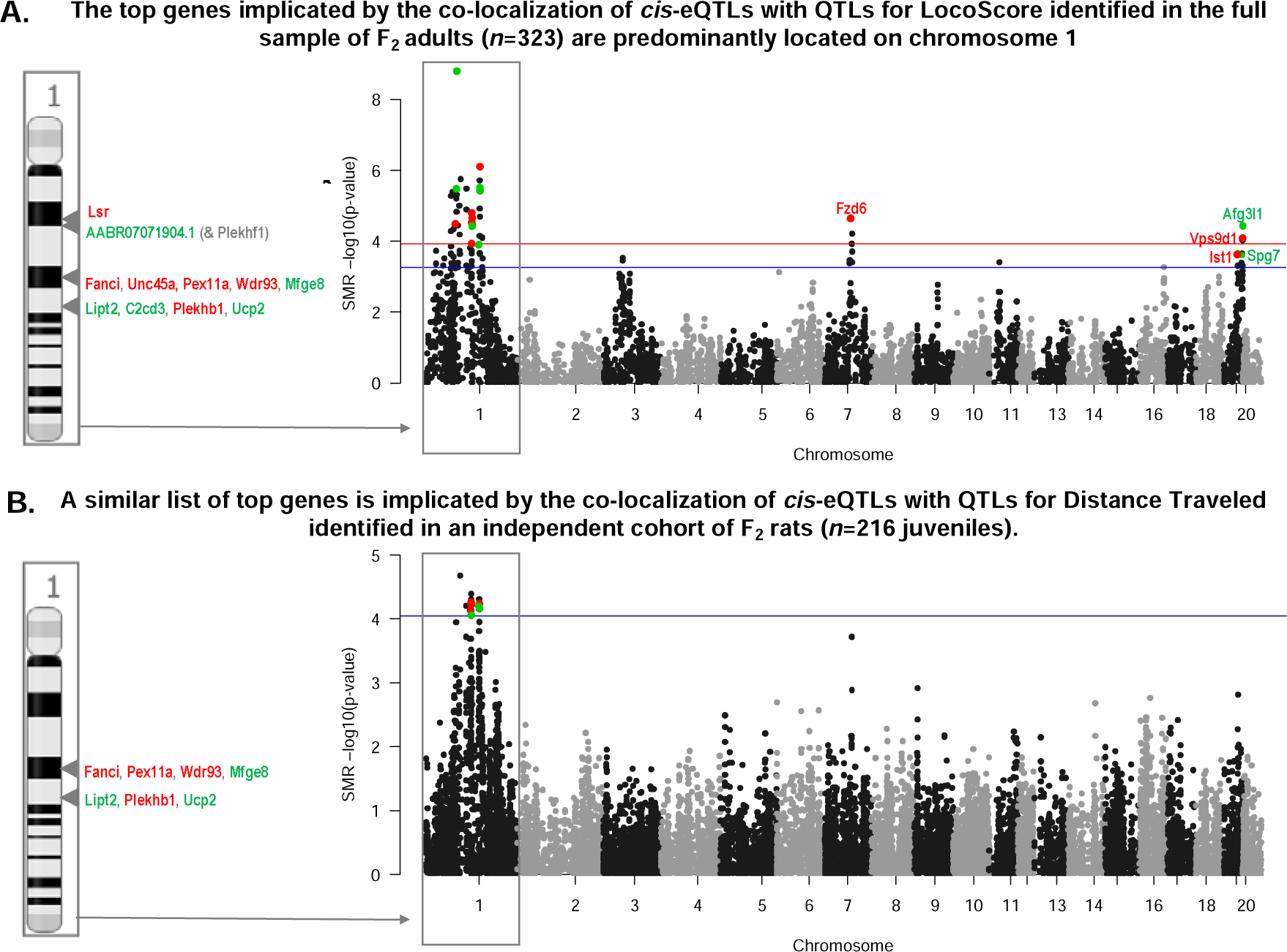
The top candidate genes for mediating the influence of genetic variation on behavioral temperament are located on chromosome 1. **A)** A Manhattan plot shows the co-localization of hippocampal cis-eQTLs with LocoScore QTLs identified in the full sample of F_2_ adults (n=323). The x-axis indicates the chromosomal location for all identified hippocampal cis-eQTLs (n=5937). Chromosomes are indicated with alternating black and grey coloring. The y-axis indicates the statistical significance (-log10(p-value)) for the co-localization as identified by the SMR analysis. The red line indicates FDR=0.05 and the blue line indicates FDR=0.10. Colored dots denote cis-eQTLs with FDR<0.10 that meet all additional desired criteria for being the most compelling candidates for mediating the effect of selective breeding on behavior. Red is used to indicate cis-eQTLs associated with genes upregulated with bLR-like behavior (decreased LocoScore), green is used to indicate cis-eQTLs associated with genes upregulated with bHR-like behavior (increased LocoScore). For labeling the cis-eQTLs with their respective gene symbols, a side panel that zooms in on chromosome 1 is used for clarity. **B)** A Manhattan plot shows the co-localization of cis-eQTLs with QTLs for open field distance traveled identified in an independent sample of F_2_ juveniles (n=216). Notably, a similar panel of cis-eQTLs on chromosome 1 are identified as meeting all desired criteria for being the most compelling candidates for mediating the effect of selective breeding on behavior.

To narrow down our pool of top candidate genes for mediating the effect of genetic variation on behavioral temperament, we used converging information from our different samples and analyses. First, we narrowed our scope to *cis*-eQTLs that we had confirmed are segregated in bHR/bLRs with differential expression matching predictions based on the distribution of alleles in the two lines (**Fig 8C**; 492 cis-eQTLs representing 456 eGenes). Within this subset of *cis*-eQTLs, the strongest co-localization with QTLs tended to predict F_2_ differential expression with behavior, especially when considering the predicted direction of effect based on the relationship between genotype and behavior within the larger F_2_ sample (adults: n=323 adults) and genotype and expression within the *cis*-eQTL analysis (n=245) (**Fig 10A**). This was particularly true for LocoScore (**Fig 10B**, R=0.56, *p*<2.2e-16), but also other F_2_ adult behaviors (**Fig S10**, R=0.33-0.57, all *p*<3.07e-14). It was also true when comparing F_2_ differential expression to SMR co-localization results with QTLs for two analogous juvenile behaviors (**Fig 10C**, **Fig S11,** OF distance traveled: R=0.35, p=1.48e-15, OF time immobile: R=0.26, p=4.44e-09).

**Fig 10.**
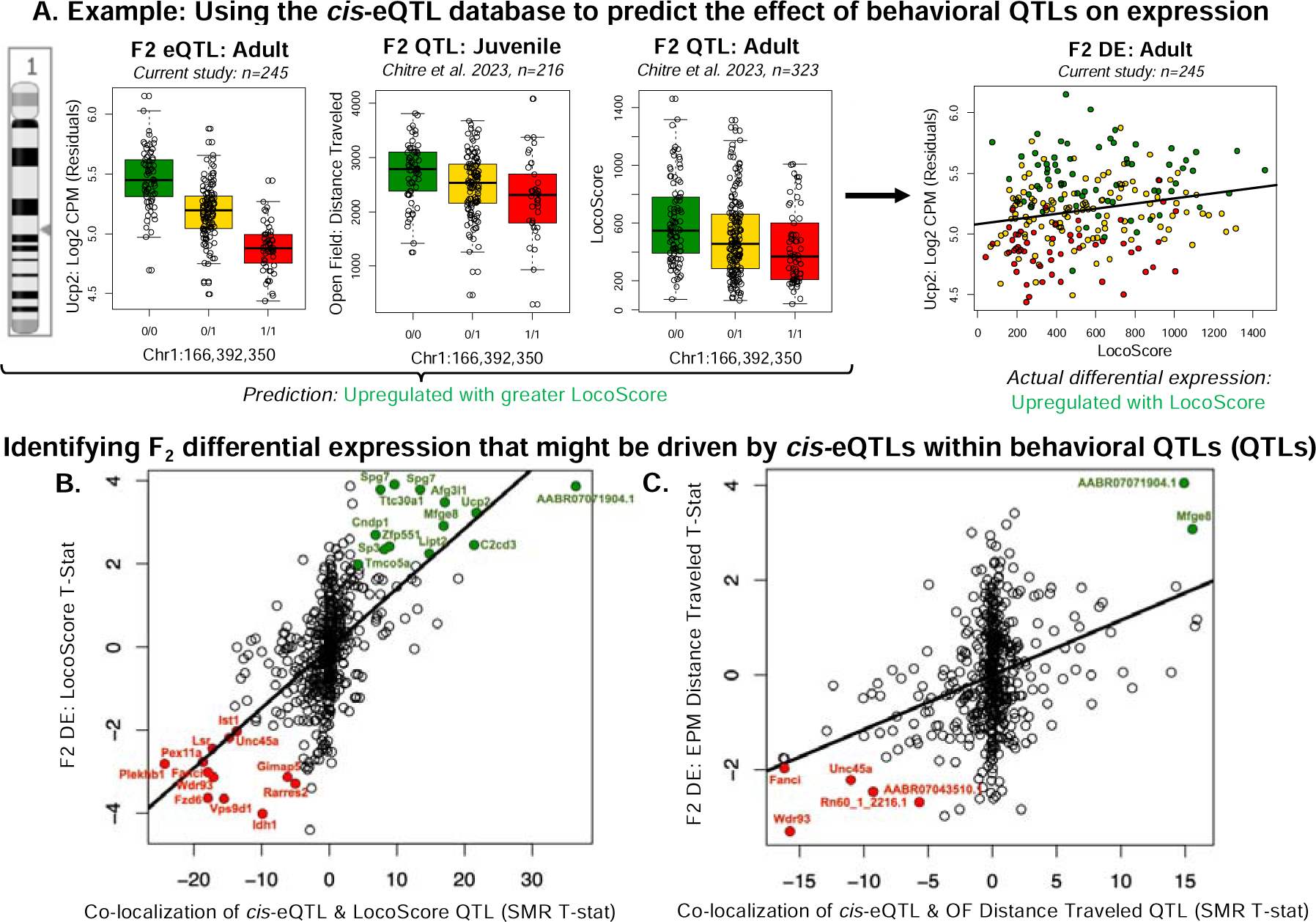
Our cis-eQTL database can also be used to predict the effect of genetic variation that correlates with behavior (QTLs) on gene expression. **A)** Following the plotting conventions of Fig 8, the boxplot illustrating the cis-eQTL for Ucp2 is shown again as an example (top eVariant: Chr1: 166,392,350), along with two boxplots illustrating the association between the alternate allele (1/1) for the top eVariant and decreased open field distance traveled in F_2_ juveniles (n=216; ^34^) and decreased LocoScore in the full sample of F_2_ adults (n=323; ^34^). Since the alternate allele was associated with decreased Ucp2 expression, we predict that decreased Ucp2 expression might also be associated with decreased distance traveled and LocoScore (i.e., positive correlation). This prediction is correct when we examine the differential expression (DE) results from the F_2_ rats (n=245). The scatterplot shows that Ucp2 was more highly expressed in hippocampus of F_2_ rats with a higher LocoScore. Similar to the eQTL plot, gene expression (Log2 CPM) is plotted as residual expression. **B-C)** Overall, some F_2_ differential expression related to behavior can be predicted by the co-localization of cis-eQTLs with QTLs for the behavior identified within the larger F_2_ sample (adults: n=323, juveniles: n=216, ^34^). This was particularly true when considering the subset of cis-eQTLs we confirmed had differential expression in bHR/bLRs matching what would be expected based on the segregated distribution of alleles in the two lines (see **Fig 8C**: n=492 cis-eQTLs representing 456 eGenes), but also weakly true within the full sample of cis-eQTLs (**Fig S10-S11**). The strength of the co-localization of hippocampal cis-eQTLs with regions of the genome associated with bHR/bLR-like behavior (QTLs) was determined using Summary Data-based Mendelian Randomization (SMR), and the direction of effect for the relationship between gene expression and behavior was predicted as described above. **B**) An example scatterplot shows the positive correlation between the strength of the co-localization of cis-eQTLs with the QTLs for LocoScore identified in the larger F_2_ sample (n=323 adults, x-axis: SMR T-statistic), with negative values indicating a predicted negative relationship between gene expression and LocoScore and positive values indicating a positive relationship between gene expression and LocoScore) and the differential expression for LocoScore (y-axis: Log2FC) (n=492; R=0.56, p<2.2e-16). Color is used to indicate the subset of genes that had nominal differential expression in the F_2_s for LocoScore (p<0.05) that matched the prediction based on the co-localization between their cis-eQTL and the QTL for LocoScore in the larger F_2_ sample (p<0.05), with green indicating bHR-like upregulation with increased LocoScore and red indicating bLR-like upregulation with decreased LocoScore. **C**) An example scatterplot shows the positive correlation between the strength of the co-localization of cis-eQTLs with the QTLs for open field distance traveled identified in an independent sample of F_2_ rats (n=216 juveniles, x-axis: SMR T-statistic), with predicted direction of effect assigned as discussed above, and the differential expression for EPM distance traveled in the F_2_ adults (y-axis: Log2FC) (n=492; R=0.35, p<1.48e-15). Coloring follows the conventions in panel E. **Fig S10-S11** contain scatterplots for other F_2_ adult and juvenile behaviors.

The most compelling candidates for mediating the effect of genetic variation on behavioral temperament should have expression strongly related to genetic variation (*cis*-eQTLs) that is segregated in bHR/bLR, correctly predicts bHR/bLR differential expression, and co-localizes with a QTL for behavior that correctly predicts F_2_ differential expression associated with that behavior (**Fig 2**). Among the SMR results, 16 of the 80 genes surviving FDR correction (FDR<0.10) met all these criteria (**Fig 11**, examples: **Figs S12-15**). By conservative estimate (**Supp. Methods**), one gene or less in our dataset is expected to show this degree of convergence due to random chance. These 16 genes were clustered within seven genomic regions on chromosomes 1, 7, and 19, suggesting that there remained some false discovery due to linkage disequilibrium (**Fig 9A&B**). That said, when cross-referencing with functional annotation, eight of these candidate genes -representing five of the identified regions - were clearly related to mitochondrial function and bioenergetics (**Fig 11**, **Fig 12**), hinting at the relevant genes in each region.

**Fig 11.**
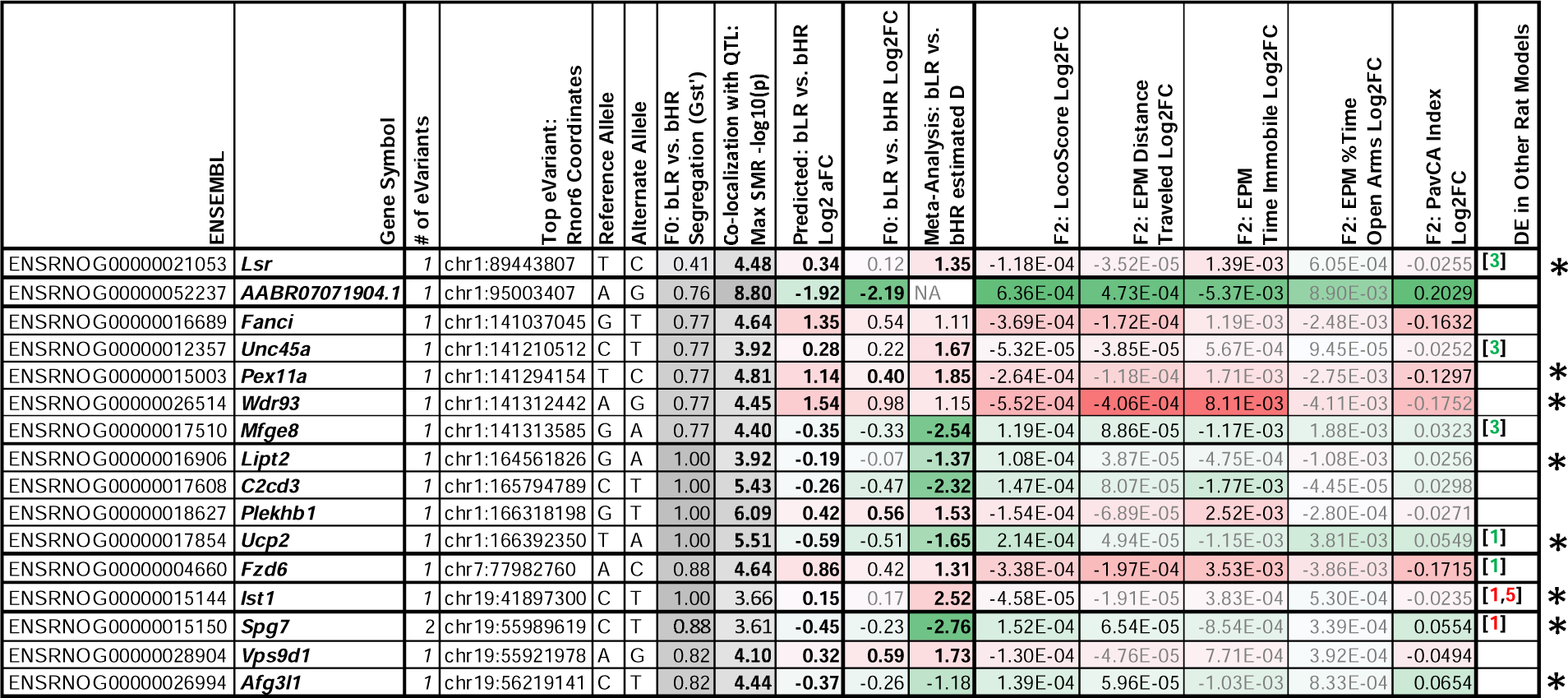
A table summarizing the converging evidence from genetic association and differential expression studies implicating 16 genes in behavioral temperament. To narrow down our pool of top candidate genes for mediating the effect of genetic variation on behavior, we used converging information from our different samples and analyses (Fig 2). We required that our top candidate genes have expression strongly related to genetic variation (cis-eQTLs) that was segregated in the bHR/bLR lines that correctly predicted bHR/bLR differential expression and co-localized with a QTL for behavior (SMR FDR<0.10) that correctly predicted at least nominal F_2_ differential expression associated with that behavior. The summary table follows the conventions of **Fig 7D**, but also includes the top eVariant associated with the expression of the gene within our cis-eQTL analysis, along with its reference and alternate alleles, its separation in our bred lines (G_st’_: ranges from 0 (no segregation) to 1 (fully segregated)), its co-localization with behavioral QTLs from the full adult and juvenile F_2_ samples (maximum -log10(p-value) from the SMR analysis, bold=FDR<0.05, black=FDR<0.10), and the differential expression that is predicted due to bLR vs. bHR segregation at the eVariant (allelic Log2 fold change or Log2aFC, bold=FDR<0.05). Genes with functions related to bioenergetics are indicated with an * and illustrated in Fig 12. Full gene names (when applicable): Lsr: Lipolysis Stimulated Lipoprotein Receptor; Fanci: FA Complementation Group I; Unc45a: Unc-45 Myosin Chaperone A; Pex11a: Peroxisomal Biogenesis Factor 11 Alpha; Wdr93: WD Repeat Domain 93; Mfge8: Milk Fat Globule EGF And Factor V/VIII Domain Containing; Lipt2: Lipoyl(Octanoyl) Transferase 2; C2cd3: C2 Domain Containing 3 Centriole Elongation Regulator; Plekhb1: Pleckstrin Homology Domain Containing B1; Ucp2: Uncoupling Protein 2; Fzd6: Frizzled Class Receptor 6; Ist1: IST1 Factor Associated With ESCRT-III; Spg7: SPG7 Matrix AAA Peptidase Subunit, Paraplegin; Vps9d1:VPS9 domain containing 1; Afg3l1: AFG3-like AAA ATPase 1.

**Fig 12.**
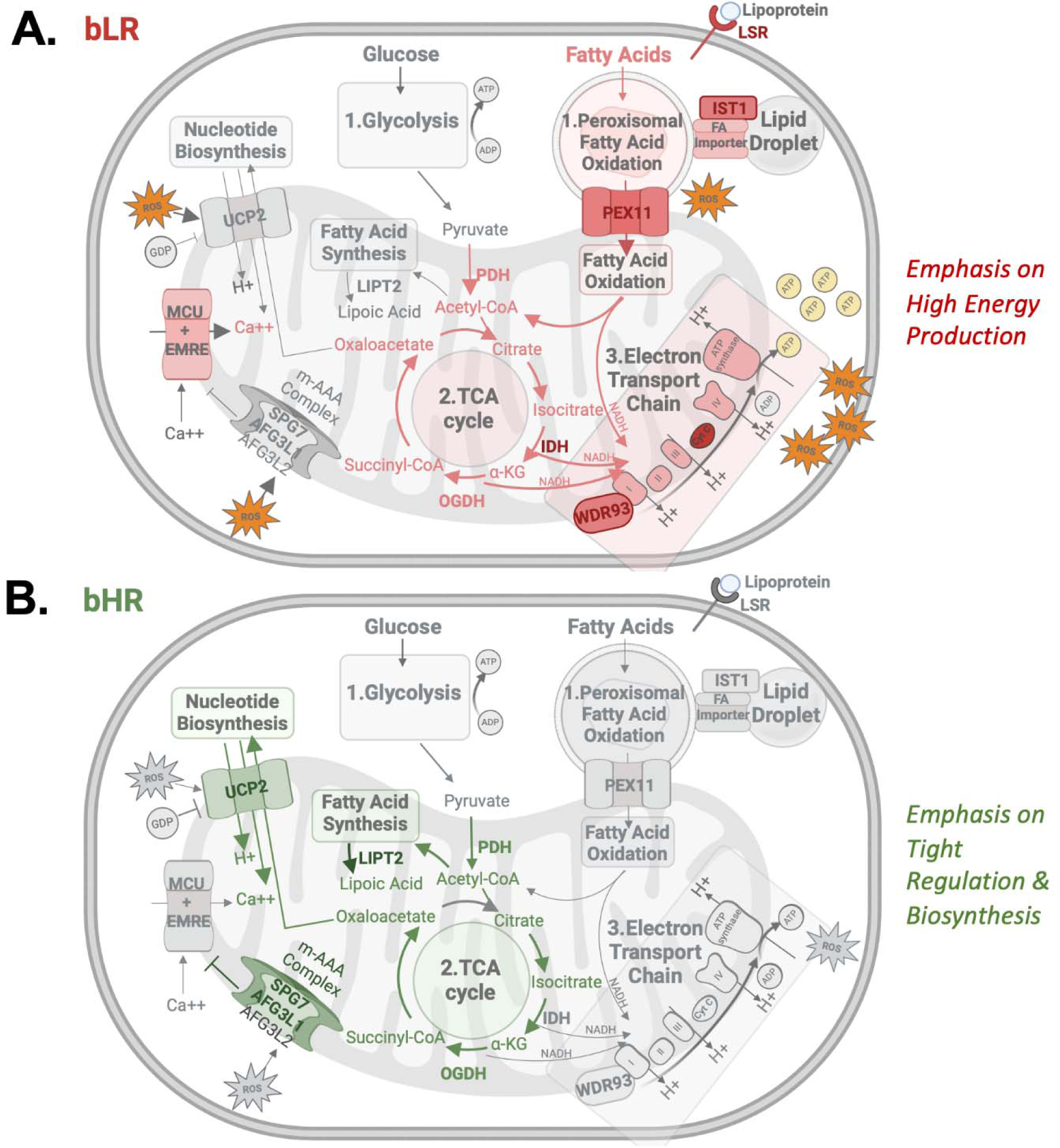
The differentially expressed genes implicated as top candidates for mediating the effect of selective breeding on behavior are often regulators of bioenergetic function. Red/pink indicates that a function is likely to be increased in bLR-like animals, green indicates increased function in bHR-like animals. **1**. In the brain, energy is primarily released from glucose within a series of biochemical reactions starting with glycolysis ^188^, but it can also be released from other energy sources, such as fatty acid oxidation ^61^. **2.** Metabolites from these processes are fed into the tricarboxylic acid (TCA) cycle within the mitochondrial matrix, which generates electron donors (NADH, FADH2) **3.** Electron donors feed into the electron transport chain, which moves protons across the inner mitochondrial membrane to produce a gradient capable of driving energy output (adenosine triphosphate: ATP). This process produces reactive oxygen species (ROS) as a byproduct ^189^. **A)** bLR-like animals have a pattern of upregulated expression suggesting elevated oxidative phosphorylation, but potentially also reduced sensitivity to cellular need in a manner leading to excessive ROS production and neuroimmune activation under conditions of elevated activity such as stress. This conclusion is supported by: **i)** previous evidence of elevated oxidative phosphorylation, including elevated activity within the electron transport chain ^60^, **ii)** Upregulation of multiple gatekeepers of fatty acid oxidation, which is a form of energy production that can release twice as much energy as glucose metabolism ^62^. These gatekeepers include lipolysis stimulated lipoprotein receptor (Lsr), which uptakes lipoproteins into the cell, IST1 factor associated with ESCRT-III (Ist1), which facilitates fatty acid trafficking into peroxisomes to begin fatty acid oxidation ^66^, and peroxisomal biogenesis factor 11 alpha (Pex11a), which encodes a fatty acid oxidation rate-limiting channel that allows lipids and fatty acid metabolites to pass from peroxisomes into mitochondria ^63–65^. **iii)** Upregulation of Idh1, encoding the isocitrate dehydrogenase 1 enzyme in the TCA cycle, which is the most important producer of the electron donor NADH in the brain ^67,190^, **iv)** Upregulation of WD repeat domain 93 (Wdr93), which is theorized to be an accessory subunit to Complex 1 in the mitochondrial electron transport chain, increasing ATP production ^68–70^. **v)** Down-regulated expression of subunits of the m-AAA (ATPases Associated with a variety of cellular Activities) complex (Spastic paraplegia type 7 (Spg7), AFG3-like protein 1 (Afg3l1)). Decreased m-AAA complex function causes constitutive activity of the mitochondrial calcium uniporter (MCU)^83,84^. Elevated mitochondrial calcium influx activates enzymes within the TCA cycle (PDH, IDH, and OGDH)^67^, and, if it becomes excessive, triggers ROS production and apoptosis ^83^. **B)** bHR-like animals have a pattern of upregulated expression suggesting that energy production is kept under tight regulation, potentially limiting oxidative phosphorylation but allowing for greater biosynthesis. These findings include **i)** Upregulated Uncoupling Protein 2 (Ucp2), encoding a mitochondrial transporter which promotes homeostasis and decreased ROS production by decreasing the mitochondrial proton gradient, exporting the rate-limiting substrate for the TCA cycle (oxaloacetate), and regulating calcium influx ^73–77,82^.; **ii)** Upregulated expression related to m-AAA complex function (Spg7, Afg3l1), which ensures that mitochondrial protein availability, including an essential regulator (EMRE) of the MCU, does not exceed cellular need. **iii)** Upregulated lipoyl(octanoyl) transferase 2 (Lipt2), which senses the input to the TCA cycle (Acetyl-CoA) and stimulates TCA cycle enzyme activity accordingly via the mitochondrial fatty acid synthesis pathway ^78–80^.

## Discussion

We have selectively bred rats for high or low propensity to explore a novel environment for many generations to produce two lines (bHRs, bLRs) with extreme, stable differences in behavior. These phenotypes resemble human externalizing and internalizing tendencies which underlie many psychiatric disorders, including substance use disorders. Our current study revealed that our selective breeding paradigm produced a robust molecular phenotype in the hippocampus in both sexes, confirming results from our previous multi-generational bHR/bLR meta-analysis ^43^. Interbreeding bHRs and bLRs produced a large cohort of F_2_ intercross rats that showed a range of intermediate phenotypes. Behaviors that diverged in the bHR/bLR lines, including anxiety-like and reward-related behavior ^26,43,59^, remained correlated with exploratory locomotion in our F_2_ sample, allowing us to investigate their shared etiology. Hippocampal gene expression related to bHR/bLR lineage predicted gene expression related to F_2_ behavior, including exploratory locomotion and anxiety. Six genes showed consistent differential expression with behavioral phenotype in bHR/bLR and F_2_ intercross rats, as well as in multiple other rat models targeting similar behavior.

Selective breeding should produce an enrichment of genetic alleles influencing the phenotype under selection. To determine which differential expression may directly mediate the effect of selective breeding on behavioral temperament, we identified hippocampal expression that was strongly correlated with genetic variation in the F_2_s (*cis*-eQTLs). This *cis*-eQTL database allowed us to accurately predict differential expression related to bHR/bLR genetic segregation. We also identified differential expression associated with F_2_ behavior that matched what would be expected due to the co-localization of *cis*-eQTLs with behavioral QTLs previously identified in the larger F_2_ cohort (adults and juveniles) ^34^. This converging evidence highlighted 16 genes within 7 genomic regions on chromosomes 1, 7, and 19 as strong candidates for mediating the effect of selective breeding on behavioral temperament.

### Functional patterns: Bioenergetic regulation of hippocampal function

Among these 16 top candidate genes, eight are directly involved in bioenergetics (**Fig 12**). The differential expression results overall similarly showed upregulation in gene sets related to mitochondria, oxidative phosphorylation, and metabolism in bLR-like vs. bHR-like animals. These findings complement previous evidence that adult bLRs have elevated oxidative phosphorylation in the hippocampus, as indicated by increased mitochondrial oxygen consumption and elevated electron transport chain activity ^60^. Since the expression of our top candidate genes was strongly correlated with genetic variation tied to behavioral phenotype in both bHR/bLR and F_2_ samples, our results imply that variation in energy production may mediate the effect of heredity on temperament and provide insight into the responsible mechanisms.

In particular, our results suggest that bLR-like animals have enhanced fatty acid oxidation, which is a pathway that is particularly important during times of high energy usage ^61^ because it can release twice as much energy as glucose metabolism ^62^. bLR-like animals had upregulation of multiple fatty acid oxidation gatekeepers (*Lsr, Ist1,* and *Pex11a* ^63–66^). Downstream, there was also upregulation that could facilitate the tricarboxylic acid (TCA) cycle (*Idh1*) ^67^ and electron transport chain *(Wdr93)* ^68–70^ to increase energy production. These findings have widespread functional implications, as the brain consumes disproportionate energy to maintain neurotransmission and synaptic repolarization ^71^, especially during times of heightened activity, such as stress ^72^.

In contrast, energy production in bHR-like animals may be kept under tight regulation by upregulation of *Spg7, Afg3l1, Ucp2,* and *Lipt2. Ucp2* encodes a mitochondrial transporter and anion carrier that promotes homeostasis by serving as a metabolic switch, decreasing TCA cycle function ^73^ and mitochondrial proton gradient ^74–77^. *Lipt2* plays a similar feedback role, coupling TCA cycle enzyme activity to its input via the mitochondrial fatty acid synthesis pathway ^78–80^. *Spg7* and *Afg3l1* encode subunits of the m-AAA complex, which tailors mitochondrial protein levels to cellular need ^81^. Moreover, *Ucp2, Spg7* and *Afg3l* all regulate mitochondrial calcium intake ^82–84^, which couples energy production to synaptic activity by stimulating TCA cycle enzymes ^67,85^. As discussed below, this tight regulation may limit oxidative phosphorylation under some conditions, but also reduce reactive oxygen species production and allow for greater biosynthesis.

### Bioenergetics and behavior

Our results bolster growing evidence that bioenergetic genes and pathways regulate behaviors like exploratory activity, anxiety, and reward learning. The mitochondrial m-AAA complex, fatty acid oxidation pathway, and fatty acid synthesis feedback pathway are all critical for movement and motor activity in animals and humans ^61,83,86–88^, with severe, pathogenic mutations in *Spg7, Afg3l1,* and *Ist1* producing hereditary paraplegia and ataxia ^61,89,90^, sometimes with altered cognition, executive function, and social/emotional function ^90–94^. Logically, more subtle changes within these pathways could alter exploratory activity.

Energy production is also theorized to critically modulate the energy-demanding circuitry necessary for behavioral inhibition ^95,96^, and some of our candidate bioenergetic genes are more broadly implicated in behavioral temperament. *Ucp2* knock-out animals consistently demonstrate bLR-like behaviors, including decreased exploration, and anxiety- and depressive-like behaviors, especially following stress ^97–103^. Human GWAS also link *UCP2, SPG7,* and *WDR93* to the stress response, psychiatric disorders, externalizing behavior, and substance use disorders ^23,104–110^.

Metabolic differences have been observed in humans and animal models with anxiety and internalizing-like behavior ^111–115^ and hyperactivity and externalizing-like behavior ^116–119^. Our findings suggest that genetic vulnerability may contribute to these metabolic differences, bolstering support for metabolic interventions in psychiatry (e.g., ^113,115,119,120^). That said, the evidence linking energy production to internalizing-like vs. externalizing-like behavior is inconsistent across measurements and models, suggesting that the critical vulnerability may lie downstream in bioenergetically regulated functions like apoptosis, oxidative stress, and biogenesis ^113^. We have evidence supporting each of these possibilities.

### Bioenergetics: Role in reactive oxygen species production

During fatty acid oxidation and oxidative phosphorylation, reactive oxygen species are produced as a byproduct ^71,121^. Both energy production and reactive oxygen species increase with elevated synaptic activity and environmental stress ^71,72,122^. Thus, many metabolic genes are regulators of oxidative stress, with upregulation in bLR-like animals linked to greater oxidative stress and upregulation in bHR-like animals sometimes appearing protective (e.g., *Ucp2*, *Spg7/Afg3l1*, *Pex11a,* ^76,98,99,123–125^). bHR-like animals also had upregulation of protective *Mfge8* ^126^ and *Nqo2*, which can enhance reactive oxygen species production or reduce oxidative stress ^127–129^ in a manner important for encoding novelty in hippocampal interneurons ^130^ and potentially stress-related disorders ^131^.

These results bolster evidence that natural and genetically-selected variation in anxiety is consistently associated with markers of oxidative damage in animals and humans ^113^. Reactive oxygen species are also implicated in the development of anxiety and depressive-like behavior following chronic stress ^71,132^. The HPC is particularly vulnerable to oxidative stress ^122^ and accumulating evidence implicates oxidative stress in psychiatric disorders, including internalizing disorders and comorbid substance abuse ^71,72,132–136^.

### Bioenergetics: Role in neuroimmune activation

Gene sets related to immune activation and microglia were upregulated in bLR-like animals. This upregulation may be driven by bLR/bHR bioenergetic differences: both the ATP and reactive oxygen species produced by fatty acid oxidation and oxidative phosphorylation can cause microglial activation ^137,138^ and promote microglial release of pro-inflammatory factors ^139,140^. Notably, two of the top candidates upregulated in bHR-like animals, *Ucp2* and *Mfge8*, are also master regulators of microglial activation, promoting an anti-inflammatory and pro-repair state ^141–143^. Both *Mfge8* and *Ucp2* encourage microglial engulfment of damaged cells and unwanted synapses. Disrupting this process causes hippocampal dysfunction, inflammation, anxiety-like behavior, insomnia, and depressive-like behavior ^103,144–146^, mirroring a bLR-like behavioral phenotype. *Fcrl2* was also upregulated in bHR-like animals and is likely abundant in microglia, dampening immune responses ^147,148^. In contrast, *Tmem144* was upregulated in bLR-like animals in our study and three others ^44–46^ and is highly expressed in microglia during development ^149,150^, but with unknown function.

These findings complement previous findings that bLR microglia exhibit an “intermediate activation” hyper-ramified morphology ^151^ resembling that observed following chronic stress ^152^, when reactive oxygen species and microglial activation are critical for the development of anxiety-like behavior ^139,153^. Moreover, inhibiting microglial activity reduced bLR-like behavior ^151^. Microglial activation has also been implicated in affective and substance use-related behaviors ^146,154^.

Neuroimmune activation could also be caused by mitochondrial regulation of apoptosis and cell death. Excessive mitochondrial calcium intake, decreased m-AAA complex function, decreased *Spg7*, and decreased *Lipt2* can all trigger the mitochondrial membrane potential collapse that drives apoptosis ^83,88,155^. m-AAA complex deficiencies can also cause dysfunctional mitochondrial protein synthesis, respiration, transport, and fragmentation ^83^ and are linked to neurodegeneration ^83,84^ whereas *Ucp2* is considered neuroprotective ^156,157^. Therefore, down-regulation of *Spg7*, *Afg3l1, Lipt2,* and *Ucp2* in bLR-like animals might increase risk for cell loss and neuroimmune activation, especially after periods of intense neuronal activity, such as occurs during stress ^72^.

### Bioenergetics: Role in growt

bHR/bLR bioenergetic differences may also contribute to the upregulation of gene sets related to nervous system development and proliferation in bHR-like animals. Both energy availability and use exert control over proliferation, cell differentiation, and growth-related processes, and biosynthesis using glucose-derived products directly competes with oxidative phosphorylation for essential substrates ^158^. Therefore, many of the candidate metabolic genes also influence proliferation and growth (e.g., *Ucp2*, *Lsr*, *Lipt2*: ^159–163^). Other top candidates regulate growth-related processes, including *Mfge8* and *Fzd6* ^164–166^. *Fzd6* has also been linked to anxiety and depressive-like behavior ^167,168^. These results are noteworthy due to known bHR/bLR differences in neurogenesis, proliferation, and growth factor response ^32,41,43^, and extensive literature implicating both hippocampal atrophy in internalizing disorders and growth-related processes in antidepressant function ^169^.

### Remaining Questions and Limitations

*AABR07071904.1* is a compelling candidate, with a *cis*-eQTL near the strongest LocoScore QTL peak ^34^. According to genome assembly Rnor6 (Ensembl v103), *AABR07071904.1* generates long non-coding RNA, but in mRatBN7.2 (Ensembl v106) the gene was retired, potentially mapping to *Zfp939-201*. In either form, it could play some important, unknown regulatory role, but it is noteworthy that the implicated *cis*-eQTL is in linkage disequilibrium with a missense coding variant for *Plekhf1* (**Fig S12,** ^34^). *Plekhf1* was not differentially expressed in our study, but has been linked to stress and mood ^34^. This ambiguity highlights a limitation to our method: we identified single nucleotide variants that may mediate effects on behavior via gene expression levels, but other mechanisms may contribute to our phenotype, including coding variants, structural variants, epigenetic modifications, tissue, and context-dependent activity. Future work will address these gaps, and further characterize gene expression at a single cell and circuit level.

One limitation of our approach is that loci which influence behavior are often part of relatively large haplotypes that may include multiple eQTLs. While some of those eQTLs may be the causal molecular intermediate that mediates the effect of a particular locus on behavior, others are merely in linkage disequilibrium with the causal locus. As a result, our approach, which emphasizes the effect of eQTLs on gene expression, will identify a mixture of truly causal genes and genes that are differentially expressed by eQTLs that are coincidentally in linkage disequilibrium with the behavioral QTLs. We have addressed this limitation by integrating individual genes into higher order biological concepts. This approach should be robust to the presence of some false positives.

### Conclusion

The bHR/bLR selectively bred rat lines display temperament vulnerabilities akin to those observed in humans. By crossbreeding bHR/bLR rats to produce a F_0_-F_1_-F_2_ intercross design, the key behaviors that diverged in the bHR/bLR lines remain correlated in the F_2_s and can be linked to hippocampal function. Transcriptional profiling revealed genes that not only exhibited robust bHR/bLR differential expression consistent with previous results ^43^, but also expression correlated with exploratory and anxiety-like behavior in the F_2_s. Using a *cis*-eQTL analysis, we identified differentially expressed genes that were linked to bHR/bLR segregated genetic variants that co-localized with genomic regions implicated in behavior. These genes are strong candidates for mediating the influence of selective breeding on temperament and related behavior, including exploratory locomotion, anxiety, and reward learning. Functional themes within our findings suggest that the bioenergetic regulation of oxidative stress, microglial activation, and growth-related processes in the hippocampus may play a role in shaping behavioral temperament, thereby modulating vulnerability to psychiatric and addictive disorders.

## Methods

Full methods are in the **Supplement**, including the MDAR and ARRIVE reporting checklists (Appendix 1&2). Analysis code (R v.3.4.1-v.4.2.2, R-studio v.1.0.153-v.2022.12.0+353) has been released at https://github.com/hagenaue/NIDA_bLRvsbHR_F2Cross_HC_RNASeq.

All procedures were conducted in accordance with the National Institutes of Health Guide for the Care and Use of Animals and approved by the Institutional Animal Care and Use Committee at the University of Michigan.

### Animals

Selectively breeding rats for high or low locomotor activity in a novel environment (LocoScore) produced the bHR line (Wakil:bHR, RRID:RGD_405847397) and bLR line (Wakil:bLR, RRID:RGD_405847400), respectively ^25^. After 37 generations, 12 bHRs and 12 bLRs (F_0_) were chosen from 24 distinct families to crossbreed. The F_1_ offspring with similarly high or low LocoScores were then bred with each other to produce a re-emergence of diverse behavioral phenotypes in the F_2_ generation (**Fig 1**). These 48 F_2_ litters generated 540 rats (n=216 behaviorally phenotyped as juveniles (1 month old), n=323 behaviorally phenotyped as young adults (2-4 months old)). Our current study sampled a subset of the adults (F_0_: n=24, n=6/phenotype per sex; F_2_: n=250, n=125/sex) that overlapped with previous genetic studies ^34,35^ but was distinct from our previous male-only meta-analysis ^43^.

### Behavioral data analysis

All F_0_ and F_2_ rats were tested for LocoScore (protocol: ^25^) and exploratory and anxiety-related behaviors on an EPM (protocol: ^34^, distance traveled (cm), time immobile (sec), and % time in the open arms). A subset of F_2_s (n=209) subsequently underwent seven sessions of PavCA training. The PavCA Index from sessions six and seven was used to reflect sign- and goal-tracking tendencies (defined in: ^170^).

### Tissue dissection, RNA extraction, and sequencing

Adults (postnatal days 113-132) were decapitated without anesthesia and brains rapidly extracted (<2 min). For the F_0_s, whole hippocampus was immediately dissected and flash frozen. For the F_2_s, whole brains were flash frozen, and later hole punches from the dorsal and ventral hippocampus were pooled from four hemisected coronal slabs per rat (-2.12 to -6.04 mm Bregma; ^171^). RNA was extracted using the Qiagen RNeasy Plus Mini Kit. A NEB PolyA RNA-seq library was produced and sequenced using a NovaSeq S4 101PE flowcell (targeting 25 million reads/sample).

### RNA-Seq analysis

RNA-Seq data preprocessing was performed using a standard pipeline including alignment (STAR 2.7.3a: genome assembly Rnor6), quantification of gene level counts (Ensembl v103, Subread version 2.0.0), and basic quality control. All downstream analyses were performed in Rstudio (v.1.4.1717, R v. 4.1.1). Transcripts with low-level expression (<1 read in 75% of subjects) were removed. Normalization included the trimmed mean of M-values (TMM) method (^172^, *edgeR* v.3.34.1; ^173^), and transformation to Log2 counts per million (Log2 cpm ^174^, *org.Rn.eg.db* annotation v.3.13.0; ^175^). Following quality control, the F_0_ dataset contained n=23 subjects (subgroups: *n*=5 bHR females, n=6 for each of the other subgroups: bHR males, bLR females, bLR males) with Log2 cpm data for 13,786 transcripts, and the F_2_ dataset contained *n*=245 subjects (subgroups: *n*=122 males, *n*=123 females) with Log2 cpm data for 14,056 transcripts.

Differential expression was calculated using the limma/voom method (^176^, package: *limma* v.3.48.3) with empirical Bayes moderation of standard error and FDR correction. For the F_2_s, the same differential expression model was used for each variable of interest (LocoScore, EPM % time in open arms, EPM distance traveled, EPM time immobile, PavCA Index). Technical co-variates were included if they were strongly related to the top principal components of variation identified using Principal Components Analysis (PCA) or had confounding collinearity with variables of interest (covariates: percent of reads that were intergenic (%intergenic) or ribosomal RNA (%rRNA), dissector, sequencing batch, and PavCA exposure (“STGT_Experience”)).

*Equation 1: F_0_*differential expression *model:*

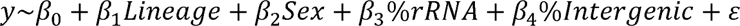

*Equation 2: F_2_*differential expression *model:*

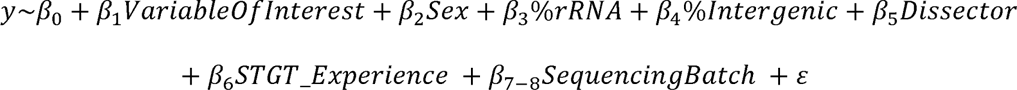

### Comparison of F_0_, F_2_, and previous meta-analysis results

The full F_0_ and F_2_ differential expression results were compared to our previous meta-analysis of late generation bHR/bLR hippocampal RNA-Seq studies (Table S2 in ^43^) using parametric and non-parametric methods (Pearson’s and Spearman’s correlation of Log2FC values) and visualized using gene rank-rank hypergeometric overlap (*RRHO v. 1.38.0* ^177,178^, ranking by t-statistics) and *VennDiagram* (v.1.7.3, ^179^). For downstream analyses, we defined bHR/bLR differentially expressed genes as the 1,063 genes with FDR<0.10 in either the F_0_s or late generation meta-analysis, or nominal replication (p<0.05) in both with consistent direction of effect.

### Gene Set Enrichment Analysis

To elucidate functional patterns, we ran Gene Set Enrichment Analysis (*fgsea v.1.2.1*, nperm=10,000, minSize=10, maxSize=1000, FDR<0.05) using a custom gene set database (Brain.GMT v.1, ^180^) that included standard gene ontology, brain cell types, regional signatures, and differential expression results from public databases. We created a continuous variable representing bLR-like vs. bHR-like differential expression for each gene by averaging the t-statistics for bLR vs. bHR comparisons in the F_0_ dataset and former late generation RNA-Seq meta-analysis ^43^ and for each of the F_2_ behaviors (with bHR-like phenotype set as reference). A second non-directional analysis used the absolute value of the average t-statistic.

### Constructing a hippocampal cis-eQTL database

Hippocampal *cis-*eQTL mapping was performed using published methods ^55^. Quality-controlled F_2_ RNA-Seq data (Log2 CPM, *n*=245 following quality control) was corrected for technical covariates (*Eq.2,* residualized), followed by rank-based inverse normal transformation ^55,181^. F_2_ genotypes were generated by low coverage whole genome sequencing followed by imputation (from data release for ^34^: https://doi.org/10.6075/J0K074G9, *n*=4,425,349 SNPs). Principal Components Analysis was run on the gene expression matrix and genotype matrix (following pruning for linkage disequilibrium, *Plink2 v.2.00a2.3* ^182^), and principal components 1-5 from both analyses included as covariates within the single-SNP linear regression for *cis*-eQTL mapping (*tensorQTL v*.*1.0.6* ^183^). We tested SNPs with minor allele frequency (MAF) >0.01 within +/-1 Mb of each gene’s transcription start site (tss). A significant eVariant-eGene relationship (*cis*-eQTL) was defined using empirical beta-approximated p-values calculated using permutations for each gene, and false discovery corrected (FDR<0.05) using results from the top SNP for all genes. When SNPs were in perfect linkage disequilibrium, a single SNP was selected randomly. Additional, conditionally independent *cis*-eQTLs for each eGene were identified using stepwise regression (*tensorQTL:* default settings). We estimated *cis*-eQTL effect size (allelic fold change, aFC) using an additive *cis*-regulatory model (package *aFC.py)* with raw expression read counts and the same covariates as *cis*-eQTL mapping^184^.

### Predicting bHR/bLR differential expression using the cis-eQTL database

We extracted F_0_ genotype information for each eVariant (n=10 bHR/n=10 bLR in ^34^, release: https://doi.org/10.6075/J0K074G9) using *VcfR* (v1.14.0, ^185^, *https://cran.r-project.org/web/packages/vcfR/vcfR.pdf*). We defined partial bHR/bLR segregation using *myDiff()* G_st’_>0.27 (^57^, akin to all 0/0 vs. all 0/1 in our dataset). We assigned the direction of effect for the Log2 aFC to reflect the bLR-enriched allele vs. bHR-enriched allele and compared these predictions to both the F_0_ differential expression results (Log2FC) and bHR/bLR late generation RNA-Seq meta-analysis results (estimated *d*) using parametric (linear regression) and non-parametric (Spearman’s rho) methods.

### Co-localization of cis-eQTLs with regions of the genome associated with bHR/bLR-like behavior

We used Summary Data-based Mendelian Randomization (SMR; ^58^) to test for colocalization of *cis*-eQTLs with QTLs from the full F_2_ cohort (GWAS results: ^34^) for adult behaviors included in our differential expression analysis (LocoScore, EPM time immobile, EPM distance traveled, EPM % time in open arms, PavCA Index) and juvenile behaviors targeting analogous traits (OF time immobile, OF distance traveled, OF % time in center). *Z*-scores for the *cis*-eQTL and GWAS associations with each top eVariant were used to calculate the SMR approximate chi-squared test statistic, with *p*-values determined using the chi-squared distribution’s upper tail (df=1, FDR correction: *mt.rawp2adjp()* (proc=”BH”) in *multtest* v.2.26.0, ^186^). Results were visualized using the *manhattan()* plot function in the *qqman* package (v.0.1.8, ^187^).

To determine whether the strength of *cis*-eQTL/QTL co-localization (SMR t-statistic) correlated with F_2_ behavioral differential expression, we assigned a predicted direction of effect based on the relationship between genotype and behavior within the larger F_2_ sample (adults: n=323 adults, juveniles: n=216) and genotype and expression within the *cis*-eQTL analysis (n=245). We then examined the correlation between the F_2_ Log2FCs for each adult behavior and the “directional” SMR T-statistics for the same adult behavior or analogous juvenile behavior (OF distance traveled, OF time immobile, OF % time in center), both in the full dataset (all 5937 cis-eQTLs) and within the subset of *cis*-eQTLs that we had already confirmed were segregated in bHR/bLRs in a manner predictive of bHR/bLR differential expression (492 cis-eQTLs representing 456 eGenes).

To narrow down our pool of top candidate genes for mediating the effect of genetic variation on behavior, we required that our final top candidate genes have expression strongly related to genetic variation (*cis*-eQTLs) that is segregated in bHR/bLR (G_st’_>0.27) in a manner that correctly predicts bHR/bLR differential expression and co-localized with a QTL for behavior (SMR FDR<0.10) in a manner that correctly predicts F_2_ differential expression. Using a conservative estimate (**Suppl. Methods**), this convergence of results should only be observable once, at most, in our dataset due to random chance.

### Data Access

Following MINSEQE reporting guidelines, all behavioral data, metadata, raw and processed sequencing data have been made available on NCBI Gene Expression Omnibus (GEO; https://www.ncbi.nlm.nih.gov/geo/, accession numbers: GSE225744 (F_0_) and GSE225746 (F_2_)) to reviewers using a private log-in token, and will be publicly released following acceptance for publication.

## Supporting information

Supplemental Methods, Results, Tables, Figures

## Acknowledgments

This study was supported by NIDA U01DA043098 (HA, JL, AAP), ONR 00014-19-1-2149 (HA), NIH R01GM140287 (PM), the Pritzker Neuropsychiatric Research Consortium (HA, SJW) and the Hope for Depression Research Foundation (HDRF) (HA). We would also like to thank our reviewers, whose thoughtful feedback greatly improved our analyses and manuscript, and whose comments provided the inspiration for many of the integrative RNA-Seq/genotype analyses (*cis*-eQTL database, SMR).

## Author Contributions

EKH – F_2_ dissections, analysis of behavioral data, coordination of RNA extraction and RNA-Seq, analysis of F_0_ & F_2_ RNA-Seq data, interpretation of results, manuscript preparation

MHH - Analysis of F_0_ & F_2_ RNA-Seq data, interpretation of results, manuscript preparation

DBM – Integration of RNA-Seq data with GWAS data (*cis*-eQTL and SMR analyses), manuscript preparation

PB – Coordination of breeding, F_0_ and F_2_ behavioral and tissue collection, analysis of behavioral data

FM - Analysis of F_0_ & F_2_ RNA-Seq data

ASC – Analysis of F_0_ & F_2_ WGS data

ABO – Analysis of F_0_ WGS data

KA – F_2_ dissections

PM – Supervision of integrative RNA-Seq/GWAS analyses

SBF – F_2_ PavCA behavioral assessment guidance

SJW – Project conception

AAP – Project conception, funding, supervision of integrative

RNA-Seq/GWAS analysis

JL - Project conception, funding

HA – Project conception, funding, supervision of experimental design execution and RNA-Seq analysis, manuscript preparation

