## Supplemental Methods, Results, Tables, Figures for "Bioenergetic-Related Gene Expression in the Hippocampus Predicts Internalizing vs. Externalizing Behavior in a F_2_ Cross of Selectively-Bred Rats"

**Table of Contents: Supplemental Tables**

### Table of Contents: Supplemental Figures

### Supplementary Methods

#### Animals

All procedures were conducted in accordance with the guidelines outlined in the National Institutes of Health Guide for the Care and Use of Animals and were approved by the Institutional Animal Care and Use Committee at the University of Michigan (IACUC Animal Protocol Number: PRO00003921, PI: Huda Akil, Title: Drug Abuse and Emotional Reactivity, Approval Date: 8/20/2012, Expiration Date: 8/20/2015).

We established two lines of rats that were selectively bred for a high or low propensity to explore a novel environment at the Michigan Neuroscience Institute, University of Michigan ^1^. These lines were named the “bred High Responders” to novelty (Wakil:bHRs, RRID:RGD_405847397) and “bred Low Responders” to novelty (Wakil:bLRs, RRID:RGD_405847400), respectively. The full procedures for generating the bred lines is detailed in ^1^.

After 37 generations of selective breeding, an initial set (F_0_) of bHRs deriving from 12 distinct families were bred with bLRs from 12 distinct families to create 12 intercross families. The offspring of this intercross (F_1_) were then bred with each other to help the re-emergence/divergence of the phenotypes in the F_2_ generation. To amplify divergence, F_1_ rats that had a lower locomotor response to novelty were bred together, and F_1_ rats that had a higher locomotor response to novelty were bred together (**Fig 3**). The resulting 48 F_2_ litters generated a total of 540 male and female rats, 323 of which were then phenotyped as young adults (2-4 months old: locomotor activity in a novel environment at two months, elevated plus maze (EPM) at three months, Pavlovian Conditioned Approach (PavCA) at four months) and 216 of which were phenotyped as juveniles (around one month old, open field). The subset of animals used in our current RNA-Seq study includes males and females from the F_0_ (n=24, six per phenotype for each sex) and F_2_ cross generations (n=250 young adults, M/F=125/125). This subset of animals overlaps with those used in our previous exome sequencing ^2^ and genome wide association studies ^3^, and is independent from the animals included in our earlier male-only bHR/bLR hippocampal meta-analysis study ^4^.

##### Housing Conditions

Breeding pairs (one male and one female) were housed together for 10 days on a 14:10 hour light/dark cycle (lights on at 0500, lights off at 1900) with *ad libitum* access to water and food (5008, PicoLab Laboratory Rodent Diet, LabDiet), and females monitored for signs of pregnancy. Upon weaning, female and male animals were housed in separate rooms on a 14:10 light:dark cycle (lights on: 0500, lights off at 1900) and 12:12 hour light:dark cycle (lights on at 0700, lights off at 1700), respectively, with *ad libitum* access to water and food (5LOD, PicoLab Laboratory Rodent Diet, LabDiet). The temperature (70–78^o^F) and humidity (30-50%) in the housing rooms were monitored at all times. All experimental animals were marked on their tails with identifying numbers via a non-toxic marker to allow researchers to track the behavior of individual animals. Cages were kept in animal biosafety level-1 housing rooms, and changed once a week, with water bottles changed twice a week. Cage bedding consisted of ¼ inch corn cob bedding (bed-o’cobs, The Andersons, Lab Bedding Products), and animals were housed in pairs or triples with littermates. All experimental animals were housed in the same rooms for all generations tested.

##### Experimental Procedures: Transport, Acclimation, Timing, and Experimenters

As distinct generations, the F_0_ and F_2_ animals were subjected to behavioral testing, sacrifice, and dissection approximately 8 months apart. As much as possible, the testing and sacrifice of the F_0_ rats was run in a manner counterbalanced for F_0_ phenotype (bHR/bLR), but true, full blinding of the experimenters to phenotype was impossible due to the huge magnitude of the behavioral differences between the bred lines. Likewise, during behavioral testing and sacrifice of both the F_0_ and F_2_ rats, the experimenters were unable to be blinded to sex, due to housing in separate colony rooms and obvious morphological differences.

During procedures, cages were transported between housing and behavioral testing rooms using a cart. Experimental animals remained within their housing room until immediately before undergoing behavioral testing and were placed immediately back into their housing room after finishing testing. Behavioral testing was performed in the morning as per established laboratory protocol. For practical reasons it was necessary to test males before females on different days, as any residual scent of the females could dramatically alter male behavior. To reduce disruption, all residents of a cage, which were same-sex siblings, were typically tested on behavioral tasks within the same batch and/or spaced closely together in time. Within each sex, the order that locomotor testing was administered was randomly distributed across cages (of same-sex siblings) and followed a different order than the administration of the Elevated Plus Maze (EPM) task. The specific method used to randomize the order was not recorded.

At sacrifice, experimental animals were placed into a holding room next door to the sacrifice room for acclimation after being moved from their housing room, to decrease the stress effects of being moved through the animal vivarium. Individual animals were carried by hand from the holding room into the sacrifice room, where they were immediately decapitated. Again, for practical reasons, males were sacrificed before females, and all residents of a cage, which were same-sex siblings, were sacrificed closely together in time.

##### Behavioral Analysis

At approximately two months of age, all F_0_ and F_2_ rats were tested for locomotor activity in a novel environment that was the same size as a home cage but located in a different room with novel cues ^1^. The number of beam breaks (counts) was measured over 60 minutes for both lateral and rearing movements, the cumulative total of which were combined to generate the total locomotor score (LocoScore). The locomotor test date, batch, rack, and box were recorded as metadata and have been released along with the behavioral data. For the F_2_ rats, locomotor testing was spread out over a two-week period. Males were tested on earlier dates than females. Siblings housed in the same cage were tested on the same date within the same batch.

Next, all F_0_ and F_2_ rats were evaluated for exploratory and anxiety-like behavior on the EPM. The EPM test began by placing animals in the intersection of the arms which allows access to the two open and the two closed arms. During the five-minute test period, the room was dimly lit (40 lux) and animals’ behavior monitored using a video tracking system (Ethovision, Noldus Information Technology) that recorded the amount of time spent in the arms (converted to percent time of five min), the distance traveled (cm), and the time immobile (sec). The number of fecal boli was counted as an additional measure of emotionality but was not used in our RNA-Seq analysis. The EPM test date and EPM test order were recorded as metadata and have been released along with the behavioral data. For the F_2_ rats, EPM testing was spread out across two weeks. On each test date, males were tested before females. Siblings housed in the same cage were tested on the same date, sequentially.

A subset of F_2_ rats (n=209) were subsequently tested for PavCA behavior as described in ^5^, to determine the re-emergence of the sign tracker (ST) and goal tracker (GT) phenotypes. Three distinct dependent variables, probability difference, response bias, and latency score, were averaged to generate a PavCA Index. The probability difference was defined as the probability of lever contact minus the probability of food magazine entries. Response bias was defined as the total conditioned stimuli (CS: lever) contacts minus the total food magazine entries, divided by the sum of the two behaviors. The latency score was defined as the average latency to enter the food magazine minus the average latency to contact the lever, divided by the length of the CS duration (eight sec). The PavCA Index for the last two days of testing (i.e., days six and seven) was used to determine if animals display ST (values>0.5) or GT (values<-0.5) behavior.

##### Sacrifice and Tissue Dissection

Adult experimental animals (postnatal days 113-132) were decapitated without anesthesia and brains removed within two minutes of death. For the F_0_ cohort (n=24), whole hippocampus was dissected upon sacrifice and flash frozen at -30^o^C before being stored at -80^o^C. F_0_ males and females were sacrificed and dissected on separate days, spaced one month apart, but sacrifice and dissection on each day was counterbalanced for bHR/bLR line. For the F_2_ rats (n=250), whole brains were extracted and flash frozen at time of sacrifice. Rats from the F_2_ litters were all sacrificed within a two-week period, with all males sacrificed during the first week and all females sacrificed during the second week. Later, brains were hemi-sected and four coronal slabs encompassing both the dorsal and ventral hippocampus (-2.12 mm to -6.04 mm Bregma; ^6^) were cut from one half of each brain. Hole punches were taken from both the dorsal and ventral hippocampus in the cryostat (-12^o^C) or on dry ice. For the F_2_ rats, dissections were counter-balanced by hippocampal side, sex, and genetic family.

#### RNA-Seq

##### RNA-Seq Basic Experimental Design

For the F_0_ and F_2_ RNA-Seq experiments, the basic primary outcome measures were normalized gene-level counts within the quality-controlled, filtered, normalized RNA-Seq data (Log2 counts per million – see full description below). For the F_0_ sample, the independent variables of interest were bred line (bLR vs. bHR) and sex. For the F_2_ sample, the independent variables of interest were locomotor activity in a novel environment (LocoScore), distance traveled and time immobile on the EPM, percent time spent in the open arms of the EPM, and PavCA index.

Individual animals were the experimental unit used in our study. The sample sizes for the F_0_ and F_2_ RNA-Seq experiments were determined by animal availability, budget, and logistical capacity. By rough estimation (<https://sample-size.net/> ), within the full F_0_ RNA-Seq sample size (n=24), there was sufficient power (80%) to reliably detect at a traditional alpha (p<0.05) group differences in gene expression between the bred lines (n=12/group) or between the sexes (n=12/sex) with very large effect sizes (Cohen’s d>1.2), or much larger (Cohen’s d>1.8) interactive effects of bred line and sex (n=6/group*line). In contrast, within the full F_2_ RNA-Seq sample size (n=250), there was sufficient power (80%) to reliably detect at a traditional alpha (p<0.05) much smaller effect sizes for relationships between the behavioral variables and gene expression (R>0.176 or R^2^>0.031, 3% of the variance explained). This greater sensitivity to smaller effect sizes was necessary for the F_2_ sample because, as expected, there was a much smaller range in behavior in the F_2_ rats than the differences present between bHR and bLR animals (**Fig 3**, **Fig S4**), as well as many additional sources of technical variation within the F_2_ behavioral and RNA-Seq measurements due to the logistical complexity of managing a large sample collection.

##### RNA Extraction and Sequencing

RNA extractions and RNA-Seq were performed by HudsonAlpha Discovery at Discovery Life Sciences. Nucleotides were extracted from hippocampal tissue using the Qiagen RNeasy Plus Mini Kit. A Ribogreen Assay was performed using 2 μl of a 7X dilution to measure RNA concentration and total RNA amount. To measure RNA quality, 2 μl of various dilutions were loaded onto a HS RNA Fragment Analyzer chip. The samples were then normalized to 500ng input into 50μl volume for use in Poly A RNA-seq library reactions that were completed through Second Strand Synthesis and Purification. Diluted samples were used for a PicoGreen assay and loaded onto a HS Caliper chip for quality analysis of the cDNA libraries. Following Kapa qPCR, samples were sequenced on a NovaSeq S4 101PE flowcell targeting an average of 25 million reads.

##### RNA-Seq Data Analysis: General Methods for Both the F_0_ and F_2_ Datasets

###### Data and Code Availability

The full F_0_ and F_2_ RNA-Seq datasets have been made available on NCBI Gene Expression Omnibus (GEO; <https://www.ncbi.nlm.nih.gov/geo/>, accession numbers GSE225744 (F_0_) and GSE225746 (F_2_)) to reviewers using a private log-in token, and will be publicly released following acceptance for publication. To enable unambiguous interpretation and facilitate reproduction of our results, these data releases each meet Minimum Information about a high-throughput nucleotide SEQuencing Experiment (MINSEQE) reporting guidelines.

The initial RNA-Seq data preprocessing was performed using a standard pipeline. All later downstream analyses were performed in Rstudio (R v.3.4.1-v.4.2.2, R-studio v.1.0.153- v.2022.12.0+353), with full code available at ([*https://github.com/hagenaue/NIDA_bLRvsbHR_F2Cross_HC_RNASeq*](https://github.com/hagenaue/NIDA_bLRvsbHR_F2Cross_HC_RNASeq)) under the MIT license.

###### Alignment and Assembly

RNA-Seq data was aligned using the STAR algorithm (2.7.3a, Encode parameter: unique hit only) to genome assembly Rnor6. Subread version 2.0.0 was used for obtaining gene level counts with the -s 2 setting for the strand-specific Illumina read libraries and with the Ensembl release 103 rat gtf file. Additional annotation was provided using the R package *org.Rn.eg.db* (v. 3.13.0; ^7^), and *AnnotationHub,* which provided more comprehensive annotation but from a slightly earlier build (Rnor6: Ensembl 88; v.2.8.3 ^8^).

###### Quality Control

Principal components (PC) analysis did not identify any extreme outlier samples during the initial preprocessing. Additional quality control and normalization was performed in Rstudio (v.1.4.1717, R v. 4.1.1). Due to the complexity of the datasets, the analyses that followed preprocessing were updated several times to improve quality control and to better control for technical variation as the analysts gained a better understanding of the experimental design and data structure. Although this process of reanalysis introduced the possibility for bias, analysis decisions were governed by the goal to identify and control for the most impactful sources of technical variation within the dataset, as defined by metrics related to the dataset as a whole (*e.g.,* principal components analysis) and not the results of individual genes of interest. The final analysis methodology is overviewed below.

HudsonAlpha Discovery provided a variety of metrics for evaluating sample RNA quality, including RNA concentration (RNAconc), RNA integrity number (RIN), the percentage of fragments >200 nucleotides (DV200), percent ribosomal RNA (%rRNA), percent messenger RNA (%mRNA), percent coding RNA (%Coding), percent untranslated region (%UTR), percent intronic (%Intronic), percent intergenic (%Intergenic), median coefficient of variation of coverage (Median_CV_Coverage), the overall number of reads (total number of lines in fastq file divided by 4), and the number of reads that passed in-house filtering (PF.Reads). The distributions of these variables, as well as our own calculated final library size, were evaluated for extreme outliers visually using boxplots and violin plots (**Fig S1**). Outlier exclusion was determined using highly conservative filters (*e.g.,* more than a full distribution beyond the full data distribution for a variable) and focused on variables traditionally used to screen RNA-Seq samples (RIN, RNAconc, library size). The analysts chose to pursue a conservative approach to outlier exclusion to preserve statistical power, and because particularly impactful technical co-variates could be later included in our final differential expression model.

The count matrix was filtered to remove transcripts with low-level expression (<1 read in 75% of the subjects). To correct for artifacts introduced by sample-level differences in RNA production, normalization factors were calculated using the trimmed mean of M-values (TMM) method ^9^ within the *calcNormFactors()* function (package: edgeR, v.3.34.1; ^10^) and then the data was transformed to Log2 counts per million (Log2 cpm, ^11^). To screen for sample mislabeling, sample sex was compared to sex chromosome gene expression (i.e., *Eif2s3y, Ddx3*). Principal components analysis was then run again on the cleaned log2 CPM data in preparation for model selection.

###### Model Selection

To determine which technical variables might be important covariates to include in the differential expression model, we first evaluated the quality-controlled datasets for confounding collinearity between variables of interest (*i.e.,* sex, bHR/bLR lineage, LocoScore, percent time in the EPM open arms, total EPM distance traveled, EPM time immobile, PavCA Index, ST/GT behavior) and technical variables using either Fisher’s exact test (categorical vs. categorical), linear regression (numeric vs. categorical), or one-way ANOVA (numeric vs. categorical). Technical variables were then further evaluated for potential impact on differential expression results by examining their relationship with the top PCs of variation in the data (PC1-3).

###### Differential Expression Analysis

Differential expression was calculated using the limma/voom method (^12^, package: *limma* v.3.48.3) with observed precision weights in a weighted least squares linear regression. Contrasts were defined by treatment, with the intercept (reference group) for Lineage set as bHR and for Sex set as Male. Standard error was then moderated towards a global value provided by an empirical Bayes distribution (function *eBayes())* and multiple comparison correction was performed using the Benjamini-Hochberg method (false discovery rate (FDR) or q-value).

##### F_0_ RNA-Seq Dataset Analysis Methods

###### Quality Control

One sample from a female bHR rat was removed due to low RNA integrity (RIN=6.3, all other RINs>8.0), leaving a final sample size of n=23 subjects (n=5 female bHRs, n=6 for each of the other subgroups (male bHRs, female bLRs, male bLRs)). Library size was found to vary from 16.8 million to 28.4 million (median=22.3 million). The initial count matrix included reads aligned to 14,802 Ensembl-annotated transcripts (rows with reads>0). After filtering to remove transcripts with low-level expression (<1 read in 75% of the subjects), data from 13,786 transcripts remained.

###### Model Selection

Since the F_0_ dataset was relatively small (n=23 following quality control), we felt that the differential expression analysis could handle, at most, three covariates. Since the F_2_ dataset had a much larger sample size (n=245 following quality control), we used the results from the F_2_ samples to help inform model choice. Within both datasets, %rRNA and %Intergenic were strongly related to the top two PCs of variation (F_0_: PC1 vs. %rRNA (R=-0.83, p=7.86E-07), PC1 vs. %Intergenic (R=0.43, p=0.0387), PC2 vs. %Intergenic (R=-0.74, p=5.66E-05)) and at least partially independent of each other (R=-0.02) (**Fig S2**). Therefore, for our final model we ended up prioritizing %rRNA and %Intergenic over other potential covariates with slightly stronger relationships with PC1 and PC2 in only the F_0_ dataset:

*Model 1 (“M1: Main Effects Model”):*

$$y\sim\beta_{0}+\beta_{1}Lineage+\beta_{2}Sex+\beta_{3}\%rRNA+\beta_{4}\%Intergenic+\varepsilon$$

When exploring the possibility of a lineage*sex interaction, we reduced the covariates in the model to only %rRNA to maintain statistical power:

*Model 2 (“M2: Interaction Model”):*

$$y\sim\beta_{0}+\beta_{1}Lineage+\beta_{2}Sex+\beta_{3}\%rRNA+\beta_{5}Lineage*Sex+\varepsilon$$

Following these analyses, we explored the results of reduced models lacking covariates to determine the sensitivity of our findings to model specification. All models produced a very similar list of top genes.

##### F_2_ RNA-Seq Dataset Analysis Methods

###### Quality Control

The RNA-Sequencing IDs fell into three distinct sequential clusters, which we deemed RNA-Sequencing batches. Library size varied from 17.9 million to 109.7 million (median=21.7 million). The sample with the unusually large library size (109.7 million, all other library sizes <40 million) was from a male rat and was discarded. Two more male samples were removed due to low RINs (RINs=6.6 and 7.5, all other RINs>8.3), and two female samples were removed because the labeled sex was incongruous with sex-specific gene expression (*Eif2s3y, Ddx3*), leaving a final sample size of n=245 subjects (n=122 males, n=123 females). Later, we also removed the distance traveled measurement for one subject with an unusually high distance traveled (5209 cm, next lowest was <3500 cm), leaving a sample size of n=244 for that variable.

The initial count matrix included reads aligned to 15,510 Ensembl-annotated transcripts (rows with reads>0). After filtering to remove transcripts with low-level expression (<1 read in a quarter (62) of the subjects), data from 14,056 transcripts remained.

###### Model Selection

Since strong relationships were identified between batch-related variables (i.e., dissector, sequencing batch, hippocampal dissection date, experience with the PavCA task (“STGT_Experience”)) and many of the mRNA metrics (**Fig S2, Fig S3**), we chose a model that included the two RNA metrics that were most strongly related to both PC1 and PC2 in both the F_0_ and F_2_ datasets (i.e., %Intergenic and %rRNA: PC1 vs. %rRNA: R=0.87, p<2.2E-16; PC1 vs. %Intergenic: R=-0.95, p=2.65E-122; PC2 vs. %rRNA: R=0.16, p=0.010680) as well as the most impactful batch variables in the F_2_ dataset:

*Model 3 (“M3: Simple, Similar to F_0_ Model”):*

$$y\sim\beta_{0}+\beta_{1}VariableOfInterest+\beta_{2}Sex+\beta_{3}\%rRNA+\beta_{4}\%Intergenic+\beta_{5}Dissector+\beta_{6}STGT\_Experience +\beta_{7-8}SequencingBatch +\varepsilon$$

We also tested two expanded models that included batch variables and five RNA metrics that were the most strongly related to top PCs (PC1-PC8) in the F_2_ dataset (i.e., %rRNA, %Intergenic, %Intronic, %UTR, RNAconc) within an automated model selection procedure using the *step()* function from the R package *stats* (using BIC as benchmark (k=log(245), direction=both). Despite these additional efforts to control for potential sources of technical variation from the data, the results remained noisy, and there weren’t any differentially expressed genes identified for our variables of interest that survived a traditional false discovery rate correction (False Discovery Rate (FDR)<0.10). We also tested whether cleaning the behavioral variables of potential batch and order effects (*LocoScore*: test date, batch, rack, box; *EPM*: test date, test order) prior to differential expression analysis tightened the results, but did not find any notable effect on outcome. Therefore, moving forward we focused on the differential expression results from M3 for comparisons with our other findings, since it included similar covariates to the F_0_ dataset.

##### Comparison of F_0_, F_2_, and bHR/bLR Meta-Analysis RNA-Seq Results

To compare the F_0_ results (which were taken from the F37 generation of breeding) to the results from our previous meta-analysis of bHR/bLR hippocampal RNA-Seq studies from late-generations of breeding (F37, F43) ^4^, we first removed all rows for which an ENSEMBL gene mapped to more than one gene symbol. We then joined these F_0_ results to our previous meta-analysis results by official gene symbol. To compare the F_0_ RNA-Seq and bHR/bLR meta-analysis results to the F_2_ RNA-Seq results, this data-frame was then further joined with the F_2_ results by ENSEMBL ID, with 13,339 ENSEMBL IDs present in both.

For downstream analyses, we defined bHR/bLR differentially expressed genes as being the 1,063 genes associated with bHR/bLR lineage with FDR<0.10 in either the F_0_ dataset or meta-analysis or nominal replication p<0.05 in both with consistent direction of effect (**Fig 5C-E**).

##### Gene Set Enrichment Analysis

Gene Set Enrichment Analysis (GSEA) was performed to elucidate functional trends in the datasets. To create a continuous variable representing the differential expression associated with bLR-like phenotype versus bHR-like phenotype, we averaged the t-statistics from the bLR vs. bHR differential expression identified in the F_0_ dataset and former meta-analysis ^4^, and each behavior in the F_2_ dataset (i.e., LocoScore, EPM distance traveled, EPM time immobile, percent time in EPM open arms, PavCA index). Before averaging, each comparison was set so that upregulation reflected the bLR-like phenotype, therefore the t-statistics from LocoScore, EPM distance traveled, percent time in EPM open arms, and PavCA index needed to be inverted (multiplied by -1). Prior to use in the GSEA, we filtered the average t-statistics to remove ENSEMBL genes that lacked gene symbol annotation and averaged the values when more than one ENSEMBL gene was annotated with the same gene symbol (22 instances), resulting in the representation of 12,695 gene symbols.

This average t-statistic was then used as the input for GSEA (fgsea package v.1.2.1 ^13^, nperm=10,000, minSize=10, maxSize=1000, FDR<0.05), using a custom gene set database that included standard gene ontology as well as brain-specific cell types, regional signatures, and differential expression results from public databases (Brain.GMT v.1 for rats, ^14^ ). A second non-directional analysis was run using the absolute value of the average t-statistic.

#### Constructing a Hippocampal *cis*-eQTL Database

To construct a hippocampal *cis-*eQTL database, we used pre-processed genotype and RNA-Seq data from the F_2_ rats. For the genotype data, we used the final quality-controlled results from the GWAS experiment performed by ^3^ (data release: <https://doi.org/10.6075/J0K074G9>, n=4,425,349 SNPs) and limited the analysis to the autosomes. For the hippocampal RNA-Seq data, we used the final quality-controlled Log2 CPM data from the F_2_ rats (n=245), which contained data from 13,547 autosomal genes. The Log2 CPM data was then corrected for the batch effects and other technical co-variates included in our differential expression statistical model (*i.e.,* residualized), and then further normalized using applied rank-based inverse normal transformation to make the eQTL analysis more robust to outlier data points ^15,16^.

We performed *cis*-eQTL mapping using single-SNP linear regression implemented in *tensorQTL (v*.*1.0.6,* ^17^). We tested SNPs with minor allele frequency (MAF) >0.01 in *cis*-windows up to 1 Mb upstream and downstream of each gene's transcription start site (TSS). To control for potential population stratification, principal components analysis (PCA) was run on the genotype matrix of alternative allele counts, following pruning for linkage disequilibrium in *Plink2 v.2.00a2.3* (^18^, filter: MAF>0.05) using the *indep-pairwise* function (*parameters*: window size: 200 kb, variant count step size: 100, unphased-hardcall-r^2 threshold: 0.1; *final matrix size*: 28,763 SNPs). PCA was also run on the gene expression matrix to identify potential latent sources of unwanted variation. The first five principal components of variation from each of these PCA analyses were included as covariates within the eQTL regression model.

The results from this initial +/-1 MB scan for each gene were then adjusted using a permutation-based procedure within *tensorQTL* to control for the variable number of SNPs present within +/-1 MB of the TSS for each gene, their correlation structure (linkage disequilibrium or LD), and overall distribution of allele frequencies. Within this procedure, a null distribution was constructed by repeating the +/-1MB *cis*-window scan using random permutations of the expression values for the gene of interest, storing the p-values for the strongest relationship each time. An empirical beta distribution was then fit to this null distribution, allowing more precise estimates for small adjusted p-values without heavy computational burden.

The permutation-based p-value for the top SNP for each gene was then corrected for false FDR using the permutation-based p-values from the top SNP for all genes, with a significant eVariant-eGene relationship (*cis*-eQTL) defined by FDR<0.05. In instances where multiple top SNPs in perfect LD were associated with the same gene, a single SNP was selected randomly for downstream analyses and visualization. We then tested for additional, conditionally independent *cis*-eQTLs for each eGene using tensorQTL's built-in stepwise regression procedure using default settings.

We calculated *cis*-eQTL effect size as allelic fold change (aFC). We used the package *aFC.py* to estimate aFC from the raw expression read counts based on an additive *cis*-regulatory model, including the same covariates as for eQTL mapping ^19^.

##### Predicting bHR/bLR Differential Expression Using the *cis*-eQTL Database

We used the *cis*-eQTL database to predict the effect of genetic variation that segregates the bHR/bLR lines on gene expression. We extracted genotype information for each eVariant for the 20 F_0_ rats (n=10 bHR/n=10 bLR) sequenced in ^3^ using the publicly released *.vcf* files (https://doi.org/10.6075/J0K074G9) and package *VcfR* (v1.14.0, ^20^, *https://cran.r-project.org/web/packages/vcfR/vcfR.pdf*). We then ran a population differentiation analysis using the function *myDiff()* to calculate the parameter G_st’_ ^21^, which ranges from 0 (no bHR/bLR segregation of alleles) to 1 (full bHR/bLR segregation of alleles). Within our sample size (n=20), we found that variants with partial bHR/bLR segregation, in which all subjects from a phenotype (*e.g.,* bHRs) had 2 reference alleles (0/0) and all subjects from the other phenotype had one reference allele and one alternate allele (0/1), had a G_st’_ of 0.2703, and decided to use this as a meaningful effect size threshold for downstream analyses. Using this threshold, 42.02% of the eVariants were partially segregated in bHR/bLR (G_st’_>0.27: 2452/5836 unique eVariants), whereas only 0.39% were fully segregated (G_st’_=1: 23/5836 of unique eVariants).

To predict the effect of the partially and fully segregated eVariants on gene expression, we determined which allele was more frequent in the bLR vs. bHR F_0_ rats, and then flipped the sign of the Log2 aFC to reflect the bLR-enriched allele vs. bHR-enriched allele (instead of alternative allele vs. reference allele). These bLR/bHR Log2 aFC predictions were then compared to the F_0_ differential expression results (Log2FC, 2500 eGene/eVariant combinations) using both parametric (linear regression) and non-parametric (Spearman’s rho) methods. As additional validation, a similar analysis was run comparing bLR/bHR Log2 aFC predictions to the bHR/bLR late generation RNA-Seq meta-analysis results (estimated *d*).

##### Examining the Overlap of Our *cis*-eQTL Results with RatGTEx

As an exploratory analysis, we examined the similarity between our *cis*-eQTL results and those from previous rat brain eQTL studies found in RatGTEx (<https://ratgtex.org/>). RatGTEx includes eQTLs from 12 tissues, 9 of which are brain-related, derived from 5 cohorts of heterogeneous stock rats (^15^, (n=75-81): Infralimbic cortex, lateral habenula, nucleus accumbens core, orbitofrontal cortex, prelimbic cortex; Mitchell & Hitzemann, *unpublished,* (n=185-191): basolateral amygdala, prefrontal cortex, nucleus accumbens; Telese et al., *unpublished,* (n=339)*:* brain hemisphere; ^22^, (n=51): Eye; ^23^, (n=401-411): adipose tissue, liver). Most of our eGenes (4786 of 5351 unique eGenes) were present in the RatGTEx eQTL database (database downloaded 10-25-2023 from https://ratgtex.org/data/eqtl/rn6.eqtls_indep.txt; 16,555 unique eGenes total, 11,365 of which are present in our full F_2_ gene expression dataset). However, our analyses typically did not converge upon the same top eVariants as the other RatGTEx studies (overlap: 115 of our 5836 unique eVariants, RatGTEx: 61,146 eVariants total). This lack of overlap is likely to derive, in part, due to the difficulty of pinpointing the exact relevant eVariant within linkage disequilibrium blocks. Therefore, we expanded our analysis to include all nominally-significant eVariants for each of our 5351 eGenes (p<0.05, as defined by the permutation-corrected p-values): 7,930,908 eQTLs (eVariant/eGene combinations) representing 2,492,126 eVariants. Of these, 10,321 *cis*-eQTLs (eGene/eVariant combinations) were also found in RatGTEx, representing 9,843 unique eVariants and 2,885 unique eGenes. We then compared the slopes of the *cis*-eQTLs from the hippocampus to the slopes from each of the RatGTEx regions using linear regression (following outlier removal, defined by abs(slope)>5) or using non-parametric methods (no outlier removal, Spearman’s rho). We decided to prioritize comparisons on the RatGTEx *cis*-eQTL databases that included tissue most akin to the hippocampus (derived from the cortical plate): brain hemisphere, basolateral amygdala, prelimbic cortex, infralimbic cortex, orbitofrontal cortex. Then, as a comparison point, we repeated the same analyses using *cis*-eQTLs derived from non-cortical brain regions (nucleus accumbens, lateral habenula) and non-brain tissue (adipose tissue, liver). We did not use the *cis*-eQTLs derived from eye tissue in RatGTEx for this comparison because there were much fewer of them (567 cis-eQTLs total, only 118 of which were shared with the hippocampal *cis*-eQTL database), most likely due to small sample size (n=51).

##### Co-localization of *cis*-eQTLs with Regions of the Genome Associated with bHR/bLR-like Behavior (QTLs)

We used Summary Data-based Mendelian Randomization (SMR; ^24^) to test for colocalization of our *cis*-eQTLs with QTLs for bHR/bLR-like behaviors from our previously published GWAS using the full F_2_ adult and juvenile cohorts. We focused on the QTLs for the F_2_ adult behaviors included in our differential expression analysis (LocoScore, EPM time immobile, EPM distance traveled, EPM % time in the open arms, PavCA Index), and juvenile behaviors targeting analogous traits (OF time immobile, OF distance traveled, OF % time in center). We selected the top eVariant per *cis*-eQTL, and then computed *z*-scores for the *cis*-eQTL and GWAS associations for the eVariant by dividing the slope by its standard error. We computed the approximate chi-squared test statistic using the SMR method, and then calculated a *P*-value using the upper tail of the chi-squared distribution with one degree of freedom. A FDR correction was applied using the *mt.rawp2adjp()* function (proc=”BH”) in the *multtest* package (v.2.26.0, ^25^). Results were visualized using the *manhattan()* plot function in the *qqman* package (v.0.1.8, ^26^).

To narrow down our pool of top candidate genes for mediating the effect of genetic variation on behavior, we used a convergence of information from our different samples and analyses. First, since we are hypothesizing that an eVariant located within a region of the genome implicated in behavior (QTL) is driving hippocampal differential expression that is mediating the effect of genetic variation on behavior, we tested whether the SMR t-statistic, which represents the strength of eVariant/QTL co-localization, correlated with F_2_ behavioral differential expression. Within this analysis, we also assigned a predicted direction of effect for the relationship between gene expression and behavior using the direction of effect of the eVariant on gene expression in the *cis*-eQTL analysis and the direction of effect of the eVariant on behavior in the QTL analysis. For example, if the alternate allele was associated with congruent effects on behavior and gene expression (*i.e.,* an increase in F_2_ behavior (GWAS Z-score>0) with an increase in gene expression (Log2 aFC>0) or a decrease in F_2_ behavior (GWAS Z-score<0) with a decrease in gene expression (Log2 aFC<0)), then the association between gene expression and behavior was predicted to be positive. If the alternate allele was associated with incongruent effects on behavior and gene expression (*i.e.,* an increase in F_2_ behavior (GWAS Z-score>0) with a decrease in gene expression (Log2 aFC<0) or a decrease in F_2_ behavior (GWAS Z-score<0) with an increase in gene expression (Log2 aFC>0)), then the association between gene expression and behavior was predicted to be negative.

The most compelling candidates for mediating the effect of selective breeding on behavior should have expression strongly related to genetic variation (*cis*-eQTLs) that is segregated in bHR/bLR (G_st’_>0.27) in a manner that correctly predicts bHR/bLR differential expression and co-localized with a QTL for behavior in a manner that correctly predicts F_2_ differential expression associated with that behavior. Therefore, we next examined the correlation between the “directional” SMR T-statistics and F_2_ Log2FCs for each behavior, both in the full dataset (all 5937 cis-eQTLs) and also within the subset of *cis*-eQTLs that we had already confirmed were segregated in bHR/bLRs with strong differential expression in bHR/bLRs matching what would be predicted based on the distribution of alleles in the two lines (492 cis-eQTLs representing 456 eGenes). We also examined the correlation between the “directional” SMR T-statistics for the F_2_ juvenile behaviors and the Log2FC for their analogous behaviors in the F_2_ adults (juvenile distance traveled in the open field vs. adult distance traveled on the EPM, juvenile time immobile in the open field vs. adult time immobile on the EPM, juvenile % time in the center of the open field vs. adult % time in the open arms of the EPM). For our final top candidate genes, we chose the *cis*-eQTLs that were significantly co-localized (FDR<0.10) with QTLs for an adult or juvenile F_2_ behavior that had a predicted direction of effect (directional SMR T-statistic) that matched what was observed for the relationship between gene expression and the same (or analogous) behavior in the F_2_ differential expression results (p<0.05).

##### Calculation of the Risk of Observing Convergent Results Due to Random Chance

Given the number of statistical results feeding into our final choice of top candidate genes – often from correlated information sources – we double-checked the probability of identifying convergence by false discovery. To do this, we took the most conservative route possible: we examined the probability of a gene in our dataset qualifying for each criterion using the percent of genes meeting that criterion in each analysis (*i.e.,* making no assumptions regarding whether each individual result is a true positive or true negative, and simply examining the likelihood of the positive results ***converging across analyses*** due to random chance). Here is a description of our calculations and logic:

**bHR/bLR differentially expressed genes:** 1,045 genes in the dataset met our criteria for being a bHR/bLR differentially expressed gene, and 13,339 genes were in our final analysis (present in both the F_0_ and F_2_ datasets). Therefore, there was a 7.8% chance (1,045/13,339) that a gene in our final analysis was defined as a bHR/bLR differentially expressed gene by our criteria.

**F_2_ behavioral differentially expressed genes:** 1,999 genes in the dataset met our loose criteria for having bHR-like or bLR-like differential expression (p<0.05) in the F_2_ dataset. Therefore, there was a 15.0% chance (1,999/13,339) that a gene in our final analysis was defined as a bHR-like or bLR-like behavioral differentially expressed gene by our criteria.

**bHR/bLR segregated *cis*-eQTLs:** 2,500 *cis*-eQTLs (representing 2395 eGenes) met our criteria for having bHR/bLR segregated eVariants. Therefore, there was an 18.0% chance (2395/13,339) that a gene in our final analysis was defined as having a bHR/bLR segregated *cis*-eQTL by our criteria.

**The probability of identifying convergence due to random chance using the first 3 criteria:** To calculate the probability of these three sets of results converging due to random chance, we calculated two probabilities:

1. The probability that all three results meet criterion and all three results show “bLR-like” upregulation (or predicted “bLR-like” upregulation for the bHR/bLR-segregated cis-eQTLs). For the sake of efficient calculation, we treated the probability of a gene being upregulated as being 50% in each analysis:

(0.078*0.5)*(0.15*0.5)*(0.18*0.5)=0.00026 or 0.026% chance

2. The probability that all three results meet criterion and all three results show “bHR-like” upregulation (or predicted “bHR-like” upregulation for the bHR/bLR-segregated cis-eQTLs). For the sake of efficient calculation, we treated the probability of a gene being upregulated as being 50% in each analysis:

(0.078*0.5)*(0.15*0.5)*(0.18*0.5)=0.00026 or 0.026% chance

These two possibilities are mutually exclusive. Therefore, the probability of a gene having results that meet either of these two forms of convergence is:

0.00026+0.00026=0.00052 or 0.052% chance.

Therefore, within a dataset with 13,339 genes, we would expect to see this convergence 6.9 times (i.e., basically 7 genes) due to random chance. In our study, we had 104 genes that met these criteria, or 15X what would be expected by random chance (104/6.9).

**Probability of identifying convergence with the SMR co-localization results:** We next examined the probability of convergence with the SMR co-localization results under conditions of random chance. Since the SMR analysis was only run with significant *cis*-eQTLs, the probability of a gene meeting this criterion is conditional on the result already being identified a *cis*-eQTL: 80 genes met criteria for having *cis*-eQTLs that were co-localized with behavioral QTLs (FDR<0.10), and there were 5351 genes that had *cis*-eQTLs. Therefore, there was a 1.50% chance (80/5351) of a gene meeting our criterion for having a *cis*-eQTL co-localized with a behavioral QTL if it had already been identified as a *cis*-eQTL.

To be considered a final result, we also required that the co-localization of the *cis*-eQTL with the behavioral QTL correctly predict the direction of effect in the F_2_ differential expression results for that behavior (or for the adult behavior analogous to the behavior in juveniles). However, since the SMR co-localization results were identified using both *cis*-eQTLs and QTLs derived from a sample of F_2_ animals overlapping that used for the F_2_ differential expression results, it is inaccurate to treat these results as independent events.

Therefore, if we take the most conservative route, and use in our calculations the probability of a gene that has already met our initial 3 criteria (0.00052 or 0.052% chance – see rationale above) also being co-localized with a behavioral QTL in a manner that correctly predicts the direction of effect in the F_2_ differential expression results (15.4% or 16/104 in our results), then we would predict that the potential for false discovery of convergence in our dataset would be:

0.00052*0.154=0.000080 or 0.008%.

Therefore, within a dataset with 13,339 genes, we would expect to see this convergence 1.07 times (*i.e.,* 1 gene) due to random chance. Since this is an overly conservative estimate, we can confidently argue that <1 of our 16 final top candidate genes is likely to represent a convergence of supporting results due to random chance.

##### Calculation of the Risk of Observing Convergent Results with Other Rat Models Due to Random Chance:

Similar to the strategy discussed above, we double-checked the probability of identifying convergence with other rat models by false discovery using the most conservative route possible: we examined the probability of a gene in our dataset qualifying for each criterion using the percent of genes meeting that criterion in each analysis (*i.e.,* making no assumptions regarding whether each individual result is a true positive or true negative, and simply examining the likelihood of the positive results ***converging across analyses*** due to random chance). Here is a description of our calculations and logic:

As described above, there was a 7.8% chance (1,045/13,339) that a gene in our final analysis was defined as a **bHR/bLR differentially expressed gene** by our criterion and a 15.0% chance (1,999/13,339) that a gene in our final analysis was defined as a **bHR-like or bLR-like behavioral differentially expressed gene** by our criteria.

**Differentially expressed genes in other rat models:** In our earlier publication ^4^, we created a database of 2,581 genes that had been previously identified as differentially expressed in the hippocampus of other bLR-like and bHR-like rat models targeting hereditary behavioral traits reflecting extremes on the internalizing/externalizing spectrum.

Within our final dataset, there were 849 genes that had been identified as upregulated in other rat models targeting internalizing-like behavior, with 55 of those genes showing upregulation in two other models, and two genes showing upregulation in three other models. Therefore, in our final dataset there was a 6.4% chance (849/13,339), 0.41% chance (55/13,339), and 0.015% chance (2/13,339) of a gene having been documented in the database as previously upregulated in at least one, two, and three models, respectively.

Within our final dataset, there were also 1,222 genes that had been identified as down-regulated in other rat models targeting internalizing-like behavior, with 80 of those genes showing down-regulation in two other rat models, and three genes showing down-regulation in three other models. Therefore, in our final dataset, there was a 9.2% chance (1222/13,339), 0.60% chance (80/13,339), and 0.022% chance (3/13,339) of a gene having been documented in the database as previously down-regulated in at least one, two, and three models, respectively.

**The probability of identifying convergence with other rat models due to random chance using these criteria:** To calculate the probability of our results converging with those of other rat models due to random chance, we calculated a set of probabilities representing convergence with upregulation in other rat models and a set of probabilities representing convergence with downregulation in other rat models:

***Convergence with upregulation in other rat models:***

1. A result is upregulated in bLRs and in association with bLR-like behavior, and is found to be upregulated in at least one other rat model of internalizing behavior:

(0.078*0.5)*(0.15*0.5)*(0.064)= 0.0001872 or 0.019% chance

In our final dataset with 13,339 genes, you would expect to see 2.5 genes showing this convergence due to random chance.

2. A result is upregulated in bLRs and in association with bLR-like behavior, and is found to be upregulated in at least two other rat models of internalizing behavior:

(0.078*0.5)*(0.15*0.5)*(0.0041)= 1.20e-05 or 0.0012% chance

In our final dataset with 13,339 genes, you would expect to see 0.16 genes (*i.e.* no genes) showing this convergence due to random chance.

3. A result is upregulated in bLRs and in association with bLR-like behavior, and is found to be upregulated in three other rat models of internalizing behavior:

(0.078*0.5)*(0.15*0.5)*(0.00015)= 4.39e-07 or 4.39e-05% chance

In our final dataset with 13,339 genes, you would expect to see 0.0059 genes (*i.e.,* no genes) showing this convergence due to random chance.

***Convergence with downregulation in other rat models:***

1. A result is upregulated in bHRs and in association with bHR-like behavior, and is found to be downregulated in at least one other rat model of internalizing behavior:

(0.078*0.5)*(0.15*0.5)*(0.092)= 0.000269 or 0.027% chance

In our final dataset with 13,339 genes, you would expect to see 3.6 genes showing this convergence due to random chance.

2. A result is upregulated in bHRs and in association with bHR-like behavior, and is found to be downregulated in at least two other rat models of internalizing behavior:

(0.078*0.5)*(0.15*0.5)*(0.0060)=1.76e-05 or 0.00176% chance

In our final dataset with 13,339 genes, you would expect to see 0.23 genes (*i.e.,* no genes) showing this convergence due to random chance.

3. A result is upregulated in bHRs and in association with bHR-like behavior, and is found to be downregulated in three other rat models of internalizing behavior:

(0.078*0.5)*(0.15*0.5)*(0.00022)= 6.44e-07 or 4.44e-05% chance.

In our final dataset with 13,339 genes, you would expect to see 0.0086 genes (*i.e.,* no genes) showing this convergence due to random chance.

Having the convergence be all upregulation or all down-regulation are possibilities that are mutually exclusive. Therefore, the probability of a gene having a convergence of results showing upregulation in bLRs, with bLR-like behavior in the F_2_s, and upregulation in at least two other rat models of internalizing like behavior *or* showing upregulation in bHRs, with bHR-like behavior in the F_2_s, and down-regulation in at least two other rat models of internalizing like behavior would be:

1.20e-05+1.76e-05=2.96e-05 or 0.00296% chance.

In our final dataset with 13,339 genes, you would expect to see 0.39 genes (*i.e.,* no genes) showing this convergence due to random chance. Since this is an overly conservative estimate, we can confidently argue that it is likely that none of the six genes that we observed showing a convergence of differential expression results with at least two other rat models are likely to be due to random chance.

### Supplementary Results

#### Constructing a Hippocampal cis-eQTL Database to Determine which Differential Expression is Most Likely to Be Driven Directly by Genetic Variation

To determine which differential expression was most likely to be driven directly by genetic variation, we integrated our current F_2_ transcriptional profiling data (n=245) with our previous whole genome sequencing results (n=4,425,349 SNPs, ^3^) to identify genes with hippocampal expression tightly correlated with nearby genetic variation (*cis*-eQTLs). This integrative analysis proved to be more highly powered than the F_2_ differential expression analysis, most likely due to less noise in the genotype data compared to behavior and identified 5,351 genes (eGenes) with expression that correlated strongly (FDR<0.05) with genetic variation within +/-1 MB of their transcription start sites (TSS). Using stepwise regression, we identified additional conditionally-independent *cis*-eQTLs beyond the strongest *cis*-eQTL for each eGene, and found that 523 of the eGenes had two eQTLs, 27 eGenes had three eQTLs, and three eGenes had four eQTLs (**Fig S5A)**. Altogether, we identified a final total of 5,937 *cis*-eQTLs representing 5,836 unique eVariants. Similar to previous *cis*-eQTL analyses, these eVariants were predominantly located within +/-400 kB of the TSS of their respective eGene (**Fig S5B**). As our hippocampal *cis*-eQTL database represents a valuable resource for the interpretation of rat genomic results, we have shared it on RatGTEx for use by other researchers (<https://ratgtex.org/download/study-data/#HPC_F2>).

##### Comparing the Hippocampal *cis*-eQTL Database to RatGTEx

As an exploratory analysis, we compared our hippocampal *cis*-eQTL database to existing rat *cis*-eQTL databases in RatGTEx. RatGTEx includes eQTLs from 12 tissues, nine of which are brain-related, derived from five cohorts of heterogeneous stock rats (RatGTEX: ^15,22,23^; Mitchell & Hitzemann, *unpublished*, Telese *unpublished),* **Fig S5C).** We found that most of our hippocampal eGenes (4,786 of 5,351 unique eGenes) were present within the RatGTEx eQTL database (database downloaded 10-25-2023; 16,555 unique eGenes total, 11,365 of which are present in our full F_2_ gene expression dataset), and the majority were significant eGenes within at least four other tissues (out of 11 tissues characterized, **Fig S5D**).

Our analyses for the most part, however, did not converge upon the same top eVariants as the other RatGTEx studies (overlap: 115 of our 5,836 unique eVariants, RatGTEx: 61,146 eVariants total). This lack of overlap is likely to derive, in part, due to the difficulty of pinpointing the exact relevant eVariant within linkage disequilibrium blocks. Therefore, we expanded our analysis to include all nominally-significant eVariants for each of our 5,351 eGenes (p<0.05): 7,930,908 eQTLs (eVariant/eGene combinations) representing 2,492,126 eVariants. Of these,10,321 eQTLs (eGene/eVariant combinations) were also found in RatGTEx, representing 9,843 unique eVariants and 2,885 unique eGenes. We then compared the slopes of the *cis*-eQTLs from the hippocampus to the slopes from each of the RatGTEx regions.

The *cis*-eQTLs identified in RatGTEx tended to have a similar slope as what we observed in the hippocampus (**Fig S7-FigS8**). This was particularly true for brain tissue, where correlations ranged from R=0.65-0.75 (all p<2.2e-16). We found that 1,855 *cis*-eQTLs identified using rat brain hemispheres (representing 1,556 eGenes) were at least nominally detectable in the hippocampus, with many showing a similar slope (*without outlier removal:* Spearman’s rho=0.628, *with outlier removal:* R=0.65, R^2^=0.391, beta+/-SE=0.902+/-0.0264, T(1819)=34.157, p<2.2e-16, 34 *cis-*eQTLs (from five eGenes) removed as extreme outliers). The correlations with *cis*-eQTLs identified from individual brain regions were even stronger. Within another subcortical region with cortical-like structure, the basolateral amygdala, 1,063 *cis*-eQTLs (representing 969 eGenes) were identified that were nominally detectable in the hippocampus, with many showing a similar slope (*without outlier removal:* Spearman’s rho=0.752, *with outlier removal:* R=0.72, R^2^=0.53, beta+/-SE=0.81+/-0.0235, T(1053)=34.56, p<2.2e-16, eight *cis-*eQTLs (from one eGene) removed as extreme outliers). For *cis*-eQTLs identified within cortical structures, this was also true (*Infralimbic cortex (IL):* 726 shared *cis-*eQTLs representing 713 eGenes, Spearman’s rho=0.692, R=0.703, R^2^=0.494, beta+/-SE=0.773+/-0.0290, T(727)=26.63, p<2.2e-16; *Orbitofrontal cortex (OFC):* 749 shared *cis-*eQTLs representing 733 eGenes, Spearman’s rho=0.704, R=0.722, R^2^=0.521, beta+/-SE=0.778+/-0.0273, T(747)=28.523, p<2.2e-16; *Prelimbic cortex (PL, from Munro et al. cohort):* 757 shared *cis-*eQTLs representing 739 eGenes, Spearman’s rho=0.694, R=0.69, R^2^=0.481, beta+/-SE=0.778+/-0.0294, T(755)=26.465, p<2.2e-16; *Prelimbic cortex* *(PL2,* from Mitchell & Hitzemann cohort): 1,108 shared *cis-*eQTLs representing 957 eGenes, *without outlier removal:* Spearman’s rho=0.769, *with outlier removal:* R=0.71, R^2^=0.506, beta+/-SE=0.994+/-0.0295, T(1105)=33.649, p<2.2e-16, one *cis-*eQTL (from one eGene) removed as an extreme outlier).

Counterintuitively, the correlations with *cis*-eQTLs from non-cortical-like brain structures were just as strong (*Lateral Habenula (LHb):* 626 shared *cis-*eQTLs representing 617 eGenes, *without outlier removal:* Spearman’s rho=0.668, *with outlier removal*: R=0.677, R^2^=0.458, beta+/-SE=0.702+/-0.0306, T(622)=22.92, p<2.2e-16, 2 *cis-*eQTLs (representing one eGene) removed as extreme outliers; *Nucleus Accumbens (NAcc*, from Munro et al): 609 shared *cis-*eQTLs representing 604 eGenes, *without outlier removal:* Spearman’s rho=0.648, *with outlier removal*: R=0.672, R^2^=0.452, beta+/-SE=0.737+/-0.0330, T(606)=22.37, p<2.2e-16, one *cis-*eQTL (representing one eGene) removed as an extreme outlier; *Nucleus Accumbens (NAcc2*, from Mitchell & Hitzemann cohort): 853 shared *cis-*eQTLs representing 804 eGenes, Spearman’s rho=0.741, R=0.745, R^2^=0.555, beta+/-SE=0.606+/-0.0186, T(851)=32.55, p<2.2e-16).

The slopes for *cis*-eQTLs from non-brain tissue (adipose, liver) were less similar to those detected in the hippocampus (*Liver:* 997 shared *cis-*eQTLs representing 915 eGenes, *without outlier removal:* Spearman’s rho=0.306, *with outlier removal:* R=0.297, R^2^=0.088, beta+/-SE=0.223+/-0.0228, T(990)=9.78, p<2.2e-16, five *cis-*eQTLs (representing three eGenes) removed as extreme outliers; *Adipose:* 1,221 shared *cis-*eQTLs representing 1,101 eGenes, *without outlier removal*: Spearman’s rho=0.348, *with outlier removal*: R=0.354, R^2^=0.126, beta+/-SE=0.274+/-0.0207, T(1215)=13.21, p<2.2e-16, four *cis-*eQTLs (representing four eGenes) removed as extreme outliers). However, these tissue differences are difficult to interpret since they were also derived from different studies with differing sample sizes.

##### bHR/bLR Differential Expression Can Be Predicted Using the Hippocampal *cis*-eQTL Database

We next used our *cis*-eQTL database to predict the effect of genetic variation that segregates the bHR/bLR lines on gene expression. We found that 2,452 of the eVariants showed at least partial bHR/bLR segregation in the F_0_ rats (n=10 bHR/n=10 bLR sequenced in ^3^), such that if all subjects from one phenotype (*e.g.,* bHRs) had two reference alleles (0/0), all subjects from the other phenotype had at least one alternate allele (0/1; population segregation statistic G_st’_>0.27 ^21^). To predict the effect of these bHR/bLR segregated eVariants on gene expression, we calculated the allelic Log2FC (aFC) for each eVariant and assigned the direction of effect based on the allele frequency within the bLR vs. bHR F_0_ rats (**Fig 8A**). We found that these predictions correlated strongly with our F_0_ differential expression results (**Fig 8B**, 2,500 eGene/eVariant combinations: R=0.77, R2=0.588, rho=0.63, beta+/SE=0.528+/- 0.00885, T(2498)=59.708, *p*<2e-16), and with effect sizes within our previous bHR/bLR late generation meta-analysis (2,114 eGene/eVariant combinations: R=0.52, rho=0.61, beta+/SE=1.16+/-0.041, T(2112)=28.21, p<2e-16, **Fig S9**). These results demonstrated the validity of our hippocampal *cis*-eQTL database and confirmed that bHR/bLR differential expression of eGenes is likely to be driven by bHR/bLR genetic segregation.

##### The Strongest Co-localization of *cis*-eQTLs with Behavioral QTLs Tended to Predict F_2_ Differential Expression with Behavior

The strongest co-localization of *cis*-eQTLs with behavioral QTLs tended to predict F_2_ differential expression with behavior, especially when considering the predicted direction of effect based on the relationship between genotype and behavior within the larger F_2_ sample (adults: n=323, juveniles: n=216) and genotype and expression within the *cis*-eQTL analysis (n=245, **Fig 10A**). This was especially true if we narrowed our scope to *cis*-eQTLs that we had confirmed are segregated in bHR/bLRs with strong differential expression in bHR/bLRs matching what would be predicted based on the distribution of alleles in the two lines (from **Fig 8C;** 492 cis-eQTLs representing 456 eGenes).

The strongest correlations were between the strength of the co-localization of *cis*-eQTLs with QTLs for LocoScore (T-statistic, with predicted direction of effect) and the differential expression associated with LocoScore in the F_2_s (T-statistic) (**Fig 10B, Fig S10A&B**, full 5,747 *cis*-eQTLs: R=0.31, R^2^=0.099, beta+/SE=0.10+/-0.0041, T(5745)=25.11, p<2.2e-16) in a manner that strengthened when only considering the subset of eVariant/eGene combinations implicated in bHR/bLR phenotype (492 cis-eQTLs: R=0.56, R^2^=0.32, beta+/SE=0.144+/-0.00953, T(490)=15.09, p<2.2e-16). The correlation was also strong for EPM distance traveled (**Fig S10C&D,** 5,747 *cis*-eQTLs: R=0.30, R^2^=0.092, beta+/-SE=0.14+/-0.0060, T(5745)=24.18, p<2.2e-16) in a manner that similarly strengthened when considering eVariant/eGene combinations implicated in bHR/bLR phenotype (**Fig S10E-J**: 492 *cis*-eQTLs: R=0.57, R^2^=0.32, beta+/-SE=0.249+/-0.163, T(490)=15.29, p<2.2e-16,). The correlation was slightly weaker for PavCA Index, EPM Time Immobile, and EPM %Time in the Open Arms (full 5,747 *cis*-eQTLs: *PavCA Index:* R=0.28, R^2^=0.078, beta+/-SE=0.165+/-0.00749, T(5745)=22.01, p<2.2e-16; *EPM Time Immobile:* R=0.26, R^2^=0.070, beta+/-SE=0.159+/-0.0077, T(5745)=20.823, p<2.2e-16; *EPM* %*Time in the Open Arms:* R=0.23, R^2^=0.054, Beta+/-SE=0.150+/-0.0082, T(5745)=18.165, p<2.2e-16), in a manner that similarly strengthened when considering eVariant/eGene combinations implicated in bHR/bLR phenotype (492 *cis*-eQTLs: *PavCA Index:* R=0.49, R^2^=0.24, beta+/-SE=0.271+/-0.0216, T(490)=12.55, p<2.2e-16; *EPM Time Immobile:* R=0.44, R^2^=0.19, beta+/-SE=0.235+/-0.0217, T(490)=10.85, p<2.2e-16; *EPM %Time Open Arms:* R=0.33, R^2^=0.11, beta+/-SE=0.192+/-0.0246, T(490)=7.829, p=3.07e-14).

We also observed a correlation when comparing F_2_ differential expression to SMR co-localization results with QTLs for two analogous juvenile behaviors (**Fig 10C**, **Fig S11A-D:** full 5,746 *cis*-eQTLs: *Adult EPM Distance Traveled vs. Juvenile OF Distance Traveled:* R=0.123, R^2^=0.0151, beta+/-SE=0.0504+/-0.00536, T(5744)=9.398, p<2e-16; *Adult EPM Time Immobile vs. Juvenile OF Time Immobile:* R=0.102, R^2^=0.0104, beta+/-SE=0.0551+/-0.0071, T(5744)=7.76, p=9.56e-15; subset of 492 cis-eQTLs implicated in bHR/bLR phenotype: *Adult EPM Distance Traveled vs. Juvenile OF Distance Traveled:* R=0.35, R^2^=0.122, beta+/-SE=0.115, T(490)=8.249, p=1.48e-15, *Adult EPM Time Immobile vs. Juvenile OF Time Immobile:* R=0.26, R^2^=0.0679, beta+/-SE=0.115+/-0.0192, T(490)=5.975, p=4.44e-09), but not when comparing the F_2_ differential expression for the anxiety measure in the adults (EPM % time in the open arms) to the SMR co-localization results using QTLs for the anxiety measure in the juveniles (OF % time in the center) (**Fig S11E&F:** full 5,746 *cis*-eQTLs: R=0.0038, R^2^= 1.453e-05, beta+/-SE= 0.00228+/-0.00789, T(5744)=0.289; subset of 492 cis-eQTLs implicated in bHR/bLR phenotype: R=0.0757, R^2^=0.00574, beta+/-SE=0.0369+/-0.0219, T(490)=1.682).

### Supplementary Tables

| **Behavior** |  | **F Stat** | **DF** | **P Value** |
| --- | --- | --- | --- | --- |
| **LocoScore** | **Phenotype** | 212.764 | 2, 268 | <2.2E-16 |
|  | **Sex** | 3.822 | 1, 268 | 0.0516 |
|  | **Phenotype x Sex** | 2.065 | 2, 268 | 0.129 |
| **EPM Distance Traveled** | **Phenotype** | 43.903 | 2, 268 | <2.2E-16 |
|  | **Sex** | 9.74 | 1, 268 | 0.002 |
|  | **Phenotype x Sex** | 3.633 | 2, 268 | 0.0278 |
| **EPM % Time Open Arms** | **Phenotype** | 14.137 | 2, 268 | 1.46E-06 |
|  | **Sex** | 7.582 | 1, 268 | 0.006299 |
|  | **Phenotype x Sex** | 4.908 | 2, 268 | 0.00806 |
| **EPM Time Immobile** | **Phenotype** | 39.289 | 2, 268 | <1.09E-15 |
|  | **Sex** | 11.792 | 1, 268 | 0.00069 |
|  | **Phenotype x Sex** | 15.306 | 2, 268 | 5.09E-07 |
| **PavCA: ST/GT Classification** | **Phenotype** |  | 2, 207 |  |
|  | **Sex** | 49.743 | 1, 207 | 2.58E-11 |
|  | **Phenotype x Sex** |  | 2, 207 |  |
|  |  | **ST** | **IN** | **GT** |
|  | **Number of M/F:** | 21/60 | 58/30 | 29/11 |
|  | **Fisher's Exact Test:** | OR: 0.13; CI: 0.05-0.34; p=1.472E-06 | | |

Table S 1: The bHR/bLR (F_0_) cross-breeding scheme produced F_2_ animals with a range of intermediate behavioral phenotypes.

Statistical results are shown in the table. Analysis of variance was performed for each of the behaviors, with sex and phenotype as factors. A Fisher’s Exact Test was performed on the ratios of male to female animals classified as Sign Trackers (ST), Intermediate (IN), or Goal Trackers (GT) on the Pavlovian Conditioned Approach (PavCA) task. F Stat = F statistic; DF = degrees of freedom; P Value = significance value; M/F = male/female ratio: OR = odds ratio; CI = confidence interval.

| **Behavioral Correlations** | **R** | **Beta +/- SE** | **T Stat** | **DF** | **P value** |
| --- | --- | --- | --- | --- | --- |
| LocoScore vs. EPM Distance Traveled | 0.47 | 0.321 +/- 0.0381 | 8.433 | 247 | 2.85E-15 |
| LocoScore vs. EPM % Time Open Arms | 0.30 | 8.519 +/- 1.74 | 4.901 | 248 | 1.72E-06 |
| LocoScore vs. EPM Time Immobile | -0.45 | -5.202 +/- 0.662 | 7.86 | 248 | 1.17E-13 |
| EPM Distance Traveled vs. EPM % Time Open Arms | 0.54 | 0.0127 +/- 0.00127 | 10.003 | 247 | 2.20E-16 |
| EPM % Time Open Arms vs. EPM Time Immobile | -0.45 | -0.184 +/- 0.0230 | -7.975 | 248 | 5.61E-14 |
| LocoScore vs PavCA Index | 0.46 | 249.36 +/- 33.67 | 7.407 | 207 | 3.22E-12 |
| EPM Distance Traveled vs. PavCA Index | 0.40 | 308.86 +/- 49.07 | 6.294 | 206 | 1.83E-09 |
| EPM % Time Open Arms vs. PavCA Index | 0.28 | 5.036 +/- 1.207 | 4.172 | 207 | 4.44E-05 |
| EPM Time Immobile vs. PavCA Index | 0.43 | -18.527 +/- 2.734 | 6.777 | 207 | 1.26E-10 |

Table S 2. Behaviors that differentiate the bHR/bLR lines, such as exploratory locomotion, anxiety, and PavCA, remained correlated in the F_2_ animals.

The correlation coefficients and related statistics for the behavioral correlations are shown in this table. R = Pearson correlation coefficient; Beta +/- SE = regression slope+/-standard error; T Stat = t-statistic for regression slope; DF= degrees of freedom; P value = p-value for regression slope.

Table S 3. Differential expression results for each of the variables of interest from the F_0_ and F_2_ datasets, as well as the full cis-eQTL and SMR results for each eGene.

This file (TableS3_F0_Meta_F2_results_ForSuppl_20240223.xlsx) contains four spreadsheets. The first spreadsheet (“Column Definitions 1”) contains the definitions for all the columns in spreadsheet 2 (“Full DE Results”). The second spreadsheet (“Full DE Results”) contains the bLR vs. bHR differential expression results for all Ensembl-annotated genes present in the F_0_ dataset (n=13,786), and their associated bLR vs. bHR results from the late generation RNA-Seq meta-analysis ^4^, as well as the differential expression results for all F_2_ behaviors (LocoScore, EPM Distance Traveled, EPM Time Immobile, EPM % Time Open Arms, PavCA Index). Conditional formatting is used to make the spreadsheet easier to navigate: Effect sizes (Log2 Fold Change (FC), meta-analysis estimated d) and t-statistics are colored using a gradient from red to green, with red indicating upregulation in bLR-like animals and green indicating upregulation in bHR-like animals. P-values and false discovery rates (FDRs) are colored using a gradient from yellow to grey, with yellow indicating lower (“more significant”) values and grey indicating higher (“less significant”) values. The third spreadsheet (“Column Definitions 2) contains the definitions for all the columns in spreadsheet 4 (“eQTLs DE Results”). The fourth spreadsheet (“eQTLs DE Results”) includes the subset of differential expression results for genes identified as having significant cis-eQTLs, aligned with the cis-eQTL and SMR results for each significant eGene/eVariant combination (“eQTL_Gene_eVariant_ID”). Formatting follows the conventions in spreadsheet 2 (“Full DE Results”).

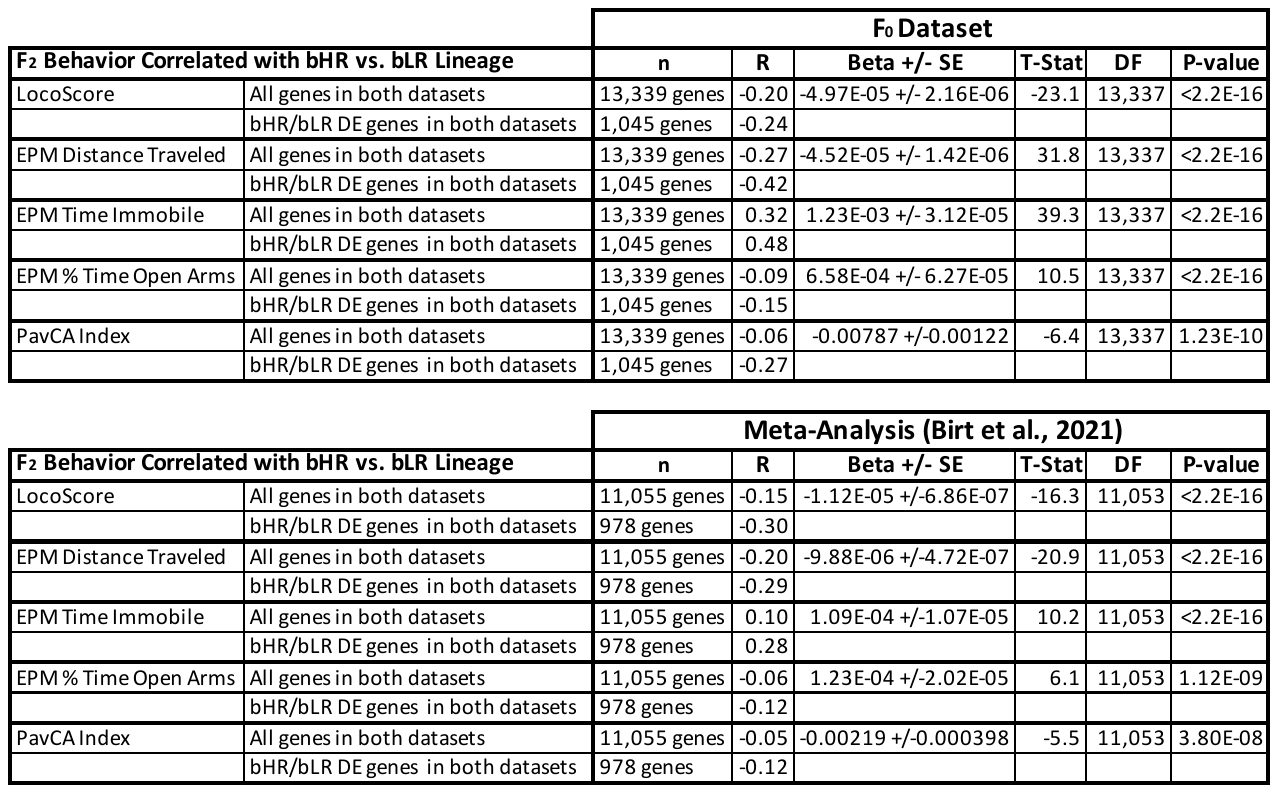

Table S 4. The pattern of gene expression differentiating bLR vs. bHR rats predicts the pattern of gene expression associated with F_2_ behavior.

This table shows the correlation coefficients and related regression statistics for the correlations between the differential expression results (Log2FC) for each of the F_2_ behaviors in comparison to the differential expression results for bHR/bLR lineage from either the F_0_ dataset (Log2FC) or from our previous late generation RNA-Seq meta-analysis (estimated d, ^4^). n= number of genes included in analysis, R = Pearson correlation coefficient; Beta +/- SE = regression slope +/- standard error; T-Stat = t-statistic for regression slope; DF= degrees of freedom; P-value = p-value for regression slope.

Table S 5. Gene set enrichment results: bHR/bLR phenotype is associated with differential expression of genes involved in growth and proliferation, microglia, mitochondrial function, and the regulation of oxidative stress.

This file (TableS5_fGSEA_F0_Meta_F2_results.xlsx) contains three spreadsheets. The first spreadsheet contains the definitions for all the columns in the other spreadsheets. The second spreadsheet contains the full results from the directional version of the gene set enrichment analysis (fGSEA algorithm: ^13^). To run this analysis we created a continuous variable representing bLR-like vs. bHR-like differential expression for each gene by averaging the t-statistics for bLR vs. bHR comparisons in the F_0_ dataset and former late generation RNA-Seq meta-analysis ^4^ and the t-statistics for each of the F_2_ behaviors (with bHR-like phenotype set as reference, so that positive t-statistics indicate upregulation in bLR-like animals). Conditional formatting is used to make the spreadsheet easier to navigate: Effect sizes (gene set enrichment score (ES) and gene set normalized enrichment score (NES)) are colored using a gradient from red to green, with red indicating an enrichment of upregulation in bLR-like animals and green indicating an enrichment of upregulation in bHR-like animals for the genes in that gene set. P-values and false discovery rates (FDRs) are colored using a gradient from yellow to grey, with yellow indicating lower (“more significant”) values and grey indicating higher (“less significant”) values. The third spreadsheet contains the full results from the non-directional version of the gene set enrichment analysis, which was run using the absolute value of the average t-statistic discussed above.

### Supplementary Figures

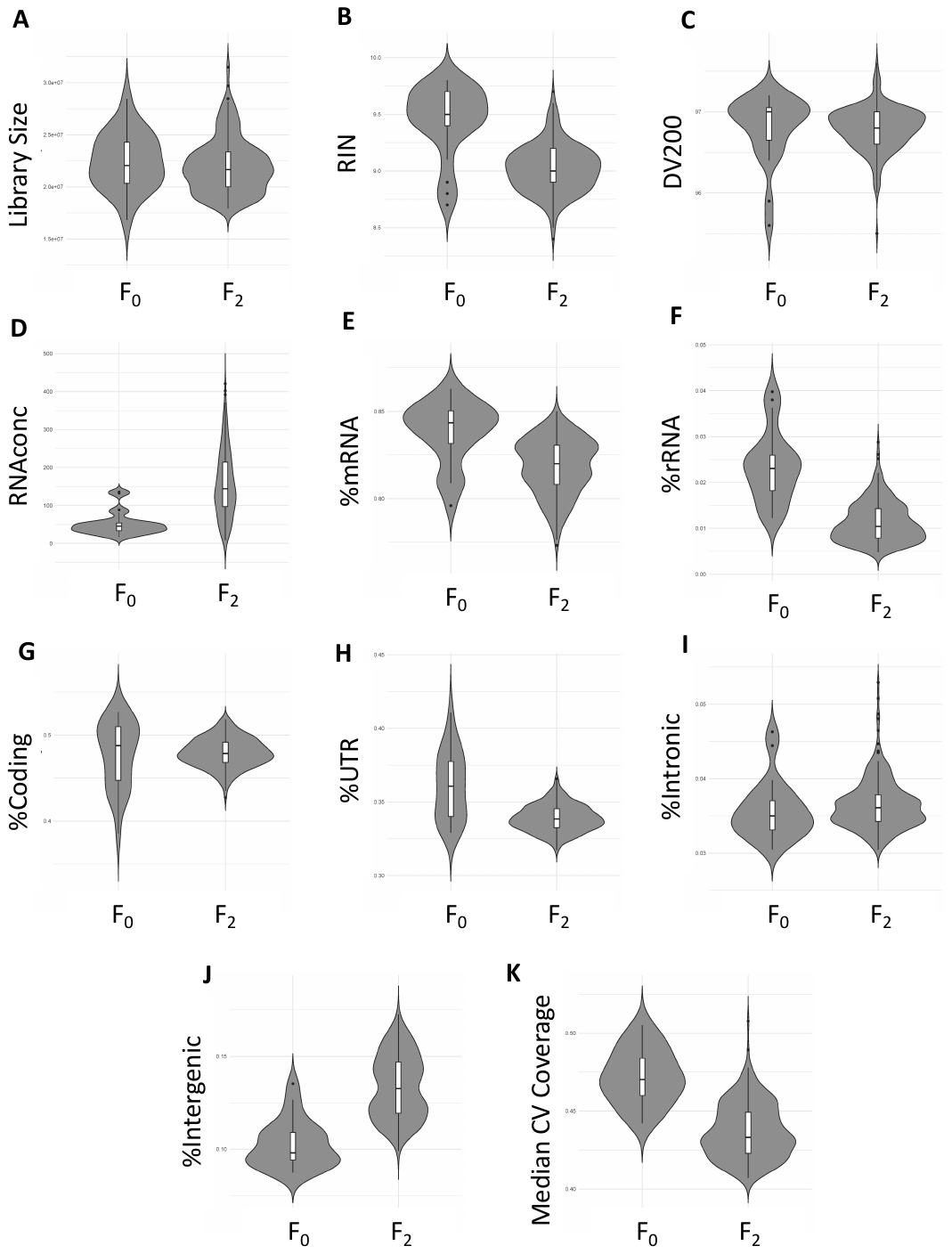

Fig S 1. Quality control: The distribution of RNA metrics within the F_0_ and F_2_ datasets indicate high quality RNA.

The distributions for a variety of RNA metrics are depicted using violin plots for F_0_ and F_2_ samples following the removal of outlier samples (F_0_: n=23, F_2_: n=245). F_0_ and F_2_ samples had a similar distribution for many RNA metrics, with the exception that F_0_ samples had a much lower RNA concentration (**D**), higher RIN (**B**), greater %rRNA (**F**) and UTR (**H**), lower %Intergenic (**J**), and higher median CV coverage (**K**). Each plot contains the median, interquartile range, and 1.5x interquartile range for the given metric of the F_0_ and F_2_ samples. The width of each density curve corresponds to the approximate frequency of data points in each region. Library Size=the total number of mapped reads, RIN=RNA integrity number, DV200=percentage of fragments >200 nucleotides, RNA conc=RNA concentration, %mRNA=percent messenger RNA, %rRNA=percent ribosomal RNA, %Coding=percent coding RNA, %UTR=percent untranslated region, %Intronic=percent intronic, %Intergenic=percent intergenic, Median CV Coverage=median coefficient of variation of coverage.

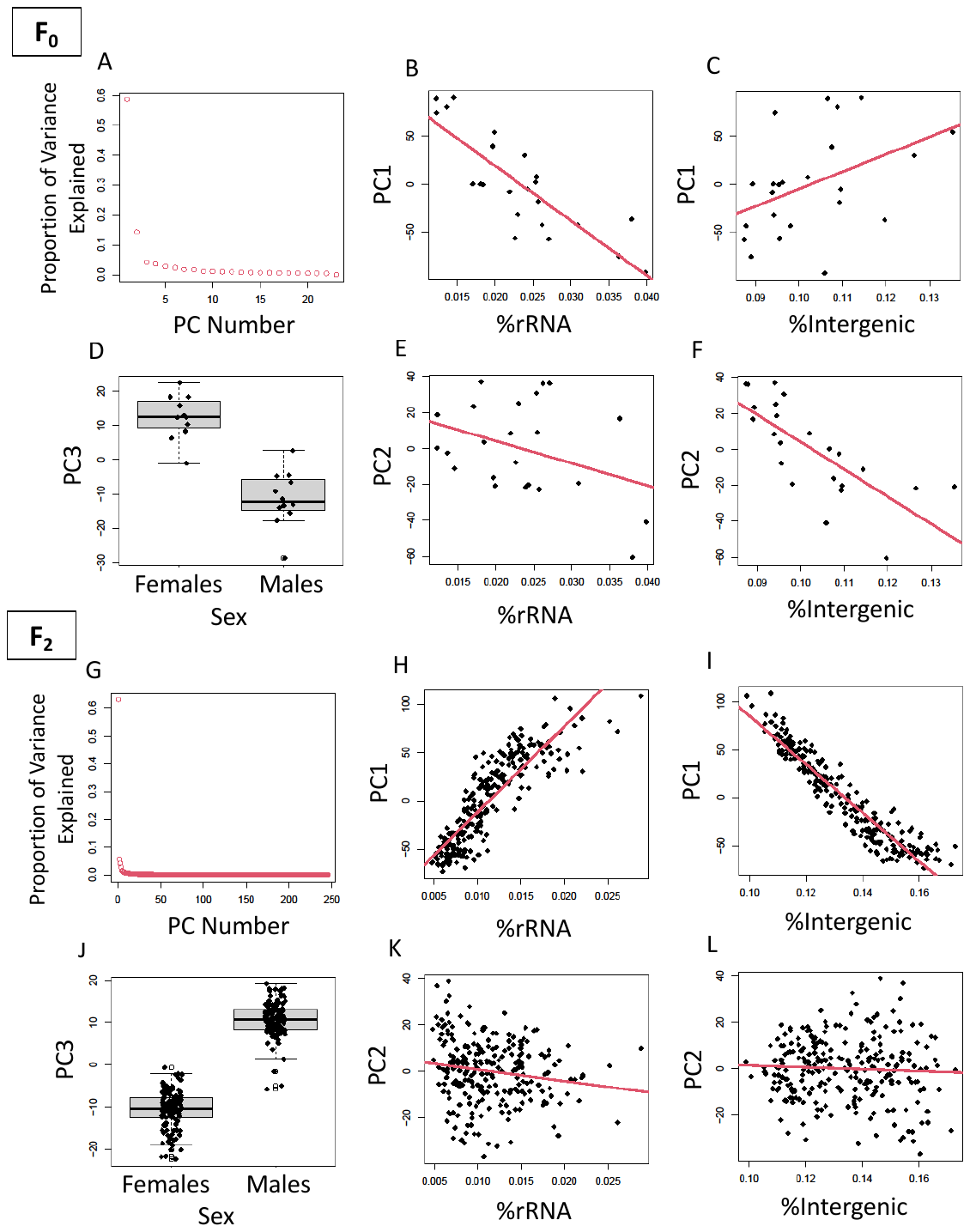

Fig S 2. Quality control: RNA metrics and sex correlate strongly with the top principal components (PCs) of variation in our datasets.

Principal Components Analysis is a dimension reduction method that identifies the strongest gradients of variation present in large datasets. These gradients are called PCs and are represented numerically, with arbitrarily assigned directionality. Within our experiment, the top PC of variation (PC1) explained most of the variation in the data (>60%) from both the F_0_s (**A**) and the F_2_s (**G**). After examining the relationships between the top PCs and numerous technical variables and RNA metrics using both simple bivariate relationships and an automated model selection procedure, we chose RNA metrics as covariates that were most strongly related to the top PCs in both datasets and were not completely redundant with each other: %rRNA and %Intergenic. For the F_0_s, %rRNA was strongly associated with PC1 (**B**) and moderately associated with PC2 (**E**), whereas %Intergenic was moderately associated with PC1 (**C**) and strongly associated with PC2 (**F**). For the F_2_s, %rRNA and %Intergenic were both strongly associated with PC1 in a manner that was at least partially independent (**H**, **I**), and weakly associated with PC2 (**K**, **L**). PC3 was strongly related to sex for both the F_0_ and F_2_ datasets (**D** and **J**).

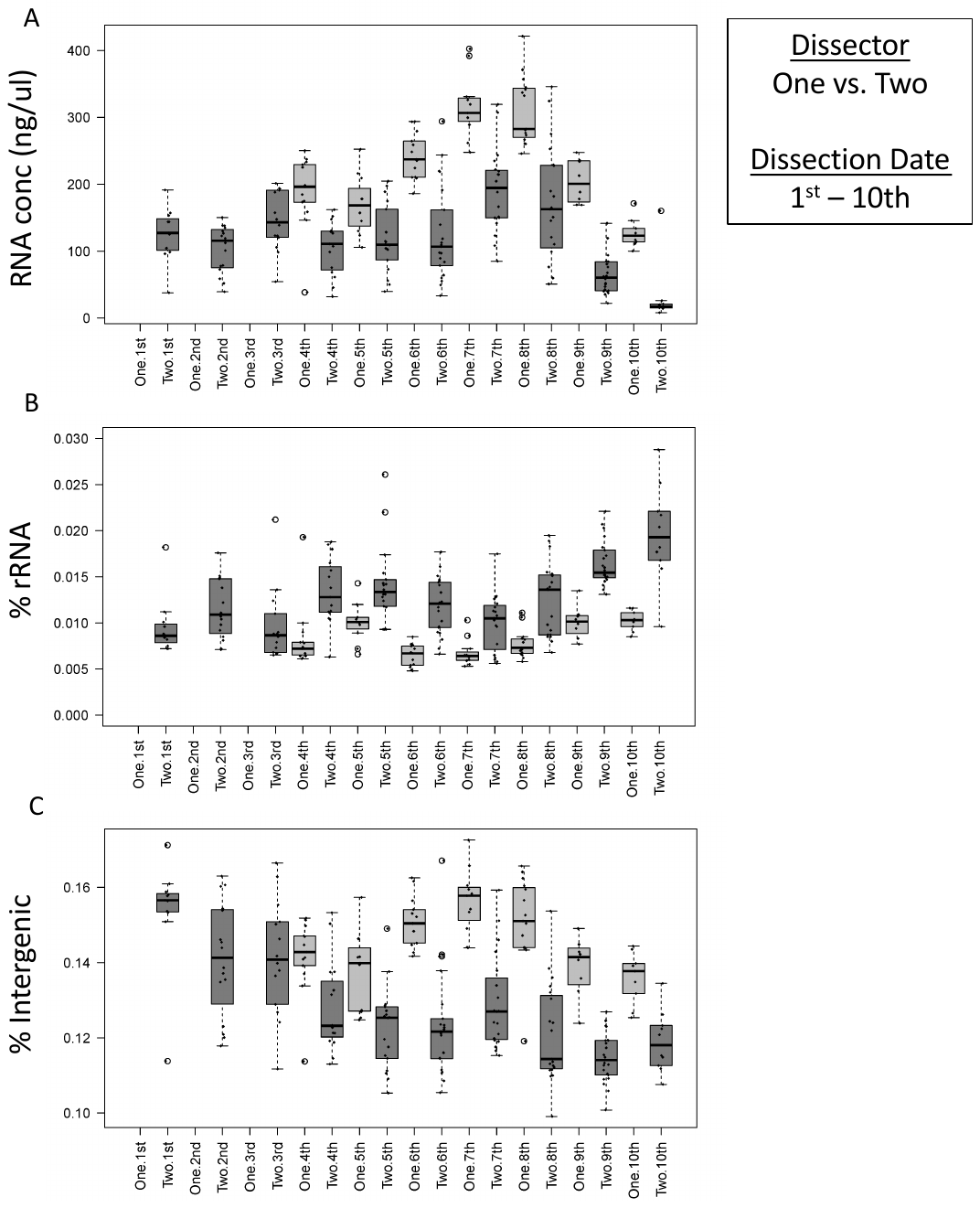

Fig S 3. Quality control: Variation in RNA metrics is likely to have arisen from variation in sample collection and processing.

For example, RNA concentration (**A**), %rRNA (**B**) and %Intergenic RNA (**C**), varied significantly across tissue dissection date (1^st^ – 10^th^) and by dissector (One vs. Two). In addition to the median value of each of the three metrics varying across date of dissection, the RNA concentration was found to be significantly higher (**A**), the %rRNA significantly lower (**B**) and the %Intergenic RNA (**C**) significantly higher in the first compared to the second dissector. These batch variables and RNA metrics were all considered as potential covariates during differential expression model selection. Dissector=One vs. Two; Dissection Date=1^st^-10^th^

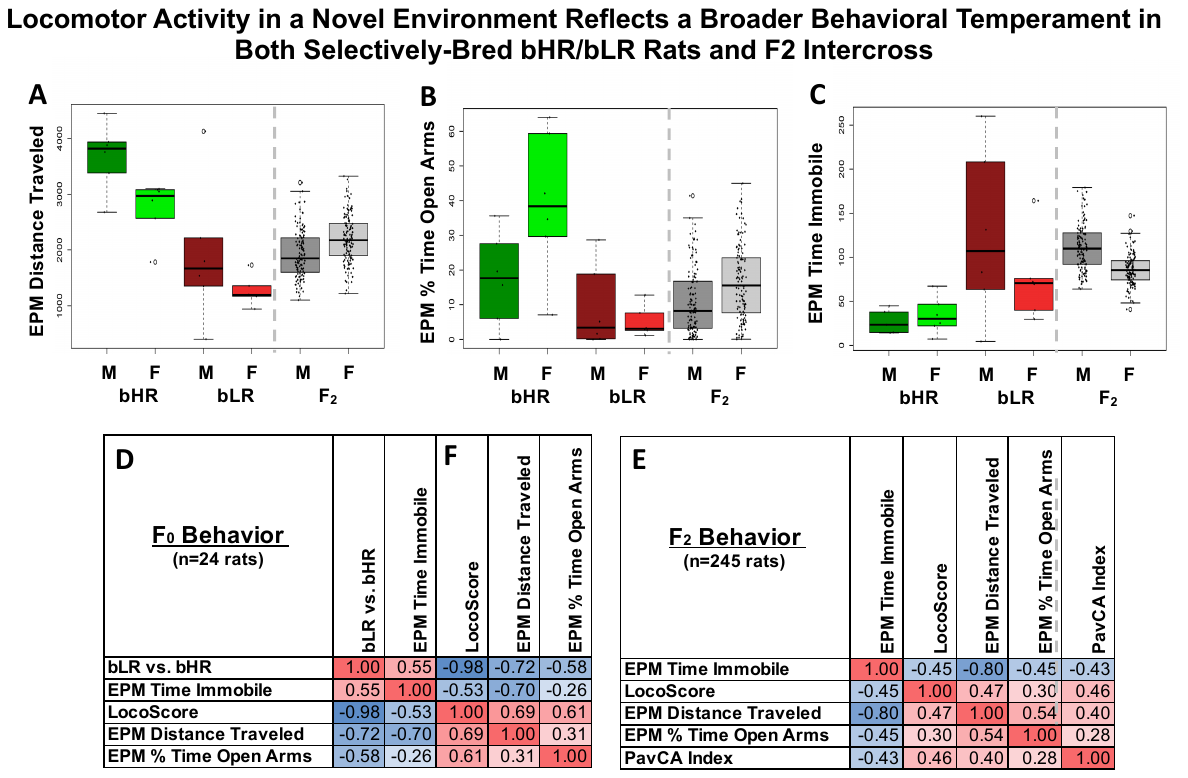

Fig S 4. Additional figure panels illustrating that locomotor activity in a novel environment reflects a broader behavioral temperament in both selectively-bred bHR/bLR lines and F_2_ intercross rats.

**A-C)** The bHR/bLR (F_0_) cross-breeding scheme produced F_2_ animals with intermediate behavior which fell between the extreme bHR-like and bLR-like phenotypes. Similar to **Fig 3B**, depicted are the behavioral measurements for the F_0_ and F_2_ rats included in the current RNA-Seq experiment. For all measured behaviors, all group differences (ANOVA: F_0_ bHR vs. F_0_ bLR vs. F_2_) were highly significant (p<1.5e-06). **A)** bHRs traveled a greater distance in the elevated plus maze (EPM) compared to bLRs, with the F_2_s exhibiting intermediate distances. **B)** bHRs show less anxiety by spending a greater percent time in the EPM open arms (especially females) than bLRs, with the F_2_s again showing intermediate values. **C)** bHRs also showed less anxiety by spending less time immobile (sec) in the EPM than bLRs or F_2_s. **D-E)** Behaviors that diverged during bHR/bLR selective breeding for LocoScore remained correlated in F_2_s, as illustrated using correlation matrices. Each correlation matrix is symmetrical along the diagonal, with correlation coefficients (R) reflecting the correlation between the complete observations within each pairwise comparison. Positive correlations are shaded in red, negative correlations are shaded in blue. Example scatterplots illustrating the correlations between LocoScore and other bHR/bLR distinctive behaviors in the F_2_s can be found in **Fig 3C-D. D)** The correlation between bLR vs. bHR lineage (bHR coded as reference (0), bLR coded as 1) and each of the behavioral variables measured in the F_0_ dataset. By definition, bLRs showed a lower LocoScore, but also less distance traveled in the EPM and less time spent in the EPM open arms (which indicates greater anxiety-like behavior). Greater PavCA GT behavior in bLRs and ST behavior in bHRs was previously shown in ^27^ but was not measured in the F_0_s. **E)** Within the F_2_ dataset, LocoScore, EPM behavior, and PavCA behavior showed a similar pattern of correlations (all correlations: p<4.5E-05).

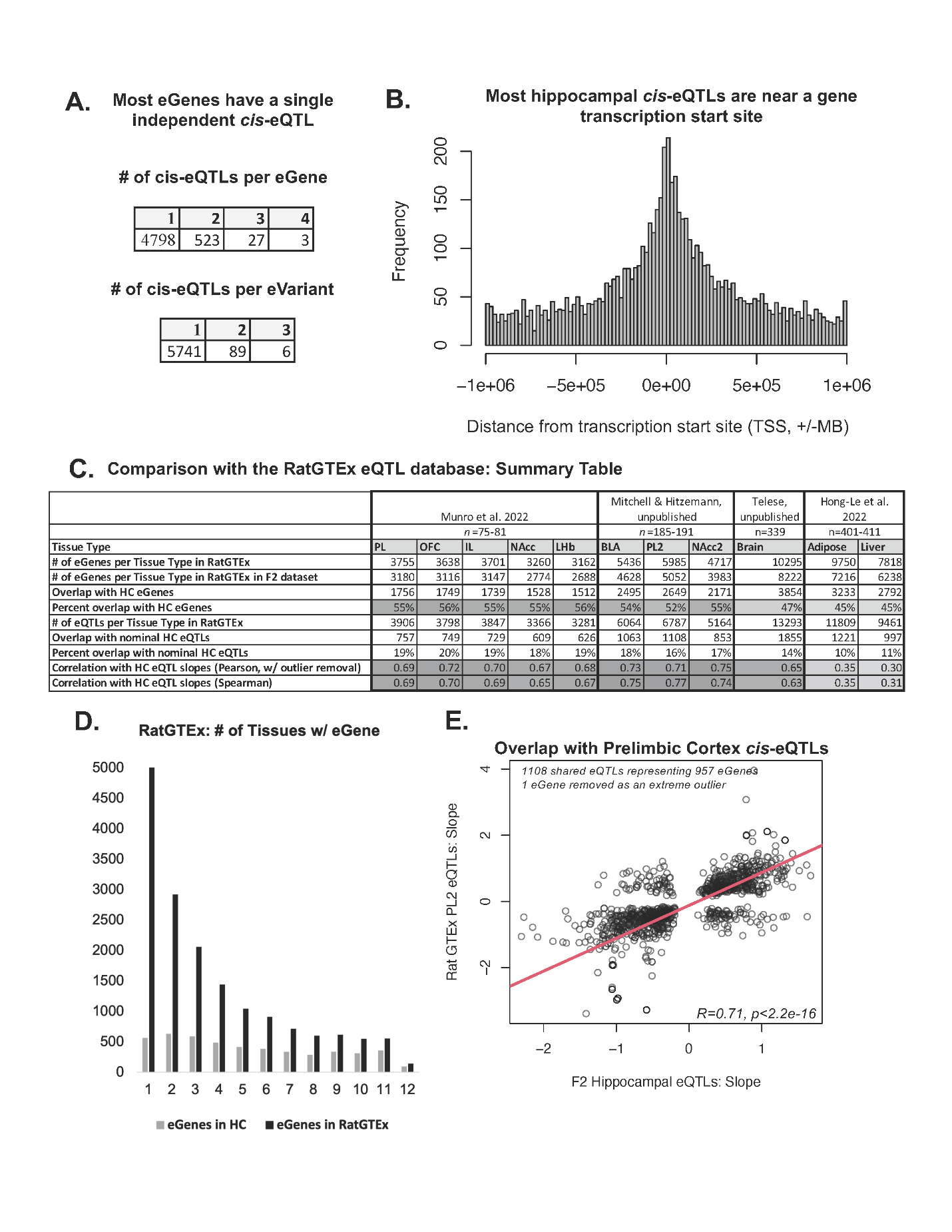
Fig S 5. Constructing a hippocampal cis-eQTL database.

We integrated our current F_2_ transcriptional profiling data (n=245) with our previous whole genome sequencing results (n=4,425,349 SNPs, ^3^) to identify genes with hippocampal expression tightly correlated with nearby genetic variation (cis-eQTLs). This analysis identified 5,351 genes (eGenes) with expression that correlated strongly (FDR<0.05) with genetic variation within +/-1 MB of their transcription start sites (TSS). **A)** Using stepwise regression, we identified additional conditionally-independent cis-eQTLs beyond the strongest cis-eQTL for each eGene, distinguishing a final total of 5,937 cis-eQTLs representing 5,836 unique eVariants. Two tables summarize the number of cis-eQTLs per eGene and the number of cis-eQTLs per eVariant. **B)** Similar to previous cis-eQTL analyses, these eVariants were predominantly located within +/-400 kB of the TSS of their respective eGene. A histogram illustrates the frequency that a cis-eQTL in our hippocampal database (n=5,937) has a particular distance (+/-Mb) from the TSS for their respective eGene (x-axis). **C)** A table comparing our hippocampal cis-eQTL database with the existing rat cis-eQTL databases in RatGTEx. The table includes the citations in which the cis-eQTLs were identified (^15,23^, Mitchell & Hitzemann, unpublished, Telese unpublished), along with their respective sample sizes and tissue type (abbreviations: PL: prelimbic cortex, OFC: orbitofrontal cortex, IL: infralimbic cortex, NAcc: nucleus accumbens, LHb: lateral habenula, BLA: basolateral amygdala, Brain: whole brain hemisphere). It also includes the number of eGenes identified per tissue type within RatGTEx, the number of those eGenes that were represented in our F_2_ dataset, the number of those eGenes that were also significant eGenes within our hippocampal cis-eQTL analysis, and the percent that hippocampal eGenes overlap with those identified in the RatGTEx tissue. The table also includes the number of cis-eQTLs identified in each tissue within RatGTEx, the number of those cis-eQTLs that were also found to be at least nominally (p<0.05) significant cis-eQTLs within the hippocampus, and their percent overlap. Finally, for those overlapping cis-eQTLs, we examined the replication of the cis-eQTL slopes identified in the two tissues using parametric (Pearson’s correlation) and non-parametric (Spearman’s rho) analyses. **D)** The majority of the eGenes that we identified within the hippocampus were also significant eGenes within at least four other tissues with RatGTEx. The bar chart illustrates the number of tissues within which each eGene was found to be significantly related to genetic variation (cis-eQTL), as either identified in our study in the hippocampus or in RatGTEx. **E)** Previously-identified brain cis-eQTLs showed a similar direction of effect on gene expression within the hippocampus when there was at least a nominal (p<0.05) relationship in our dataset. Shown is an example comparison of the cis-eQTL slopes identified within our hippocampal study (x-axis) and the Prelimbic Cortex (y-axis: PL2, from Mitchell & Hitzemann cohort). There were 1,108 shared cis-eQTLs representing 957 eGenes that were found in the Prelimbic cortex that had at least nominal (p<0.05) representation in the hippocampus. The slopes of the cis-eQTLs from the two brain regions were found to be strongly correlated (without outlier removal: Spearman’s rho=0.769, with outlier removal: R=0.71, R^2^=0.506, beta+/-SE=0.994+/-0.0295, T(1105)=33.649, p<2.2e-16, one cis-eQTL (from one eGene) removed as an extreme outlier).

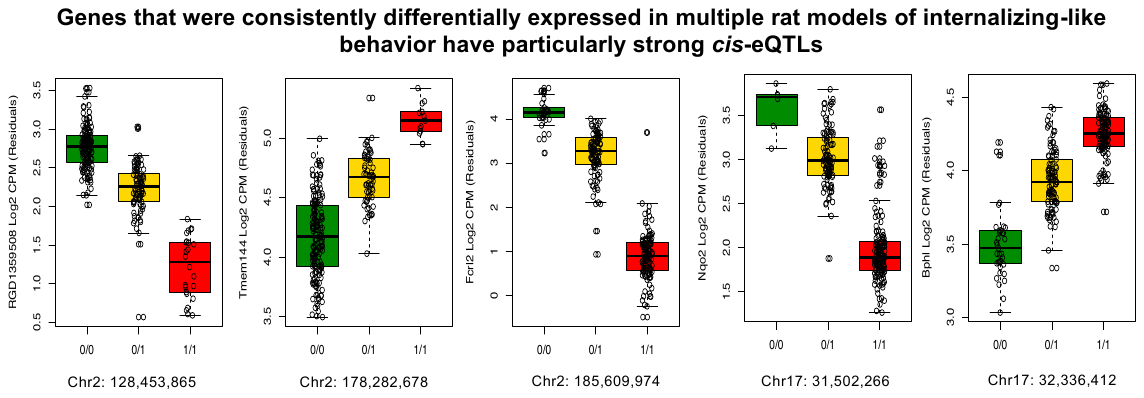

Fig S 6. Five of the six genes that were consistently differentially expressed in multiple rat models of internalizing-like behavior have particularly strong cis-eQTLs.

Each boxplot illustrates the relationship between eVariant genotype (0=reference allele, 1=alternate allele) and gene expression (Log2 counts per million (CPM)) within the F_2_ dataset (n=245) after correcting for the technical covariates included in our differential expression model. The boxes for each allele are colored to indicate segregation within our bred lines (green=more of the allele is present in bHRs, red=more of the allele is present in bLRs).

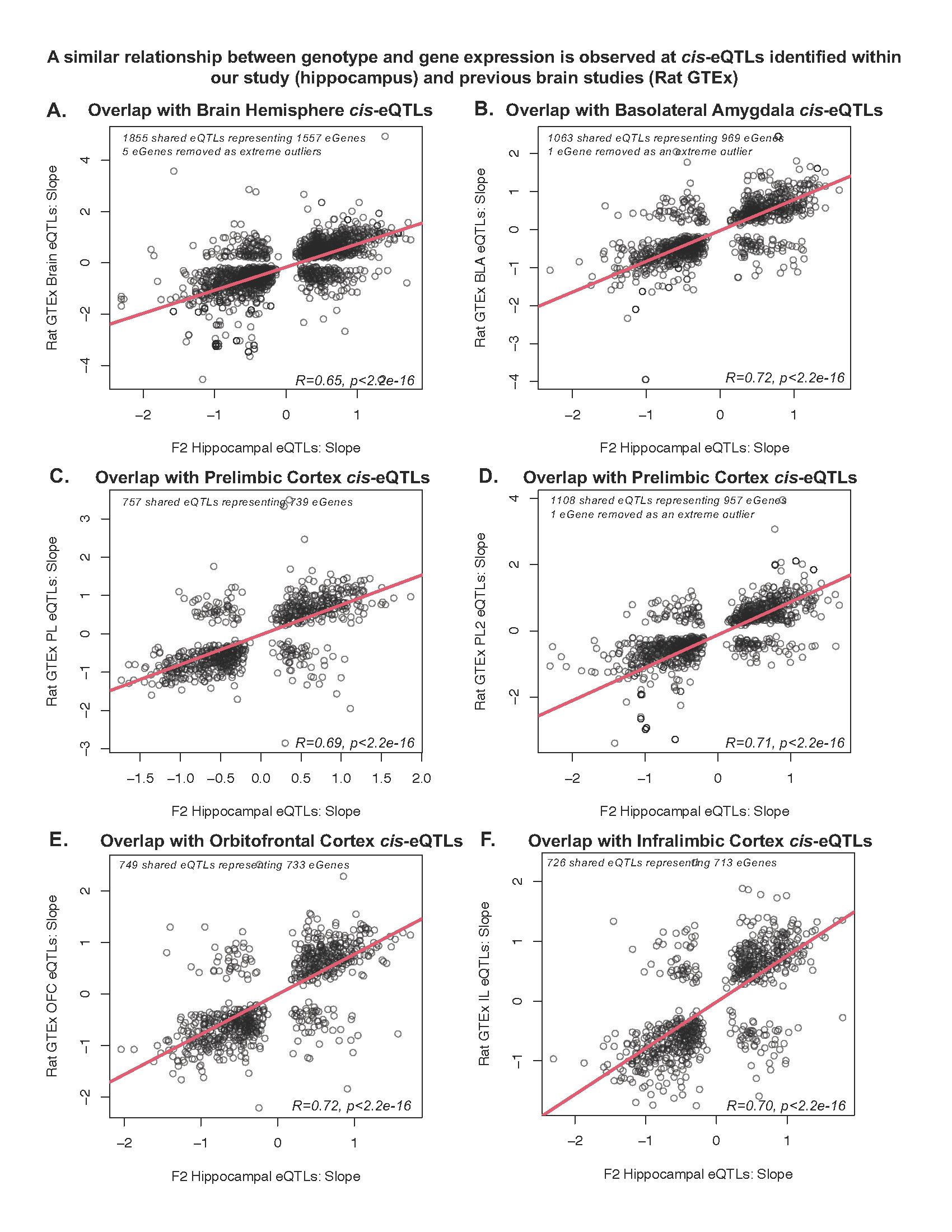

Fig S 7. A similar relationship between genotype and gene expression is observed at cis-eQTLs identified within our study (hippocampus) and previous brain studies (RatGTEx).

In general, the cis-eQTLs identified in RatGTEx tended to have a similar slope as what we observed in the hippocampus. This was particularly true for brain tissue. **A)** We found that 1,855 cis-eQTLs identified using whole rat brain hemispheres (representing 1,556 eGenes) were at least nominally detectable in the hippocampus, with many showing a similar slope (y-axis: cis-eQTL slope in rat hemispheres, x-axis: cis-eQTL slope in the hippocampus) (without outlier removal: Spearman’s rho=0.628, with outlier removal: R=0.65, R^2^=0.391, beta+/-SE=0.902+/-0.0264, T(1819)=34.157, p<2.2e-16, 34 cis-eQTLs (from five eGenes) removed as extreme outliers). **B)** Within another subcortical region with cortical-like structure, the basolateral amygdala (BLA), 1,063 cis-eQTLs (representing 969 eGenes) were identified that were nominally detectable in the hippocampus, with many showing a similar slope (y-axis: cis-eQTL slope in the BLA, x-axis: cis-eQTL slopes in the hippocampus) (without outlier removal: Spearman’s rho=0.752, with outlier removal: R=0.72, R^2^=0.53, beta+/-SE=0.81+/-0.0235, T(1053)=34.56, p<2.2e-16, eight cis-eQTLs (from one eGene) removed as extreme outliers). **C)** Within the prelimbic cortex (PL, from Munro et al. cohort): 757 cis-eQTLs (representing 739 eGenes) were identified that were nominally detectable in the hippocampus, with many showing a similar slope (y-axis: cis-eQTL slope in the PL, x-axis cis-eQTL slopes in the hippocampus) (Spearman’s rho=0.694, R=0.69, R^2^=0.481, beta+/-SE=0.778+/-0.0294, T(755)=26.465, p<2.2e-16. **D)** This same pattern can be seen in relationship with cis-eQTLs detected in an independent prelimbic cortex dataset (PL2, from Mitchell & Hitzemann cohort). In this case, 1,108 cis-eQTLs (representing 957 eGenes) were identified that were nominally detectable in the hippocampus, with many showing a similar slope (y-axis: cis-eQTL slope in PL2, x-axis: cis-eQTL slopes in the hippocampus) (without outlier removal: Spearman’s rho=0.769, with outlier removal: R=0.71, R^2^=0.506, beta+/-SE=0.994+/-0.0295, T(1105)=33.649, p<2.2e-16, one cis-eQTL (from one eGene) removed as an extreme outlier). **E)** Within the orbitofrontal cortex (OFC), 749 cis-eQTLs (representing 733 eGenes) were identified that were nominally detectable in the hippocampus, with many showing a similar slope (y-axis: cis-eQTL slope in OFC, x-axis: cis-eQTL slopes in the hippocampus) (Spearman’s rho=0.704, R=0.722, R^2^=0.521, beta+/-SE=0.778+/-0.0273, T(747)=28.523, p<2.2e-16). **F)** Within the infralimbic cortex (IL) 726 cis-eQTLs (representing 713 eGenes) were identified that were nominally detectable in the hippocampus, with many showing a similar slope (y-axis: cis-eQTL slope in the IL, x-axis cis-eQTL slopes in the hippocampus) (Spearman’s rho=0.692, R=0.703, R^2^=0.494, beta+/-SE=0.773+/-0.0290, T(727)=26.63, p<2.2e-16).

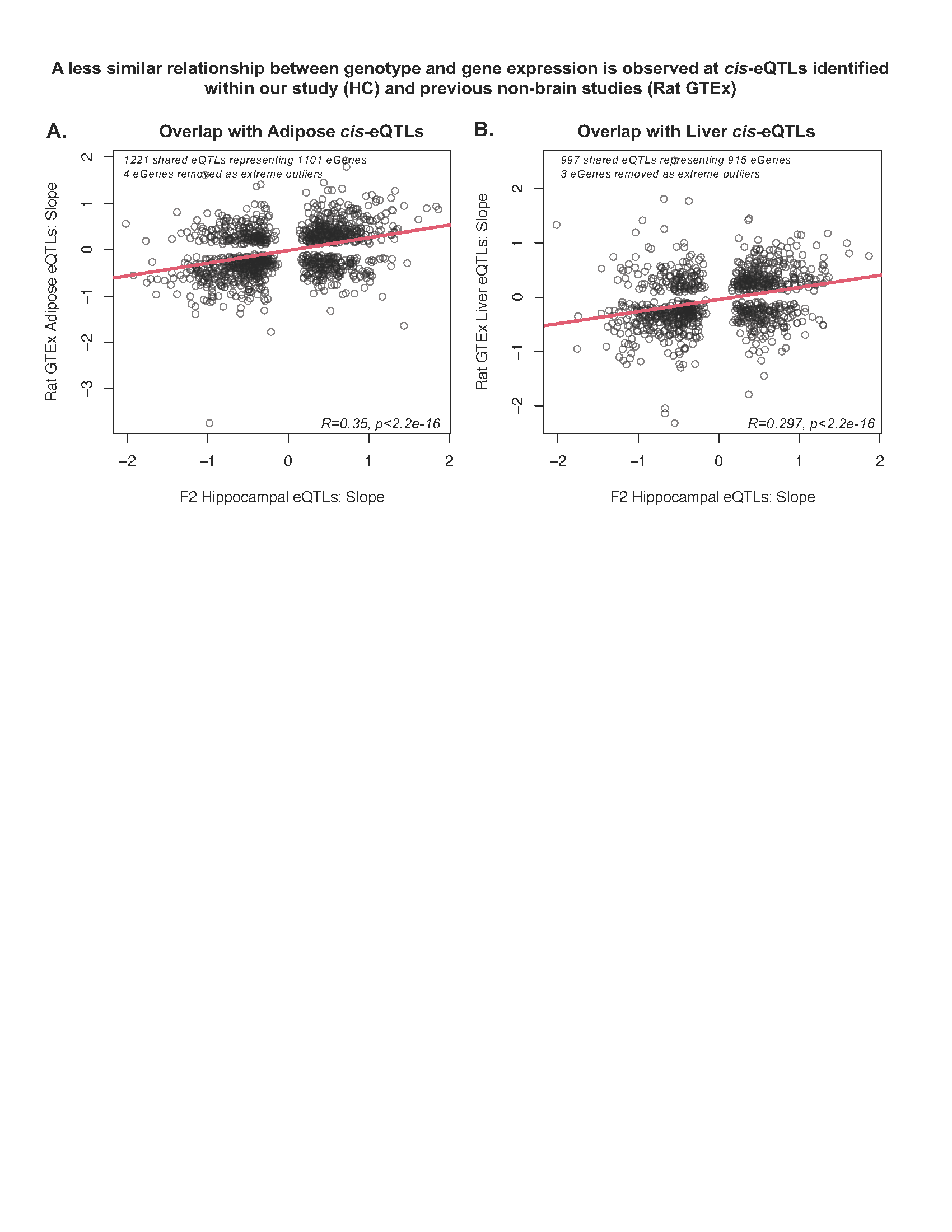

***Fig S 8. A less similar relationship between genotype and gene expression is observed at cis-eQTLs identified within our study (hippocampus; HC) and previous non-brain studies (RatGTEx).***

**A)** We found that 1,221 cis-eQTLs identified in adipose tissue (representing 1,101 eGenes) were at least nominally detectable in the hippocampus, with some showing a similar slope (y-axis: cis-eQTL slope in adipose tissue, x-axis: cis-eQTL slope in the hippocampus) (without outlier removal: Spearman’s rho=0.348, with outlier removal: R=0.354, R^2^=0.126, beta+/-SE=0.274+/-0.0207, T(1215)=13.21, p<2.2e-16, four cis-eQTLs (representing four eGenes) removed as extreme outliers). **B)** We found that 997 cis-eQTLs identified in liver tissue (representing 915 eGenes) were at least nominally detectable in the hippocampus, with some showing a similar slope as shown with a scatterplot (y-axis: cis-eQTL slope in liver tissue, x-axis: cis-eQTL slope in the hippocampus) (without outlier removal: Spearman’s rho=0.306, with outlier removal: R=0.297, R^2^=0.088, beta+/-SE=0.223+/-0.0228, T(990)=9.78, p<2.2e-16, five cis-eQTLs (representing three eGenes) removed as extreme outliers).

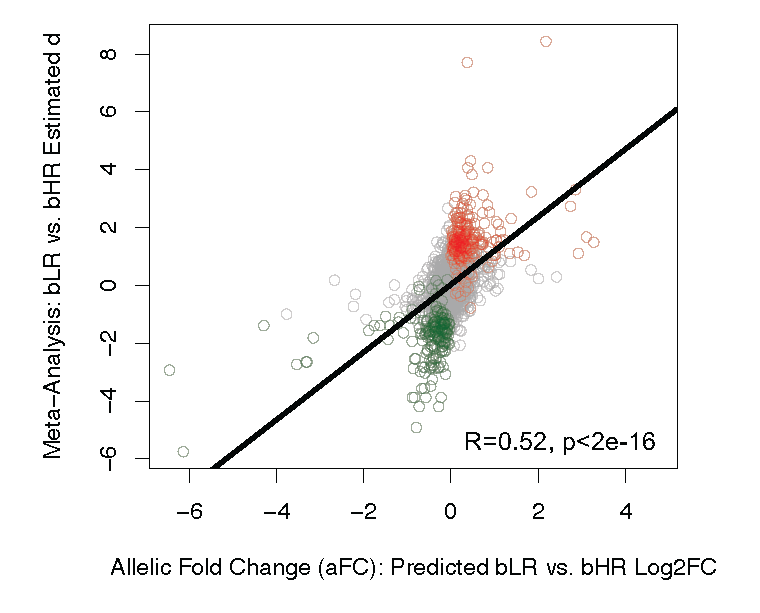

Fig S 9. Predicting the effect of bHR/bLR segregating genetic variation on gene expression.

We used our cis-eQTL database to predict the effect of genetic variation that segregates the bHR/bLR lines on gene expression. We found that 2,452 of the eVariants showed at least partial bHR/bLR segregation in the F_0_ rats (n=10 bHR/n=10 bLR sequenced in ^3^), such that if all subjects from one phenotype (e.g., bHRs) had 2 reference alleles (0/0), all subjects from the other phenotype had at least 1 alternate allele (0/1; population segregation statistic G_st’_>0.27, ^21^). To predict the effect of these bHR/bLR segregated eVariants on gene expression, we calculated the allelic Log2FC (aFC) for each eVariant and assigned the direction of effect based on the allele frequency within the bLR vs. bHR F_0_ rats. We found that these predictions correlated strongly with our F_0_ differential expression results (see main text **Fig 8B**) and with effect sizes within our previous bHR/bLR late generation meta-analysis as demonstrated in the above scatterplot (2,114 eGene/eVariant combinations: y-axis: bLR vs. bHR estimated d from the late generation meta-analysis, x-axis: Log2 allelic fold change (aFC), with direction of effect set as bLR vs. bHR based on the distribution of alleles in the F_0_s) (R=0.52, rho=0.61, beta+/SE=1.16+/-0.041, T(2112)=28.21, p<2e-16).

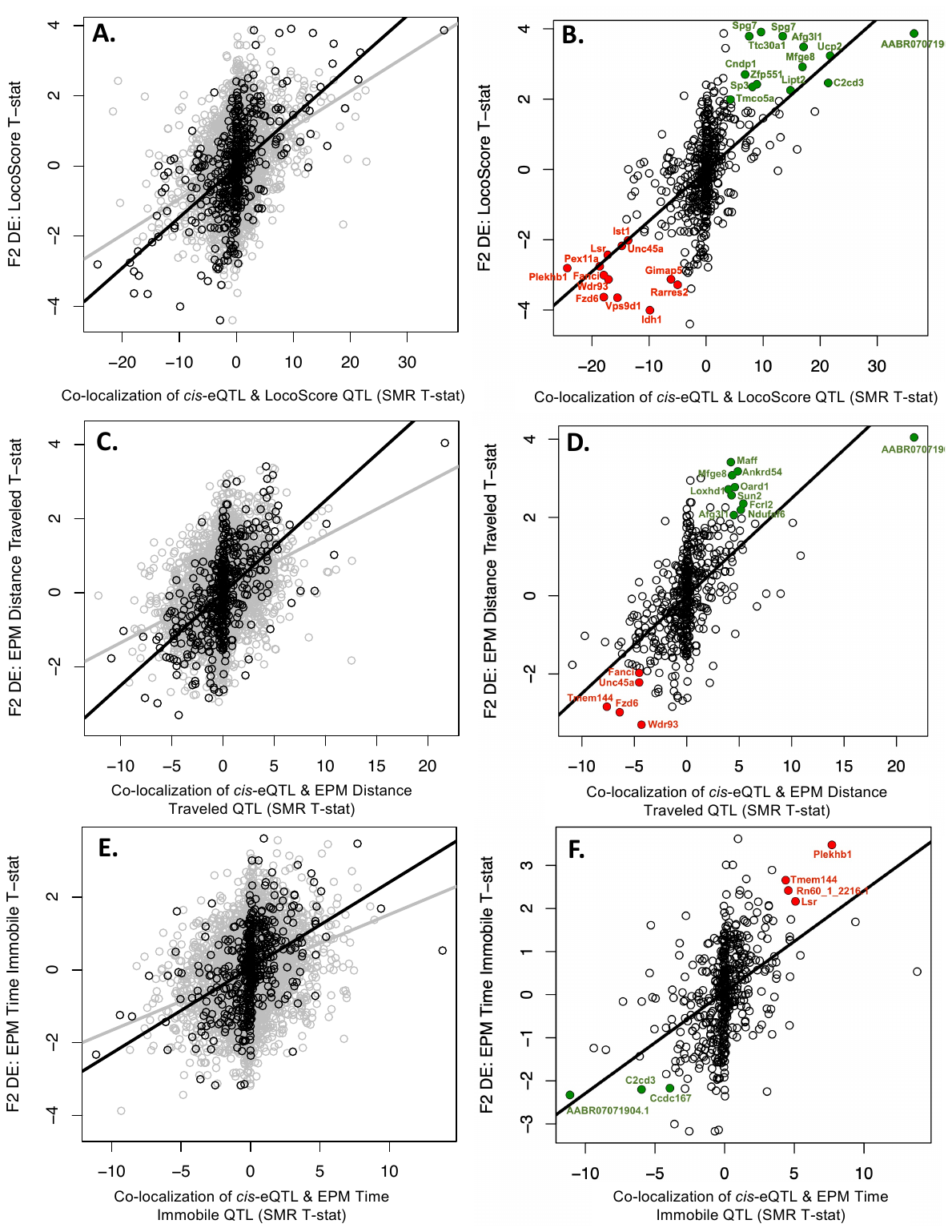

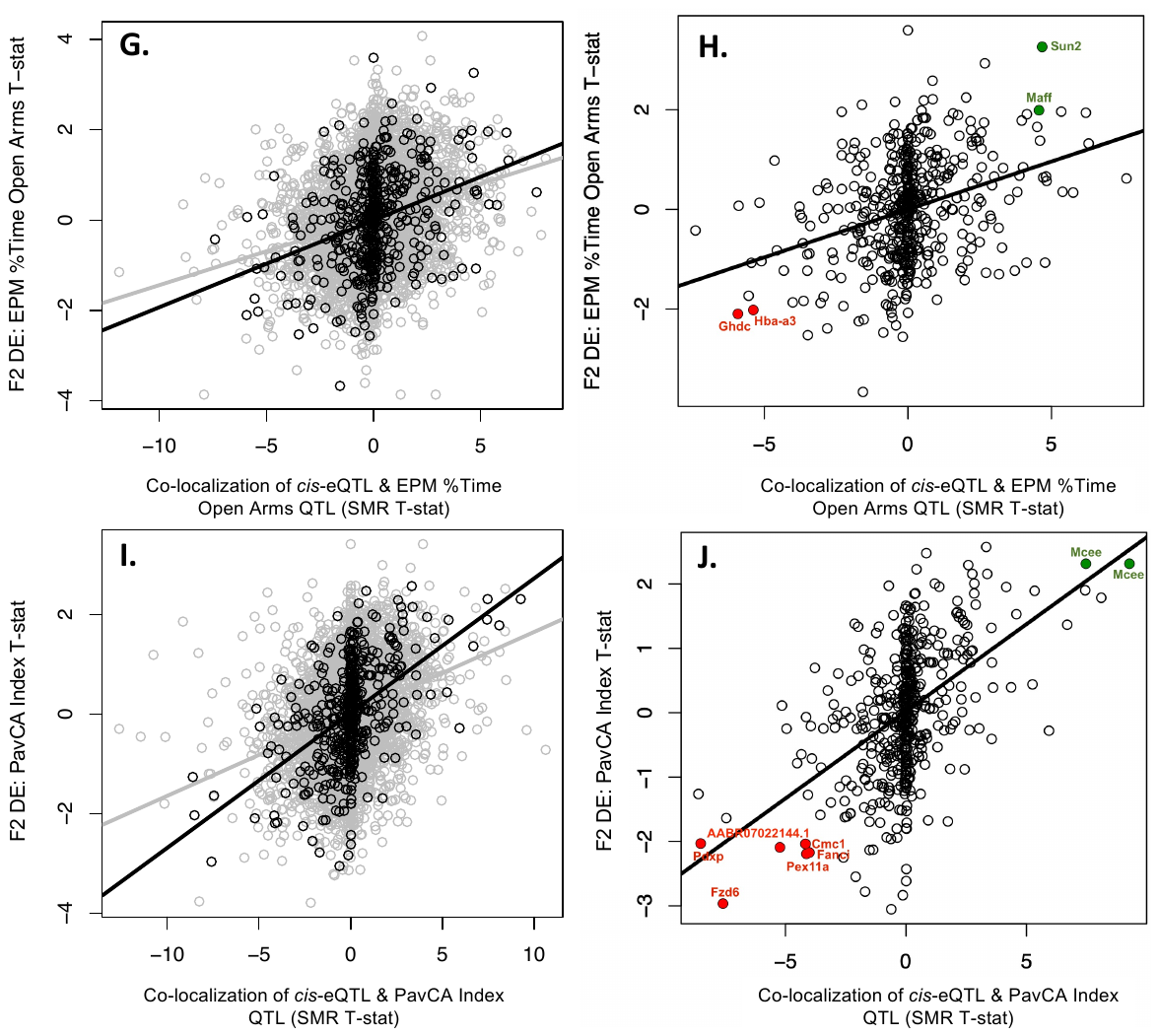

Fig S 10. Some F_2_ differential expression (DE) related to behavior can be predicted by the co-localization of cis-eQTLs with quantitative trait loci (QTLs) for the behavior identified within the larger F_2_ adult sample.

This was particularly true when considering the subset of cis-eQTLs that we had confirmed had differential expression in bHR/bLRs matching what would be expected based on the segregated distribution of alleles in the two lines (n=492 cis-eQTLs representing 456 eGenes), but also weakly true within the full sample of cis-eQTLs. The strength of the co-localization of hippocampal cis-eQTLs with regions of the genome associated with bHR/bLR-like behavior (QTLs) within the larger F_2_ sample (adults: n=323, juveniles: n=216,^3^) was determined using Summary Data-based Mendelian Randomization (SMR), and the direction of effect for the relationship between gene expression and behavior was predicted as described in **Fig 10A.** This figure contains scatterplots illustrating the positive correlation between the differential expression for each bHR/bLR-like behavior in the F_2_ sample (n=245 adults, y-axis: Log2FC) and the strength of the co-localization of cis-eQTLs with the QTLs for that behavior identified in the larger F_2_ sample (n=323 adults, x-axis: SMR T-statistic, with negative values indicating a strong predicted negative relationship between gene expression and the behavior and positive values indicating a strong positive relationship between gene expression and the behavior). Each row in the figure (e.g., **AB, CD, EF, GH, IJ**) contains the scatterplots for a particular behavior. In the left column of the figure (panels **A, C, E, G, I**), grey is used to indicate the relationship within the full 5,747 cis-eQTLs (eGene/eVariant combinations), while black is used to indicate the relationship within the subset of cis-eQTLs that we had confirmed had differential expression in bHR/bLRs matching what would be expected based on the segregated distribution of alleles in the two lines (n=492 cis-eQTLs). The right column of the figure (panels **B, D, F, H, J**) focuses on just the subset of cis-eQTLs that we had confirmed had differential expression in bHR/bLRs matching what would be expected based on the segregated distribution of alleles in the two lines (n=492 cis-eQTLs, as is depicted in black in the left column), but with colored labeling indicating genes that had nominal differential expression in the F_2_s for the behavior (p<0.05) that matched what would be predicted based on the co-localization between their cis-eQTL and the QTL for the behavior in the larger F_2_ sample (p<0.05), with green indicating upregulation with “bHR-like” behavior and red indicating upregulation with “bLR-like” behavior. **A**) A scatterplot shows the positive correlation between the differential expression for LocoScore (y-axis: Log2FC) and the strength of the co-localization of cis-eQTLs with the QTLs for LocoScore identified in the larger F_2_ sample (n=323 adults, x-axis: directional SMR T-statistic) (grey (full 5,747 cis-eQTLs): R=0.31, R^2^=0.099, beta+/SE=0.10+/-0.0041, T(5745)=25.11, p<2.2e-16), black (492 cis-eQTLs): R=0.56, R^2^=0.32, beta+/SE=0.144+/-0.00953, T(490)=15.09, p<2.2e-16). **B)** Colored labeling indicates genes that had nominal differential expression in the F_2_s for LocoScore (p<0.05) that matched what would be predicted based on the co-localization between their cis-eQTL and the QTL for LocoScore in the larger F_2_ sample (p<0.05), with green indicating “bHR-like” upregulation with increased LocoScore and red indicating “bLR-like” upregulation with decreased LocoScore. **C)** A scatterplot shows the positive correlation between the differential expression for EPM distance Traveled (y-axis: Log2FC) and the strength of the co-localization of cis-eQTLs with the QTLs for EPM Distance Traveled identified in the larger F_2_ sample (n=323 adults, x-axis: directional SMR T-statistic). (grey (full ,5747 cis-eQTLs): R=0.30, R^2^=0.092, beta+/-SE=0.14+/-0.0060, T(5745)=24.18, p<2.2e-16; black (492 cis-eQTLs): R=0.57, R^2^=0.32, beta+/-SE=0.249+/-0.163, T(490)=15.29, p<2.2e-16). **D)** Colored labeling indicates genes that had nominal differential expression in the F_2_s for EPM Distance Traveled (p<0.05) that matched what would be predicted based on the co-localization between their cis-eQTL and the QTL for EPM Distance Traveled in the larger F_2_ sample (p<0.05), with green indicating “bHR-like” upregulation with increased EPM Distance Traveled and red indicating “bLR-like” upregulation with decreased EPM Distance Traveled. **E)** A scatterplot shows the positive correlation between the differential expression for EPM Time Immobile (y-axis: Log2FC) and the strength of the co-localization of cis-eQTLs with the QTLs for EPM Time Immobile identified in the larger F_2_ sample (n=323 adults, x-axis: directional SMR T-statistic) (grey (full 5,747 cis-eQTLs): R=0.26, R^2^=0.070, beta+/-SE=0.159+/-0.0077, T(5745)=20.823, p<2.2e-16; black (492 cis-eQTLs): R=0.44, R^2^=0.19, beta+/-SE=0.235+/-0.0217, T(490)=10.85, p<2.2e-16). **F)** Colored labeling indicates genes that had nominal differential expression in the F_2_s for EPM Time Immobile (p<0.05) that matched what would be predicted based on the co-localization between their cis-eQTL and the QTL for EPM Time Immobile in the larger F_2_ sample (p<0.05), with green indicating “bHR-like” upregulation with decreased EPM Time Immobile and red indicating “bLR-like” upregulation with increased EPM Time Immobile. **G)** A scatterplot shows the positive correlation between the differential expression for EPM % Time in the Open Arms (y-axis: Log2FC) and the strength of the co-localization of cis-eQTLs with the QTLs for EPM % Time in the Open Arms identified in the larger F_2_ sample (n=323 adults, x-axis: directional SMR T-statistic) (grey (full 5,747 cis-eQTLs): R=0.23, R^2^=0.054, beta+/-SE=0.150+/-0.0082, T(5745)=18.165, p<2.2e-16; black (492 cis-eQTLs): R=0.33, R^2^=0.11, beta+/-SE=0.192+/-0.0246, T(490)=7.829, p=3.07e-14). **H)** Colored labeling indicates genes that had nominal differential expression in the F_2_s for EPM %Time in the Open Arms (p<0.05) that matched what would be predicted based on the co-localization between their cis-eQTL and the QTL for EPM %Time in the Open Arms in the larger F_2_ sample (p<0.05), with green indicating “bHR-like” upregulation with increased EPM % Time in the Open Arms and red indicating “bLR-like” upregulation with decreased EPM % Time in the Open Arms. **I)** A scatterplot shows the positive correlation between the differential expression for PavCA Index (y-axis: Log2FC) and the strength of the co-localization of cis-eQTLs with the QTLs for PavCA Index identified in the larger F_2_ sample (n=323 adults, x-axis: directional SMR T-statistic)(grey (full 5 747 cis-eQTLs): R=0.28, R^2^=0.078, beta+/-SE=0.165+/-0.00749, T(5745)=22.01, p<2.2e-16; black (492 cis-eQTLs): R=0.49, R^2^=0.24, beta+/-SE=0.271+/-0.0216, T(490)=12.55, p<2.2e-16). **J)** Colored labeling indicates genes that had nominal differential expression in the F_2_s for PavCA Index (p<0.05) that matched what would be predicted based on the co-localization between their cis-eQTL and the QTL for PavCA Index in the larger F_2_ sample (p<0.05), with green indicating “bHR-like” upregulation with increased PavCA Index and red indicating “bLR-like” upregulation with decreased PavCA Index.

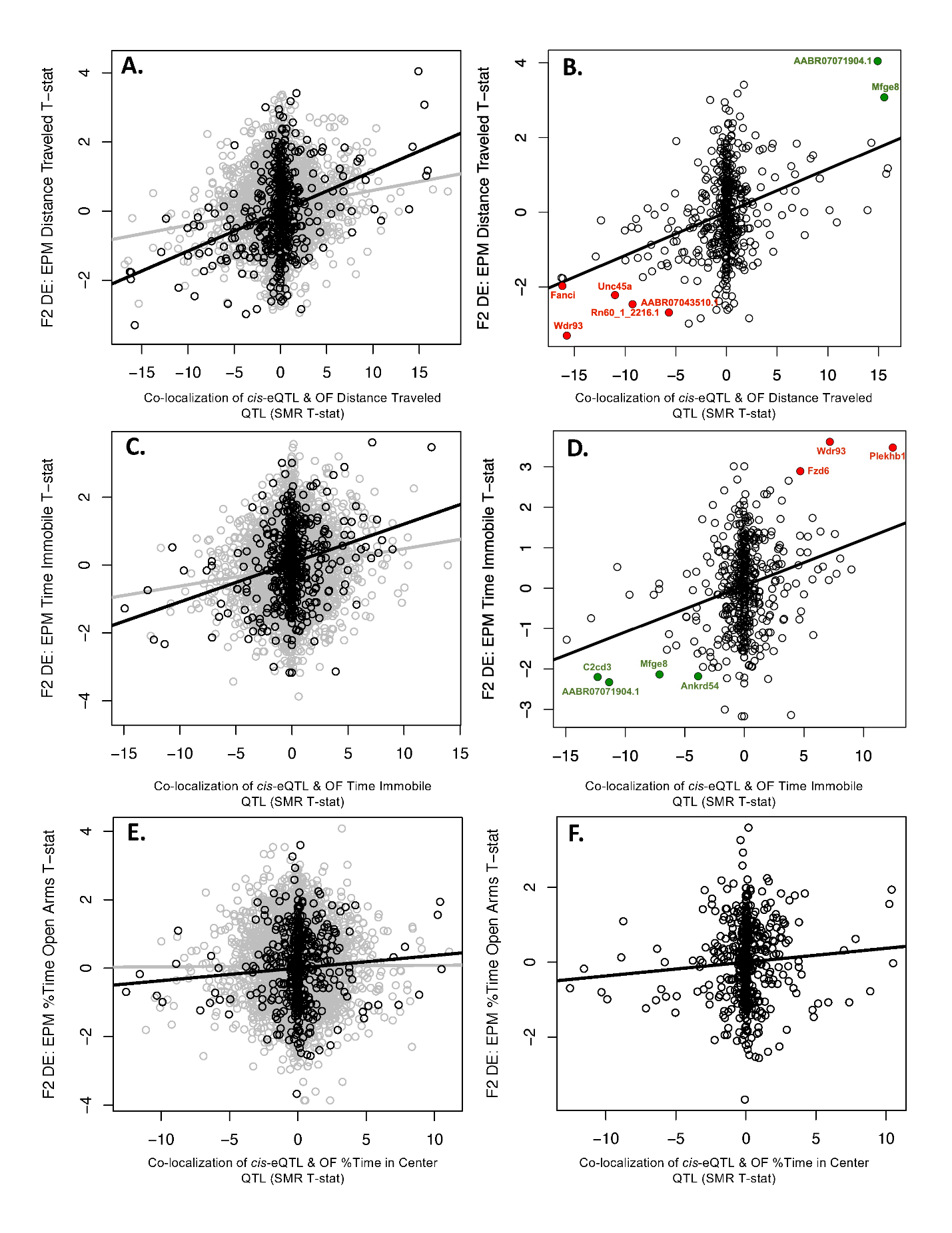

Fig S 11. Some F_2_ differential expression related to behavior can be predicted by the co-localization of cis-eQTLs with quantitative trait loci (QTLs) for analogous behaviors identified within an independent sample of F_2_ juveniles.

This figure follows the format of **Fig S10 but** contains scatterplots illustrating the positive correlation between the differential expression for bHR/bLR-like behaviors in the F_2_ adults (y-axis: Log2FC) and the strength of the co-localization of cis-eQTLs with QTLs for analogous bHR/bLR- like behaviors identified in an independent sample of F_2_ rats (n=216 juveniles, x-axis: directional SMR T-statistic). **A)** A scatterplot shows the positive correlation between the differential expression for EPM distance Traveled in the F_2_ adults (y-axis: Log2FC) and the strength of the co-localization of cis-eQTLs with the QTLs for OF Distance Traveled identified in the F_2_ juveniles (n=216, x-axis: directional SMR T-statistic) (grey (full 5,746 cis-eQTLs): R=0.123, R^2^=0.0151, beta+/-SE=0.0504+/-0.00536, T(5744)=9.398, p<2e-16; black (492 cis-eQTLs): R=0.35, R^2^=0.122, beta+/-SE=0.115, T(490)=8.249, p=1.48e-15). **B)** Colored labeling indicates genes that had nominal differential expression in the F_2_s for EPM Distance Traveled (p<0.05) that matched what would be predicted based on the co-localization between their cis-eQTLs and the QTLs for OF Distance Traveled in the F_2_ juveniles (p<0.05), with green indicating “bHR-like” upregulation with increased Distance Traveled and red indicating “bLR-like” upregulation with decreased Distance Traveled. **C)** A scatterplot shows the positive correlation between the differential expression for EPM Time Immobile in the F_2_ adults (y-axis: Log2FC) and the strength of the co-localization of cis-eQTLs with the QTLs for OF Time Immobile identified in the F_2_ juveniles (n=216, x-axis: directional SMR T-statistic) (grey (full 5,746 cis-eQTLs): R=0.102, R^2^=0.0104, beta+/-SE=0.0551+/-0.0071, T(5744)=7.76, p=9.56e-15; black (492 cis-eQTLs): R=0.26, R^2^=0.0679, beta+/-SE=0.115+/-0.0192, T(490)=5.975, p=4.44e-09). **D)** Colored labeling indicates genes that had nominal differential expression in the F_2_s for EPM Time Immobile (p<0.05) that matched what would be predicted based on the co-localization between their cis-eQTLs and the QTLs for OF Time Immobile in the juveniles (p<0.05), with green indicating “bHR-like” upregulation with decreased Time Immobile and red indicating “bLR-like” upregulation with increased Time Immobile. **E)** A scatterplot shows the lack of positive correlation between the differential expression for EPM % Time in the Open Arms in the F_2_ adults (y-axis: Log2FC) and the strength of the co-localization of cis-eQTLs with the QTLs for OF % Time in the Center identified in the F_2_ juveniles (n=216, y-axis: SMR T-statistic) (grey (full 5,746 cis-eQTLs): R=0.0038, R^2^= 1.453e-05, beta+/-SE= 0.00228+/-0.00789, T(5744)=0.289; black (492 cis-eQTLs): R=0.0757, R^2^=0.00574, beta+/-SE=0.0369+/-0.0219, T(490)=1.682). **F)** The lack of colored labeling indicates that no genes had nominal differential expression in the F_2_s for EPM % Time in the Open Arms (p<0.05) that matched what would be predicted based on the co-localization between their cis-eQTLs and the QTLs for OF % Time in the Center in the F_2_ juveniles (p<0.05).

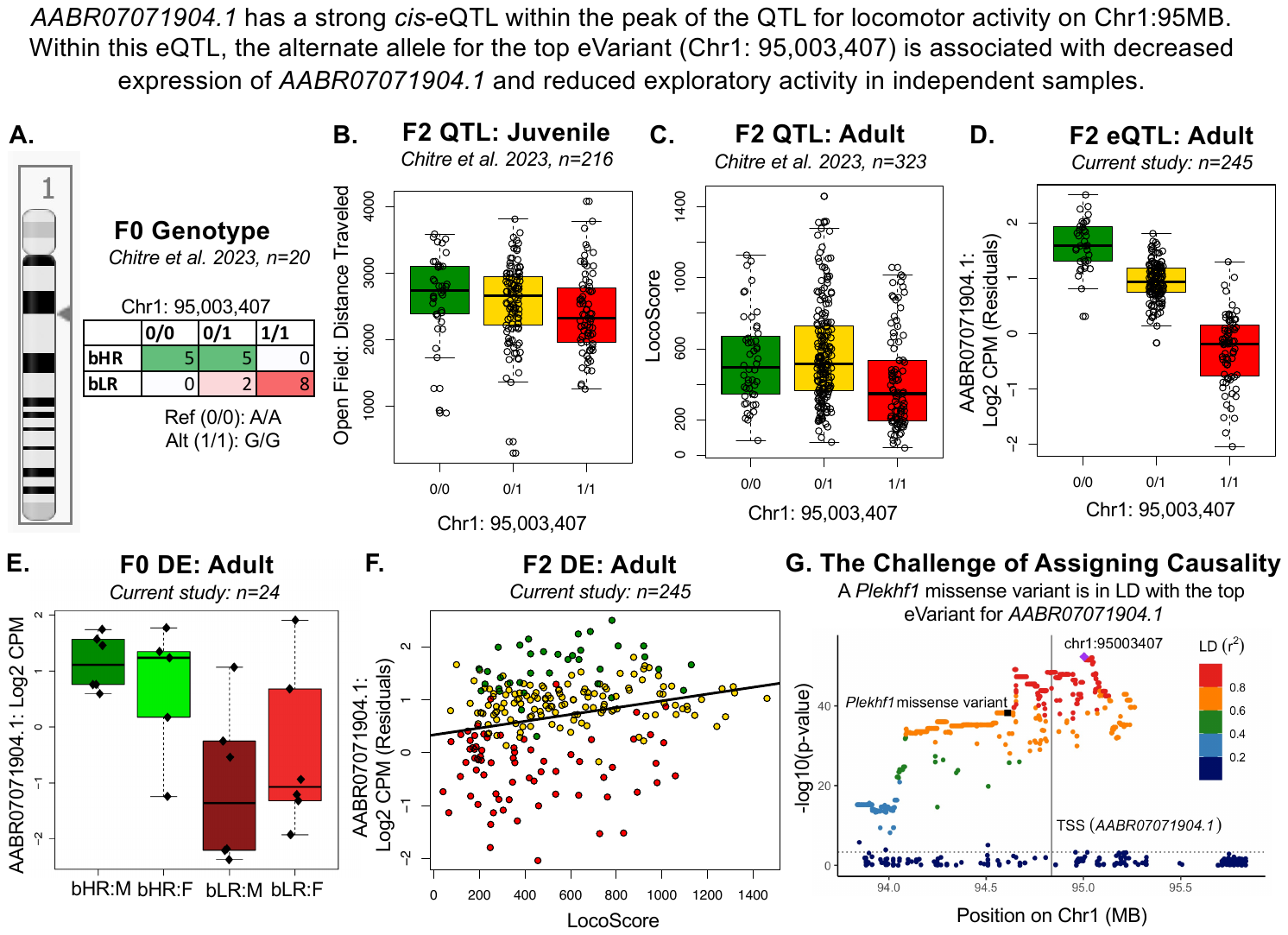

Fig S 12. Example top candidate gene: Evidence supporting the potential for AABR07071904.1 to mediate the relationship between genetic variation and behavior.

AABR07071904.1 has a strong cis-eQTL within the peak of the QTL for locomotor activity on Chr1:95MB. The provided plots illustrate the relationship of the top eVariant (Chr1: 95,003,407) within this cis-eQTL with bHR/bLR phenotype and locomotor activity, as well as the relationship with the expression of AABR07071904.1 itself. Within all plots, green vs. red color is used to indicate either bHR vs. bLR phenotype or the allele overrepresented in each respective phenotype. Gold is used to indicate heterozygotes (0/1). **A)** The alternate allele (G/G) for the top eVariant is more prevalent in bLRs, whereas bHRs are more likely to carry the reference allele (A/A) ^3^. **B)** The alternate allele (G/G) for the top eVariant is associated with decreased distance traveled in the open field in F_2_ juveniles (n=216; ^3^). **C)** The alternate allele (G/G) for the top eVariant is associated with decreased LocoScore in the full sample of F_2_ adults (n=323; ^3^). **D)** The alternate allele (G/G) for the top eVariant is associated with decreased expression of AABR07071904.1. To illustrate this relationship, gene expression (Log2 CPM) is plotted as residual expression after quality control and controlling for the technical covariates included in our differential expression model (n=245), similar to the eQTL analysis. **E)** AABR07071904.1 is more highly expressed (y-axis: Log2 CPM) in the hippocampus of male (M) and female (F) bHR rats than male (M) and female (F) bLR rats in our F_0_ sample. **F)** AABR07071904.1 is more highly expressed in the hippocampus of F_2_ rats with a higher LocoScore. Similar to the eQTL plot, gene expression (Log2 CPM) is plotted as residual expression after quality control and controlling for the technical covariates included in our differential expression model (n=245). Note that the trendline is an approximation illustrating a simple linear regression of the presented data, and not the final results of the more sophisticated differential expression analysis in our paper. **G)** The top eVariant for AABR07071904.1 is in linkage disequilibrium (LD) with the Plekhf1 missense variant identified as a top candidate in ^3^, illustrating the challenges of assigning causality. Plekhf1 is not differentially expressed in bHR/bLR or in the F_2_ intercross sample. To illustrate this LD, the –log(10)p-values are plotted from the cis-eQTL analysis for the relationship between AABR07071904.1 expression and all SNPs tested within the cis-window (+/-1 Mb of the transcription start site or TSS). The purple diamond indicates chr1:95,003,407, the cis-eQTL top variant. The black square indicates the Plekhf1 missense variant of interest. All remaining points are colored according to their LD (r^2^) with the top variant chr1:95,003,407 (red=greater LD, blue=less LD). The dashed horizontal line shows the gene-specific significance threshold for the cis-eQTL analysis.

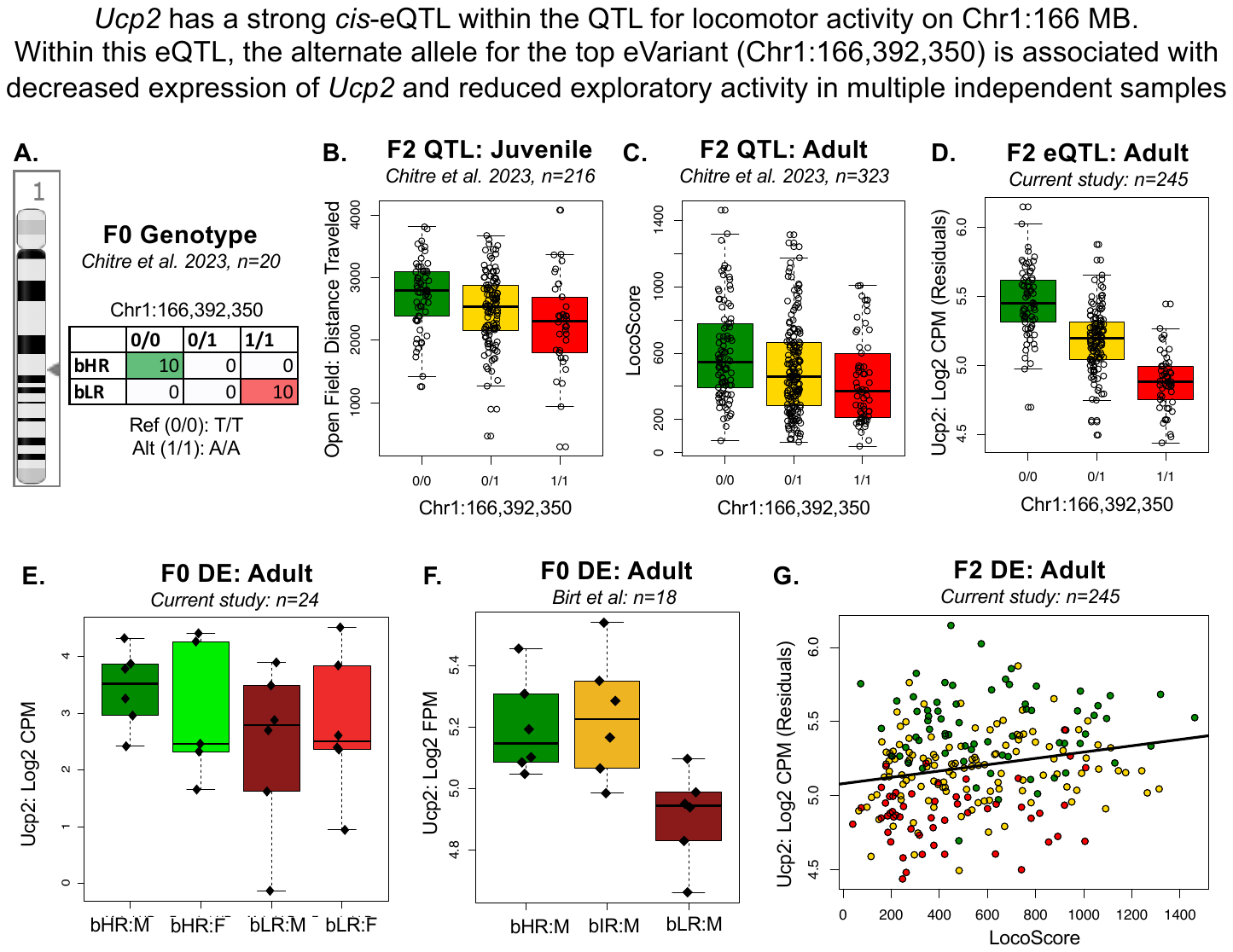

Fig S 13. Example top candidate gene: Evidence supporting the potential for Ucp2 to mediate the relationship between genetic variation and behavior.

Ucp2 has a strong cis-eQTL within a QTL for locomotor activity on chromosome (chr) 1. The provided plots illustrate the relationship of the top eVariant (Chr1: 166,392,350) within this cis-eQTL with bHR/bLR phenotype and locomotor activity, as well as the relationship with the expression of Ucp2 itself. Within all plots, green vs. red color is used to indicate either bHR vs. bLR phenotype or the allele overrepresented in each respective phenotype. Gold is used to indicate heterozygotes (0/1). A) The alternate allele (A/A) for the top eVariant is more prevalent in bLRs, whereas bHRs are more likely to carry the reference allele (T/T) ^3^. B) The alternate allele (A/A) for the top eVariant is associated with decreased distance traveled in the open field in F_2_ juveniles (n=216; ^3^). C) The alternate allele (A/A) for the top eVariant is associated with decreased LocoScore in the full sample of F_2_ adults (n=323; ^3^). D) The alternate allele (A/A) for the top eVariant is associated with decreased expression of Ucp2. To illustrate this relationship, gene expression (Log2 CPM) is plotted as residual expression after quality control and controlling for the technical covariates included in our differential expression model (n=245), similar to the cis-eQTL analysis. E) Ucp2 is more highly expressed (y-axis: Log2 CPM) in the hippocampus of male (M) and female (F) bHR rats than male (M) and female (F) bLR rats in our F_0_ sample. E) Ucp2 is more highly expressed (y-axis: Log2 FPM) in the hippocampus of male bHR rats than male bLR rats in our previous F_0_ sample ^4^. Cross-bred intermediate responders (F_1_ bIRs) resemble bHRs. G) Ucp2 is more highly expressed in the hippocampus of F_2_ rats with a higher LocoScore. Similar to the eQTL plot, gene expression (Log2 CPM) is plotted as residual expression after quality control and controlling for the technical covariates included in our differential expression model (n=245). Note that the trendline is an approximation illustrating a simple linear regression of the presented data, and not the final results of the more sophisticated differential expression analysis in our paper.

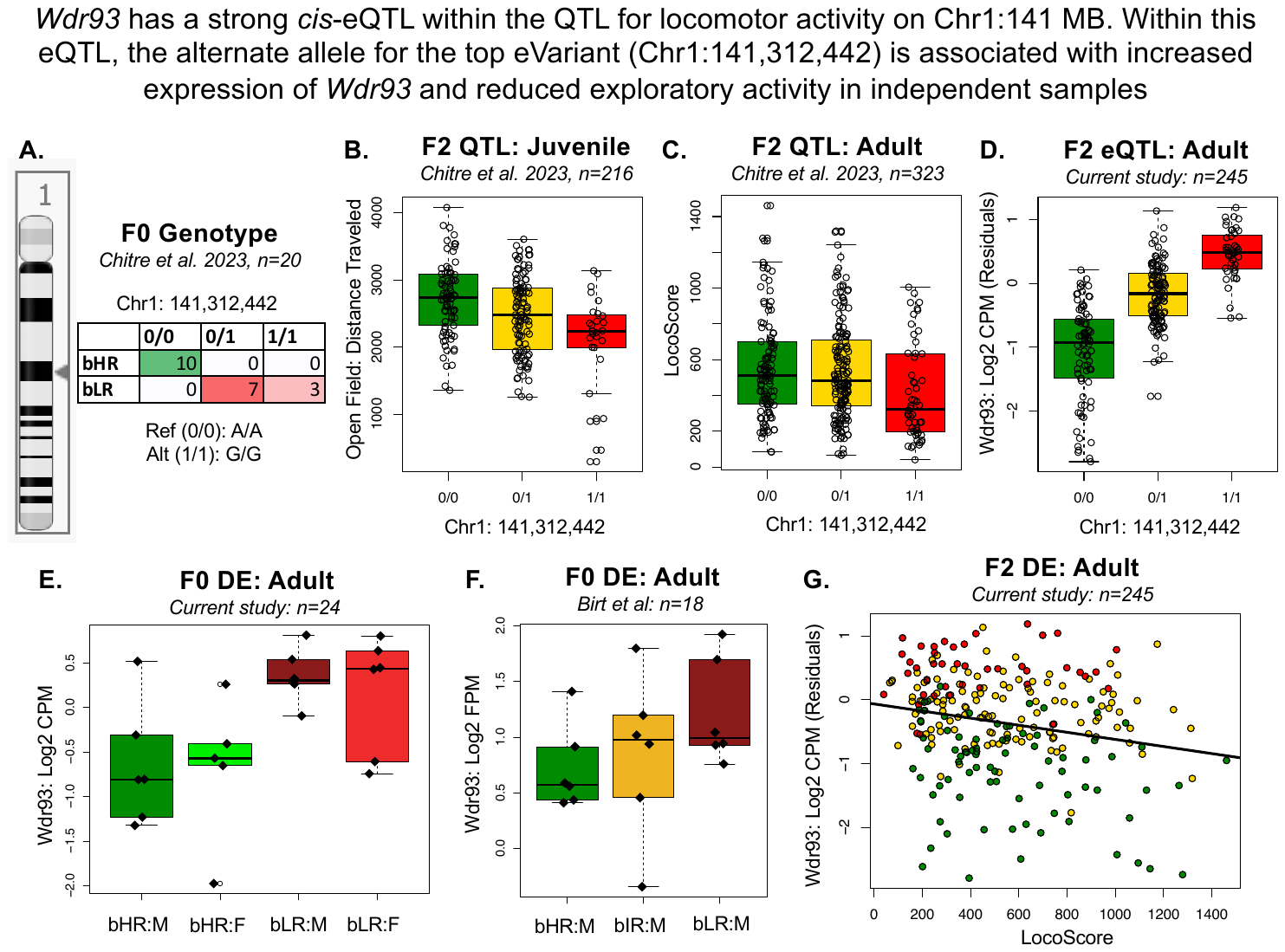

Fig S 14. Example top candidate gene: Evidence supporting the potential for Wdr93 to mediate the relationship between genetic variation and behavior.

This figure follows the format of Fig S12-FigS13.

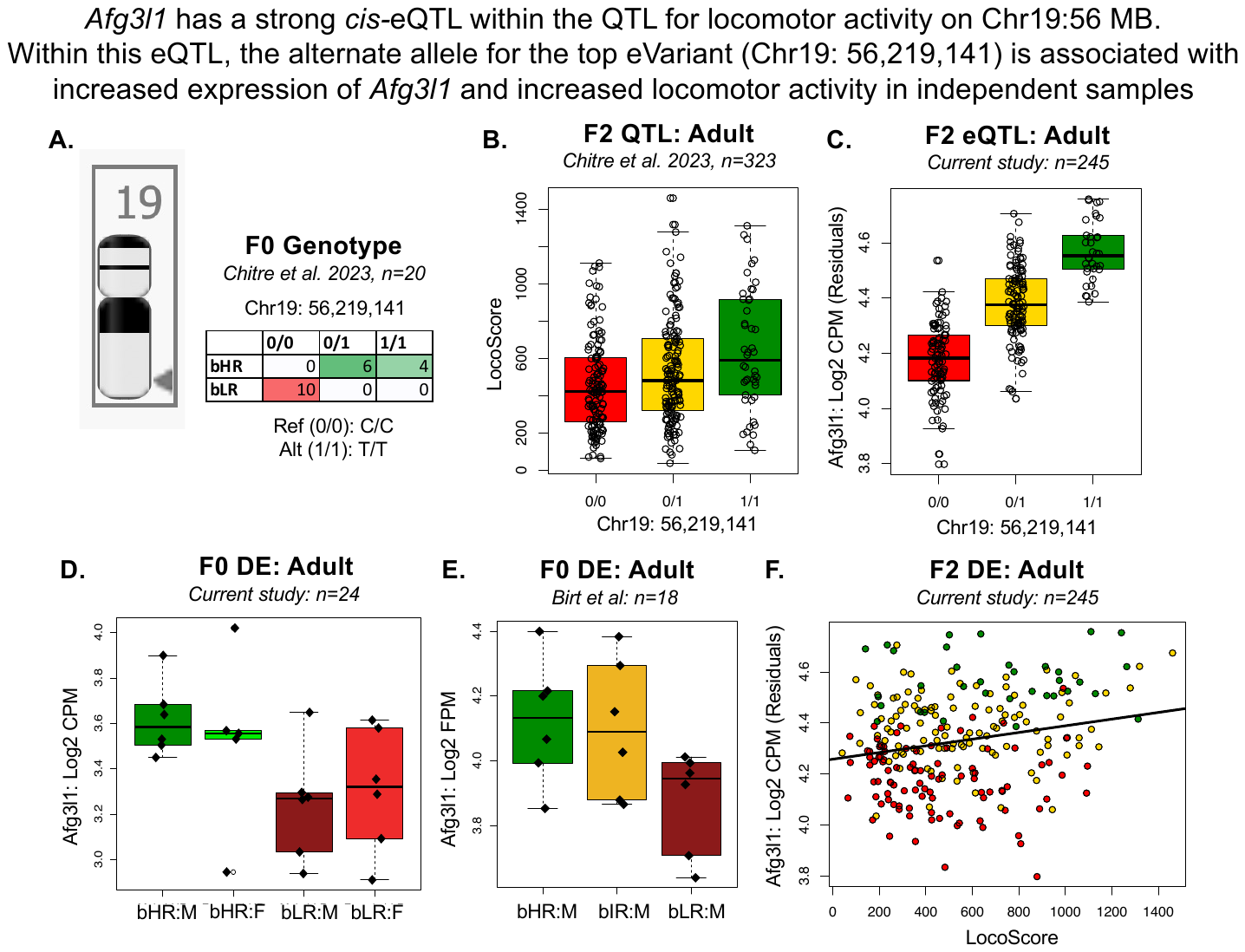

Fig S 15. Example top candidate gene: Evidence supporting the potential for Afg3l1 to mediate the relationship between genetic variation and behavior.

This figure follows the format of Fig S12-FigS13 but does not overlap a F_2_ juvenile QTL.

### Appendices

Appendix 1. Materials Design Analysis Reporting (MDAR) Checklist for Authors

Appendix 2. The ARRIVE guidelines 2.0: author checklist. The ARRIVE Essential 10.
